# Observing biological spatio-angular structures and dynamics with statistical image reconstruction and polarized fluorescence microscopy

**DOI:** 10.1101/2025.09.23.678164

**Authors:** Junyu Liu, Talon Chandler, Yue Li, Atharva Agashe, Mingzhe Wei, Yijun Su, Yicong Wu, Tobias I. Baskin, Valentin Jaumouillé, Jiji Chen, Pengcheng Xu, Huihui Ye, Wentao Zhu, Robert S. Fischer, Vinay Swaminathan, Amrinder S. Nain, Shalin B. Mehta, Patrick J. La Riviere, Hari Shroff, Huafeng Liu, Min Guo

## Abstract

Understanding molecular orientation and density distributions is essential for exploring biological structure and function. Polarized fluorescence microscopy (PFM) provides insights into molecular architecture but struggles to resolve three-dimensional (3D) molecular orientation distributions, particularly in densely labeled or structurally complex specimens. To address this, we introduce the efficient generalized Richardson-Lucy (eGRL) algorithm, a robust framework for reconstructing 3D molecular density and orientation (spatio-angular) distributions from PFM data. By modeling the imaging process in spatio-angular hyperspace, we propose a maximum-likelihood solution enhanced by dimensionality reduction and angular domain transformation to overcome computational challenges. eGRL improves accuracy and efficiency across different PFM implementations, enabling use on standard platforms. We utilize our methods to resolve biological spatio-angular structures and dynamics otherwise impossible to resolve, including the tangential alignment of actin filaments in U2OS cells, nanowire-guided cytoskeletal organization in NIH3T3 cells, rotational actin patterns in live HeLa protrusions, and membrane tension-induced anisotropy in live macrophages.

## Introduction

Most fluorescent probes tethered to biological structures display polarization-dependent characteristics that are intricately linked to the physicochemical attributes of their environment. These characteristics, determined by fluorophore orientation, can elucidate molecular function, yet are often neglected in conventional microscopy. Recent attempts to map fluorophore orientations have begun to yield biological insights, revealing heterogeneities between amyloid fibrils^1,2^ or lipid membranes^3^, periodic orientations of intertwined supercoiled DNA^4^, and stress-sensitive orientational distributions along actin filaments^5^.

To facilitate investigation in densely labeled specimens with complex and/or highly dynamic structures, rapid acquisition and extraction of biological insights at the ensemble scale are essential. Many efforts estimate the average lateral orientation of fluorophores within diffraction-limited volumes by measuring polarization modulation (PM)^6–14^. These advances have revealed the 90° rotation of hourglass septin filaments oriented along the budding neck in yeast^8^, mapped the orientation of nuclear pore proteins^15^, and, when combined with structured illumination microscopy (pSIM) enabled investigation of the orientational dynamics of green fluorescent protein-labeled microtubules in live U2OS cells^16^. Most PM studies use insights gained from probing the average two-dimensional (2D) orientation of fluorescent samples. The corresponding 3D spatio-angular distributions have been much less explored.

Addressing this gap, we previously introduced a theoretical framework for spatio-angular fluorescence imaging^17^. This framework promised to obtain both the 3D density and orientation distributions of fluorophores by capturing the fluorescence emissions under polarization modulation. In practice, this framework is hindered by the underdetermined nature of the inverse problem and the computational challenges associated with reconstructing high-dimensional spatio-angular distributions. Conventional reconstruction algorithms, such as those based on singular value decomposition (SVD), require tuning a parameter which is sensitive to noise and often introduces artifacts or biases that compromise the fidelity of orientation mapping^18,19^.

Here we introduce the efficient generalized Richardson-Lucy (eGRL) algorithm, an iterative spatio-angular image reconstruction framework tailored for polarized fluorescence microscopy (PFM). By statistically modelling the forward process of the oriented fluorophores imaging, we first derive a general maximum-likelihood expectation-maximization solution to the inverse problem, enabling the joint estimate of 3D density and orientation distributions of fluorescent molecules. Then we reduce the number of angular dimensions and modify our iterative framework to dramatically alleviate computational load without sacrificing reconstruction accuracy. Collectively, these advances enhance the accuracy and efficiency of spatio-angular image reconstruction, allowing eGRL to be deployed effectively on standard computational platforms. We validated the eGRL framework on both simulated imaging systems and our home-built polarized dual-view inverted selective-plane illumination microscope (pol-diSPIM)^18^. Compared to previous methods, eGRL better delineates biological structures and dynamics. Finally, we built a pipeline to mitigate the exponential increase in RAM and time consumption caused by hyper-dimensional computation in large time-lapse datasets, enabling further efficiency gains when performing spatio-angular deconvolution on consumer-grade graphical processing unit (GPU) cards. Using eGRL, we reveal the angular pattern of actin grown on nanowires and visualize short-timescale molecular dynamics in living cells, demonstrating the potential of spatio-angular estimation for biological investigation.

## Results

### An improved framework for spatio-angular estimation

Polarized fluorescence microscopy (PFM) has historically estimated pixelwise fluorescence anisotropy factors or underlying lateral molecular orientations^8,20–22^. These estimates are subsets of the spatio-angular distribution, which quantifies the number and orientation of fluorescence emitters at each resolved position in three-dimensions^18^ (**Fig. 1a**). Conversely, a 3D spatio-angular distribution provides direct access to the orientation distribution functions (ODFs) at each distinguishable location in 3D space and allows for the extraction of spatial density maps, peak orientation maps, and other molecular properties (**Supplementary Fig. 1, Supplementary Video 1**).

**Fig. 1.**
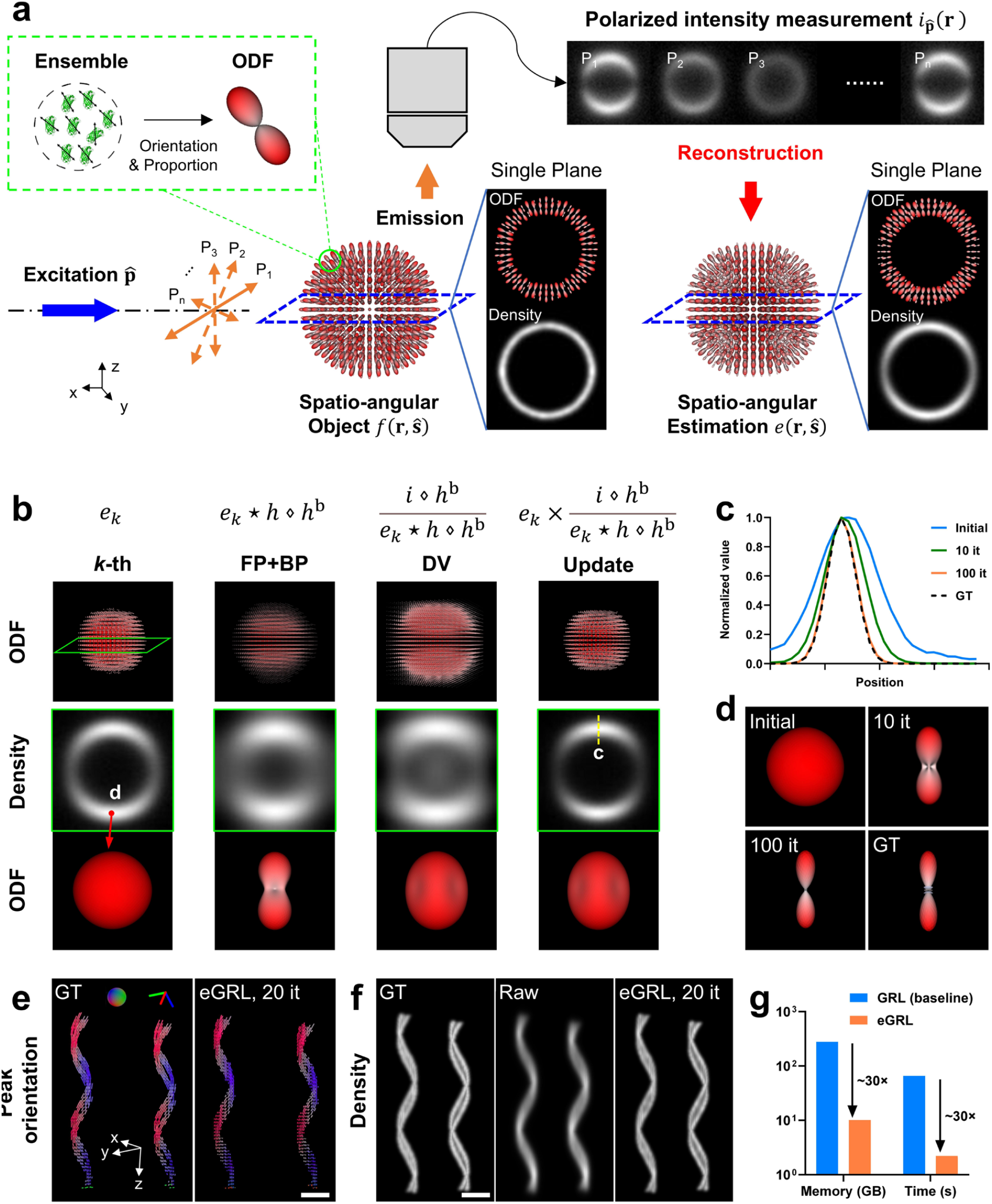
Iterative reconstruction algorithm enables efficient estimate of 3D spatio-angular distribution. **a)** General imaging and reconstruction process for polarized fluorescence microscopy. An ensemble of fluorescent dipoles in a diffraction-limited region can be represented by an orientation distribution function (ODF), which is a surface with a radius proportional to the number of dipoles in the measurement volume that are oriented along each direction. Fluorophores in the object are then described by a spatio-angular distribution, i.e. the density distribution and ODF in each voxel. With different polarized excitations P_1_∼P_n_, fluorescence emissions are collected by the objective lens and form the polarized intensity measurements on the camera, which can then be used to estimate the underlying spatio-angular distribution. **b)** Schematic design of eGRL (efficient Generalized Richardson-Lucy) reconstruction algorithm consisting of three parts in each iteration: FP+BP (forward and back projection), DV (division) and Update, simultaneously estimating density and orientation distributions. Top raw visualizes the 3D ODF map, middle row is the density map of central slice from the highlighted region in top row while the bottom row shows the ODF from the spot location in the row above. **c), d)** The profiles along the yellow dashed line and the shape of the ODF at the red pointer in **b** after increasing numbers of iterations, gradually converging to the ground truth. **e)** Spatio-angular reconstructions on synthetic double-helix phantoms. The left helices are separated by 478 nm inter-helix space, and the right helices have 554 nm inter-helix space. Each dipole moment is parallel to the tangential direction of the helix, and each helix has a uniform density but periodic peak orientations and varying ODF anisotropy increasing from bottom to top, reconstruction is performed with eGRL, 20 iterations. Peak orientation maps with pseudo-color are demonstrated. **f)** density maps of the ground truth (left), the simulated raw data (middle, average of 42 polarization measurements), and eGRL reconstruction (right) corresponding to phantoms in **e. g)** Time and memory cost of eGRL reconstruction in a dataset with 100×100×100 voxels and 42 polarization modulations (ran on single CPU thread without multithread acceleration), demonstrating the 30-fold reduction obtained by transferring the computation from GRL (baseline) to eGRL with angular dimensionality reduction. Scale bars: **e, f** 5 μm. See also **Supplementary Figs. 1-4**.

Assuming an imaging system with translationally invariant detection, the imaging process of PFM (**Fig. 1a**) can be reformulated as a spatio-angular convolution in hyperspace of (1) 3D spatial coordinates, (2) angular distributions on a sphere, and (3) polarization states. The fluorescence intensity captured on the camera is a joint projection of structural density and orientation information, with spatial blurring and angular aliasing:

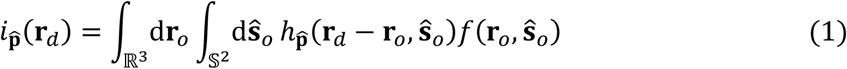

where 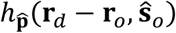 is the irradiance at position **r**_*d*_ on the camera coordinates, created by a point source under polarized excitation 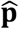 at **r**_*o*_ with orientation **ŝ**_*o*_, referred to as dipole point spread function (dipole PSF). *f*(**r**_*o*_, **ŝ**_*o*_) denotes the spatio-angular distribution of the object which we are interested in estimating and 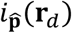 the intensity measurement under corresponding polarization modulation. ℝ^3^ and 𝕊^2^ represent the volume in 3D space and the orientation ensemble on the spherical surface, respectively.

Spatial and angular information are coupled together during the imaging process; consequently, the image reconstruction requires solving a high-dimensional inverse problem that connects the measured fluorescence intensity to the spatio-angular distribution *f*(**r**_*o*_, **ŝ**_*o*_) in the hyperspace. We use spatio-angular deconvolution to deduce the spatial distribution in the entire volume and the omnidirectional angular distribution within each diffraction-limited subvolume. This deconvolution is based on 1) the information contained in a series of polarized intensity measurements (**Fig. 1a**) and 2) system modelling via a dipole PSF (**Methods**); nevertheless, the additional polarization and orientation dimensions in the hyperspace require a new approach distinct from traditional intensity deconvolution strategies. By modeling the photon detection as a Poisson process, we extend the maximum-likelihood expectation-maximization (MLEM) theory^23^ to a high-dimensional framework accounting for the spatio-angular hyperspace (**Supplementary Note 1**), and derive an iterative update formula to reconstruct both the 3D density (spatial) and orientation (angular) distributions of fluorophores (**Methods, Supplementary Note 1**):

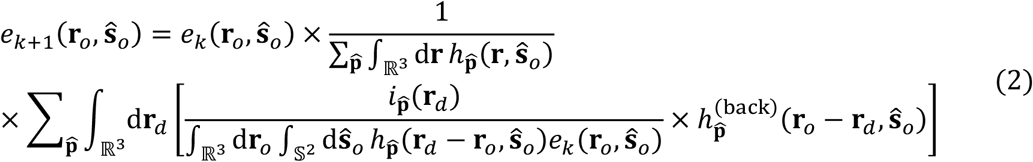

where *e*_*k*_ denotes the current estimate and *e*_*k*+1_ the updated estimate. *h*^(back)^ is the back projector which typically is the spatial transpose of *h*. With this key formulation, the resulting algorithm extends the conventional Richardson-Lucy deconvolution (RLD) to the spatio-angular hyperspace, resulting in the generalized Richardson-Lucy (GRL) algorithm.

Compared with conventional RL deconvolution, this baseline GRL algorithm contains additional polarization (usually multiple modulations) and angular dimensions (typically thousands of orientations) and thus poses higher demands on computational memory and time, practically impeding the implementation of such an iterative algorithm. To address this challenge, we further develop a mathematical optimization strategy with two changes to dramatically reduce the data dimensionality and computational load, including an angular domain transformation and a restructuring of the iterative framework^24^ (**Supplementary Note 2**), culminating in the final reconstruction framework (**Methods**), which we term the efficient generalized Richardson-Lucy (eGRL) algorithm

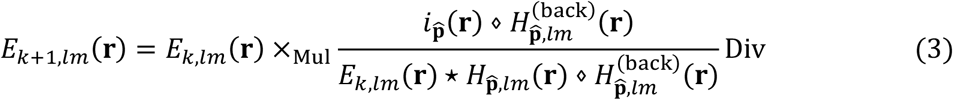

where 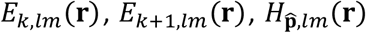 and 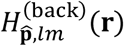 are terms corresponding to the angular spectra of 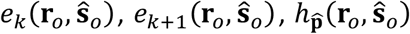 and 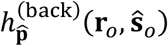, ⋆ and ⋄ denote the forward and backward projections between 3D spatio-angular object space and polarization measurement space operating in spatial-frequency and angular-spectrum domains, and the subscripts Mul and Div denote specialized multiplication and division operations in spatio-angular space (**Methods, Supplementary Note 2**). 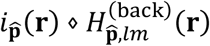 can be precomputed outside the iteration loop. The same applies to the successive forward and backward projection 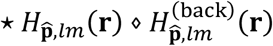, which can further be combined to one single operator independent of the polarization components, thereby reducing the computational burden during each iteration, which is especially important when using an extensive collection of polarization measurements (**Supplementary Note 2**).

### Characterizing eGRL performance with synthetic data

To illustrate the iterative progress of the algorithm, we demonstrate the intermediate outputs (**Fig. 1b**) during the first iteration of eGRL on simulated, noise-free data of a shell object with an ideal uniform density distribution and molecular orientations perpendicular to the spherical surface. The current estimate in object space is projected directly to a virtual ‘back-projected object space’ (FP+BP), divided by the back-projected raw data for a comparison in that space (DV), and finally multiplied by the current estimate to yield an updated estimate of the sample (Update). As the algorithm iterates, both the full width at half maximum (FWHM) of the shell thickness (**Fig. 1c**) and the orientation distribution (**Fig. 1d**) converge towards the ground truth (see **Supplementary Video 2** for an example of reconstruction on another phantom).

Since spatial and angular information are coupled during the imaging process, we then applied eGRL to synthetic datasets with more complex spatio-angular distributions to show how the spatial and angular structures can be resolved from the mixed intensity measurements. Here, we used double-helix phantoms, in which each double-helix contains two helices in close proximity and with periodic variations in molecular orientation (**Fig. 1e, Supplementary Fig. 2**). eGRL produces an orientation distribution matching the ground truth (GT) with high fidelity (**Fig. 1e, Supplementary Fig. 2a**), and recovers the distance between the two helices, which is difficult to discern in the raw images (**Fig. 1f, Supplementary Fig. 2b**). We also found the reconstructed spatial density distribution is equivalent to the result obtained by the traditional Richardson-Lucy (RL) deconvolution^25,26^ (**Supplementary Fig. 2b, c**), with a structural similarity index measure (SSIM) of 0.997. We further confirmed the close match between the eGRL and RL estimates of the density distributions based on fluorescent beads acquired without polarization modulation (**Supplementary Fig. 3, Methods**).

Next, we checked the computational efficiency of eGRL by monitoring the memory usage and the time cost for one iteration, comparing the baseline GRL performance, to the final application of eGRL with its angular dimensionality reduction and modified iterative framework (**Fig. 1g**). For a typical dataset (100×100×100 voxels with 42 polarization modulations), while eGRL and GRL produce similar reconstructions with similar performance in the presence of noise (**Supplementary Fig. 4**), the eGRL required 2.2 s per iteration, vs. 65.9 s for the original GRL method (without transforming from angle to angular-spectrum and reshuffling the order of RL intermediate steps), offering a ∼30-fold improvement in speed. eGRL also reduced the memory footprint from 278.0 GB to 10.1 GB, corresponding to a ∼28-fold reduction, making it feasible to deploy the eGRL algorithm on standard workstations.

Next, we assessed eGRL on synthetic datasets mimicking various optical configurations of PFM, including wide-field microscopy, light sheet microscopy (SPIM), and dual-view light sheet microscopy (diSPIM)^27,28^ (**Fig. 2, Supplementary Figs. 5, 6**). These three microscopy configurations are commonly used for 3D imaging due to their rapid imaging rate and therefore are preferential for spatio-angular data acquisition with polarization modulation (**Methods**). We simulated the imaging process, generating synthetic raw data (**Supplementary Fig. 7**) of carefully designed helix phantoms (**Fig. 2a**), and then used eGRL as well as the traditional singular-value-decomposition based algorithm (SVD)^18^ to reconstruct density and orientation distributions from the raw data. Our eGRL method reconstructs density and orientation distributions of 3D objects with superior performance compared to SVD for all three microscopy configurations (**Fig. 2b**). By examining the density and peak orientation maps from each microscopy configuration (**Fig. 2c, d**), we found that eGRL produces density structures that are sharper than the SVD results, and produces orientation distributions closer to the ground truth, where the orientational rotation along helices can be easily observed.

**Fig. 2.**
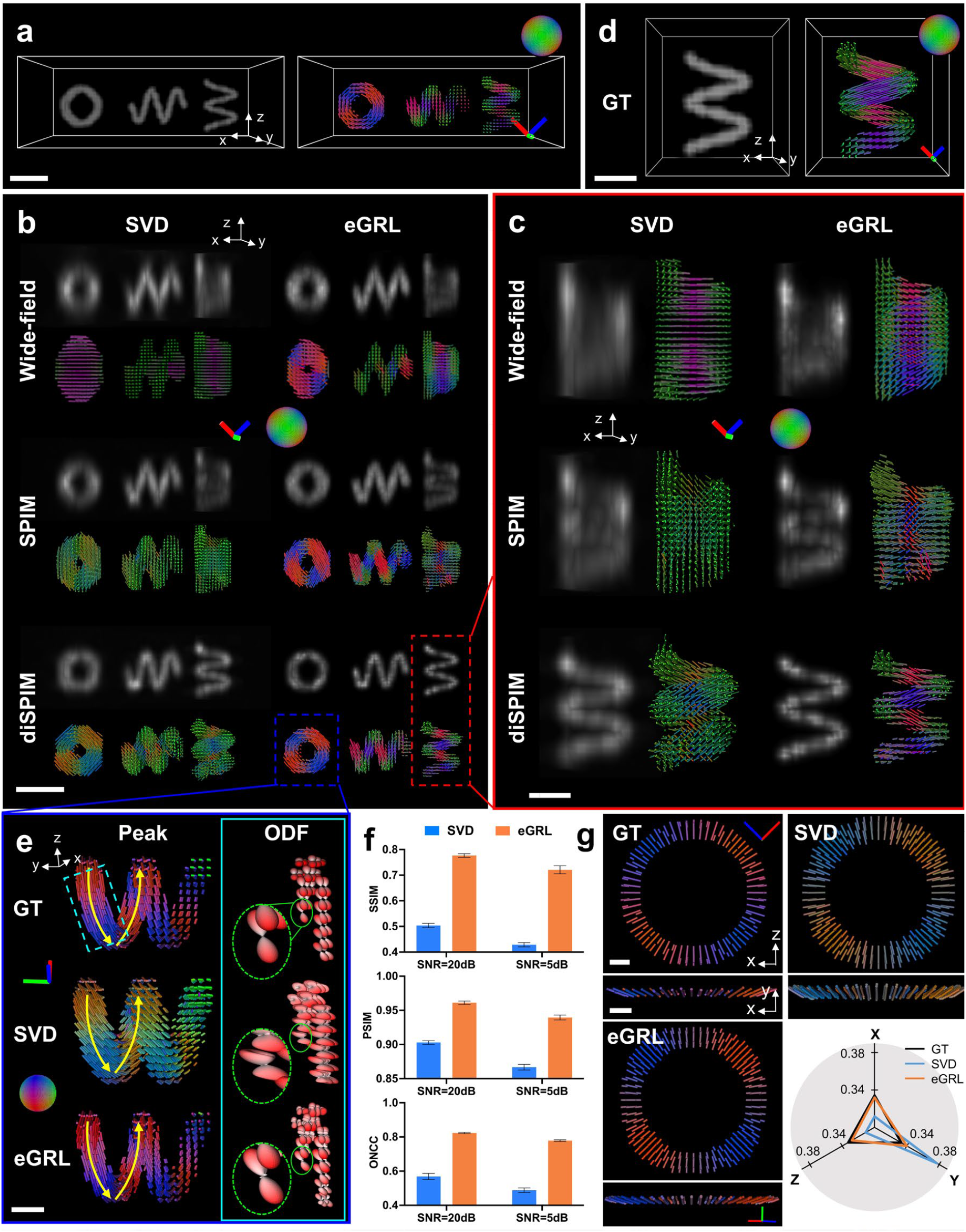
Spatio-angular image reconstruction from simulated data of different polarized fluorescence microscopes. **a)** Density (left) and peak orientation (right) distributions of three synthetic helices, each oriented along a different axis (y, x, or z). **b)** Density and peak orientations reconstructed by eGRL (right) and traditional SVD (left) algorithms from simulated raw data captured by wide-field microscopy (1.1NA), light sheet microscopy (SPIM, 1.1NA), and multi-view light sheet microscopy (diSPIM, 1.1NA and 0.67NA). Raw data has added Poisson noise for an SNR = 5 dB (see also **Supplementary Fig. 8**). **c)** Higher-magnification views corresponding to the helix in the red dashed rectangle region in **b**. Note eGRL provides better spatio-angular reconstruction than SVD in all microscope configurations, and the dual-view configuration (diSPIM) provides the best performance in estimation of density and peak orientations. **d)** Ground truth corresponding to the helix in **c. e)** Higher-magnification views of the diSPIM results corresponding to the dashed blue rectangle region in **b**. The yellow curved arrows indicate the expected orientation trend along the helix. Note the peak orientations of eGRL align much better with this expected orientation trend than those of SVD. The ODF maps of the cyan dashed rectangle region are also shown in the right cyan inset to confirm the closer match of the eGRL reconstructions to the ground truth. **f)** Quantitative analysis of the reconstruction performance on dual-view light sheet data (diSPIM) with two different noise levels (SNR=20 dB and SNR=5 dB), with SSIM, PSIM and ONCC quantifying the reconstruction accuracy of density, peak orientation, and ODF, respectively. **g)** Comparison of peak orientations reconstructed from synthetic shell data, showing orientation bias in traditional SVD reconstructions. Note the SVD produces peak orientations biased towards the y axis as observed in the xy plane also visualized by the pseudo color biased towards green, e.g., blue color shifts to cyan and red shifts to orange), while eGRL compensates for this bias. Radar chart shows the proportion of all peak orientations in x, y, z axes (ranging from 0 to 1), highlighting less biased peak orientation estimate offered by eGRL compared to SVD. Raw data has a noise level of SNR=5dB. Scale bars: **a, b** 2 μm, **c-e** 1 μm, **g** 2 μm. See also **Supplementary Figs. 5-8**.

When comparing the results across different microscopy configurations, the reconstructions from wide-field microscopy and light sheet microscopy (**Fig. 2b, c**, top and middle rows) exhibit limited axial resolution and orientation aliasing due to single-view polarization encoding. By contrast, the dual-view light sheet microscopy (diSPIM) implementation provides superior spatial resolution (density map) and the ability to resolve orientation (**Fig. 2b, c**, bottom rows), which we attribute to the improved information content when detecting from two orthogonal views and the more comprehensive 3D polarization modulation when exciting from the two views. These results highlight the necessity of dual-view polarization encoding and detection for better resolving complex spatio-angular distributions than their single-view counterparts.

We thus shifted our focus to algorithm performance on dual-view light sheet microscopy. We found SVD reconstruction introduces a significant orientational bias, clearly visible in the ODFs (**Fig. 2e**), while eGRL better matches the ground truth in terms of both peak orientations and ODFs. We further conducted quantitative analysis on eGRL’s performance by defining image quality metrics, including the SSIM for density, peak-orientational similarity index measure (PSIM) for peak orientation, and orientational normalized cross-correlation (ONCC) for the entire orientation distribution (**Methods**). As expected, eGRL outperforms the SVD algorithm on data corrupted with different Poisson noise levels as assessed by all metrics (**Fig. 2f, Supplementary Fig. 8, Supplementary Video 3**). We suspect the inferior performance of the SVD algorithm arises from its Gaussian noise model, which is mismatched to the usually dominant Poisson noise encountered in fluorescence microscopy, and by the bias arising from the regularization used for noise suppression^29,30^. In support of this assertion, when examining a spherical phantom with molecular orientations uniformly distributed and perpendicular to the sphere surface (**Fig. 2g**), the SVD reconstruction was always biased toward the Y axis (**Supplementary Video 4**), i.e., the direction orthogonal to the plane formed by the two views’ optical axes, whereas eGRL avoided this bias due to its implicit regularization(**Fig. 2g, Methods**).

### eGRL enables robust spatio-angular reconstruction for diverse samples

The performance of eGRL on synthetic datasets across different PFM modalities spurred us to investigate more samples acquired on our recent polarized dual-view light sheet microscope (pol-diSPIM)^18^ (**Supplementary Fig. 9, Methods, Supplementary Table 1**). The pol-diSPIM enables dual-view light sheet excitation and complementary orthogonal detection, flexible tuning of the polarization modulation in each illumination arm, and additional light sheet tilting^31,32^ to extend the polarization encoding in 3D space.

To confirm the reliability and accuracy of spatio-angular reconstruction in experimental imaging, we first used a calibration target image set of FM1-43 labelled giant unilamellar vesicles (GUVs, a spherical membrane model with dipole orientations normal to the surface^33^), acquired by pol-diSPIM with polarization modulations (9 from each view for a total of 18 volumes **Supplementary Table 2**). eGRL prediction restored the 3D spatio-angular distribution of the GUVs after 10 iterations (**Supplementary Video 5**), and visualizing the peak orientation (**Fig. 3a i**) and sliced ODF map (**Fig. 3a ii**) confirms that fluorophores displayed a dipole transition moment oriented normal to the membrane (**Supplementary Video 6**). For comparison, we projected all 3D orientations into a 2D plane to demonstrate the loss of information when considering only 2D orientations (**Fig. 3a iii** left), confirming the need for 3D PFM and reconstruction in 3D samples. Compared to SVD in the experimental GUV data, the angular distribution estimated by eGRL was more in line with the prior knowledge of FM1-43 dye bound within lipid vesicles, as evidenced by the orientation-encoded colors in dense visualizations (**Fig. 3b, Supplementary Video 7**). eGRL results on experimental data also better matched synthetic phantom GUVs stained with dipoles oriented normal to the GUV surface, further supporting the reconstruction (**Fig. 3b**).

**Fig 3.**
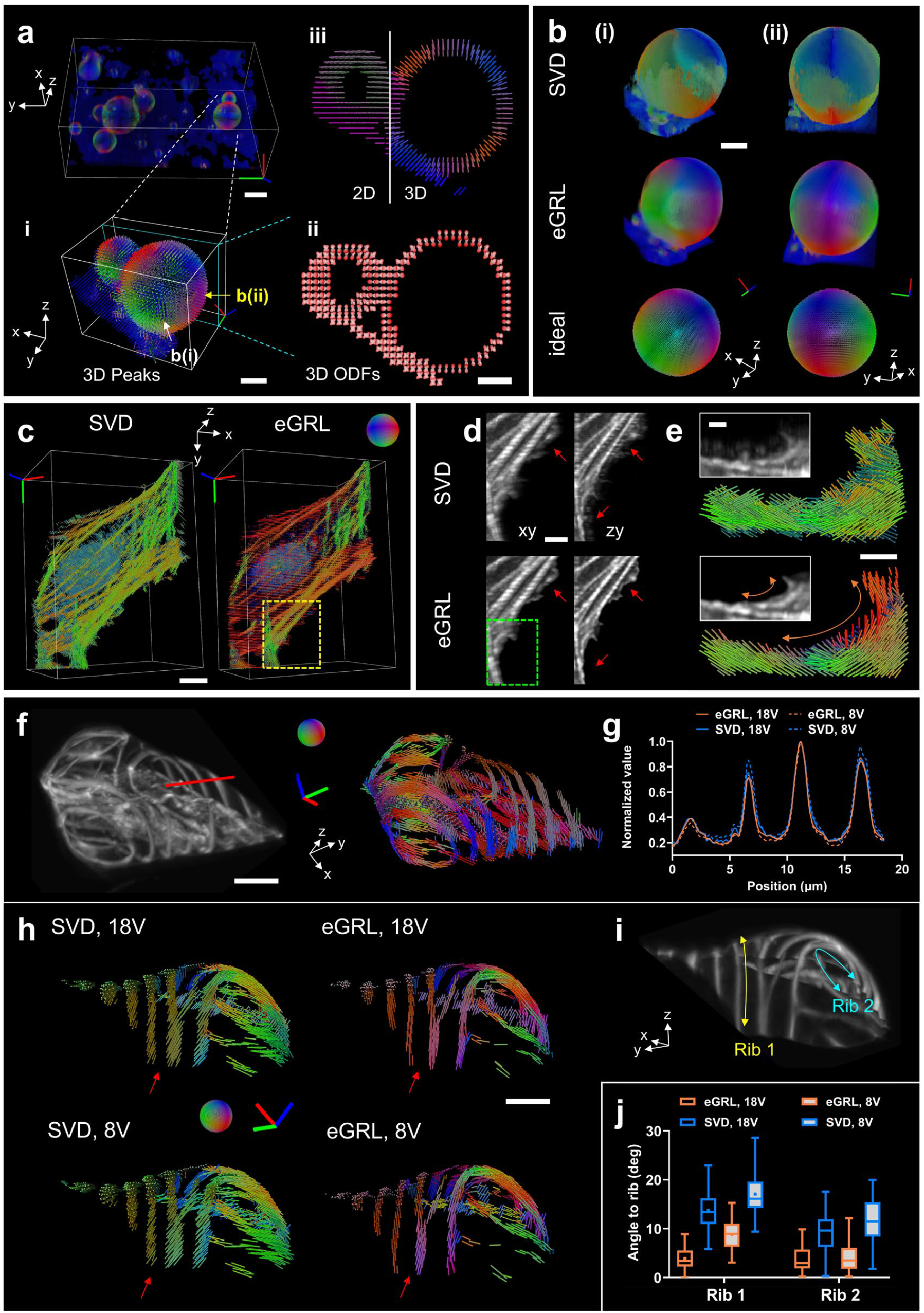
Spatio-angular structure reconstructions on multiple fixed samples. **a)** eGRL reconstruction of FM1-43 labelled GUVs acquired by pol-diSPIM. i, Higher-magnification view of a pair of GUVs (∼ 10 μm and ∼ 5 μm diameter); ii, ODF map of the slice indicated by the cyan dashed rectangle in i; iii, 3D peak orientation vs. 3D orientations projected into 2D plane corresponding to ii for comparison. **b)** The 10 μm GUV viewed from two different perspectives as indicated by the white and yellow arrows in **a**. SVD reconstruction (top), eGRL reconstruction (middle) and ideal peak orientation maps (bottom, for reference) are presented. Prior knowledge suggests that molecular orientations should be normal to the membrane surface, which is better approximated by the eGRL result than the SVD reconstruction. **c)** Reconstructions by SVD and eGRL of a fixed U2OS cell labeled with Alexa Fluor 488 phalloidin. **d)** The lateral (left) and axial (right) maximum-intensity projections from the density map of the yellow dashed rectangular region in **c**. Note eGRL produces a better resolved density map with fewer artifacts (red arrows). **e)** Higher magnification views of the density and peak orientation maps of the green dashed rectangular region in **d**. The orange curved bidirectional arrows indicate that the peak orientations reconstructed by eGRL smoothly change along the tangential direction, while those by SVD are disorganized. **f)** The density and peak orientation maps of a xylem cell (labelled by Pontamine fast scarlet) reconstructed by eGRL. **g)** Density profiles along the red line in **f** of the eGRL and SVD reconstructions with 18 and 8 polarization modulations (Scheme 1, 18V and Scheme 2, 8V. See **Supplementary Table 2**). **h)** Comparison of the peak orientations reconstructed with 18 and 8 polarization modulations by eGRL and SVD, showing that eGRL better preserves reconstruction than SVD even when reducing the number of polarizations from 18 to 8 by eGRL. Note the bottom of the cell has been cut away for easier visualization. **i)** Density perspective to accompany **h. j)** The angles of the peak orientations deviating from the rib directions at ∼260 points uniformly selected on each rib (yellow and cyan bidirectional arrows) in **h**. Scale bars: **a** top left 10 μm, **a** i-iii 5 μm, **b** 5 μm, **c** 10 μm, **d** 5 μm, **e** 2 μm, **f-i** 10 μm. See also **Supplementary Figs. 9-13**.

Next, we examined reconstructions corresponding to a fixed U2OS cell stained with Alexa Fluor 488 phalloidin, acquired under the same 18 polarization modulations via pol-diSPIM (**Supplementary Video 8**). The eGRL result showed differences between the orientation distribution of fluorophores close to the nucleus in predominantly one direction vs. those decorating flat or slender filaments, which showed a range of directions that correlated with the long axis of the actin filaments (**Supplementary Fig. 10**). In comparison, the SVD result tended to bias all orientations towards the y-axis (overall pseudo colors tend to be green, **Fig. 3c**). Overall, eGRL provides finer structural details with reduced blurring and fewer artifacts compared to SVD, as pointed out by the arrows (**Fig. 3d**), and more continuous local orientation detail along the peripheral arc structure at the edge of the cell (**Fig. 3e**).

To investigate the effectiveness of eGRL for reconstructing spatial and angular distributions, we constructed an ablated variant of eGRL termed eGRL-p (**Methods**), which applied an RL-deconvolution separately to all raw stacks of each polarization channel to restore the spatial distribution, followed by an independent voxel-by-voxel eGRL reconstruction to estimate the angular distribution at each spatial location. eGRL-p led to blurred density images and introduced structural artifacts (**Supplementary Fig. 11a**), also producing obvious errors in angular estimation (**Supplementary Fig. 11b, c**) resulting in an overall tendency towards isotropic ODFs (**Supplementary Fig. 11d**). These results can be attributed to the RL algorithm’s properties of sharpening the images (emphasizing certain local structures while lowering the background) and its slight nonlinear behavior^34^, so that deconvolving each polarization channel separately corrupts relationships between each polarization channel. These results underscore the significance of the joint estimation of spatial and angular distribution provided by the full eGRL model.

Since PFMs typically acquire image stacks sequentially under different polarization modulations, employing fewer polarization modulations offers a practical approach to reduce phototoxicity, acquisition time, and the burden of data storage and postprocessing. To compare the performance of eGRL and SVD under conditions of polarization downsampling, we analyzed data from the comprehensive 18 modulations (Scheme 1, 18 V. See **Supplementary Table 2**) to a reduced set of 8 or 6 modulations (Scheme 2, 8V or Scheme 3, 6V. See **Supplementary Table 2**) from a tobacco xylem cell labelled by Pontamine fast scarlet (**Fig. 3f, Supplementary Video 9**), the GUV data (**Supplementary Fig. 12**) and another fixed U2OS cell labeled with Alexa Fluor 488 phalloidin (**Supplementary Fig. 13, Supplementary Video 10**). Using fewer polarization modulations did not notably affect reconstructions for either eGRL or SVD, as assessed from the density maps (**Supplementary Fig. 13b, c**) and profiles (**Fig. 3g, Supplementary Fig. 13d-g**). The eGRL estimate consistently preserved the expected direction of the dipoles in Pontamine fast scarlet such that fluorophores tend to orientate along curved ribs in xylem cells (**Supplementary Video 11**), visually from magnified orientation views (**Fig. 3h, i, Supplementary Fig. 13h, i**), and quantitatively as assessed by the average angle between the orientation and ribs (**Fig. 3j**). The eGRL reconstruction with fewer modulations also surpassed the SVD reconstruction with all modulations, further underscoring eGRL’s superior performance over SVD.

### eGRL unveils distance-dependent attraction from suspended nanofiber networks to cellular actin distributions

Nanofiber scaffolds provide a versatile and efficient platform for the exploration of a wide variety of cellular phenotypes, e.g., cellular polarization (dependent on fiber length, diameter, and inter-fiber spacing) as well as cellular motion (motile fraction and locomotion speed)^35–39^. Compared to 2D flat substrates, these nanowires provide more physiologically relevant substrates to understand relationships between molecular architecture and cellular behaviors. Currently, the ability of nanowires to affect cell shape has only been assessed macroscopically through observations of cell morphology^40–43^. To advance our understanding of cell adhesion and cytoskeletal organization by external spatial cues at the molecular level^44^, we investigated how nanowires affect actin orientation as assessed using our eGLR reconstructions. We cultured NIH3T3 cells on nanofiber scaffolds with configurations of single, multiple parallel, and multiple crossed nanowires, stained the cells with Alexa Fluor 568 phalloidin, and imaged them using pol-diSPIM in two channels: a 561 nm channel with 6 polarization modulations for the actin channel, and a 488 nm channel without polarization modulation for labBDP FL maleimide labeled nanowires^18^.

First, we focused on a spindle-shaped NIH3T3 cell grown on a single nanofiber (**Fig. 4a**) and observed that molecules exhibit an overall orientation predominantly towards the nanofiber direction (y-axis) when considering only average orientation (**Fig. 4b**). However, we suspected that in addition to this general orientation tendency, there might also be more subtle orientation changes in the vicinity of the nanowires. To further examine this possibility, we utilized the Order Parameter (OP) metric, derived from the ODF (**Methods**), as a biological indicator to measure the proportion of molecules aligned in a specified direction at each position (**Fig. 4c, d**). OP by its definition provides a more comprehensive description of orientation than average orientation, as it assesses the degree to which an orientation ensemble is aligned to a particular direction (**Supplementary Fig. 14**). For each voxel, we calculated the OP value with respect to the nanowire’s direction and as a function of its proximity to the wire (**Fig. 4e**). We visualized the correlation between them (**Fig. 4f**) and found that OP increases as the voxel approaches the nanowire, suggesting that the nanowire exerts a distance-dependent directional influence on the orientation of probes. The fluorescent probes bound to the cytoskeleton firmly anchor along the direction of actin filaments, indicating that the nanowire promotes actin accumulation and growth in adjacent regions along the nanowire direction. In addition, the orientational aggregation of probes on actin correlates with the mechanical tension within these filaments^45^, thereby presumably mirroring the rise in local stress due to the cellular elongation along nanowires.

**Fig 4.**
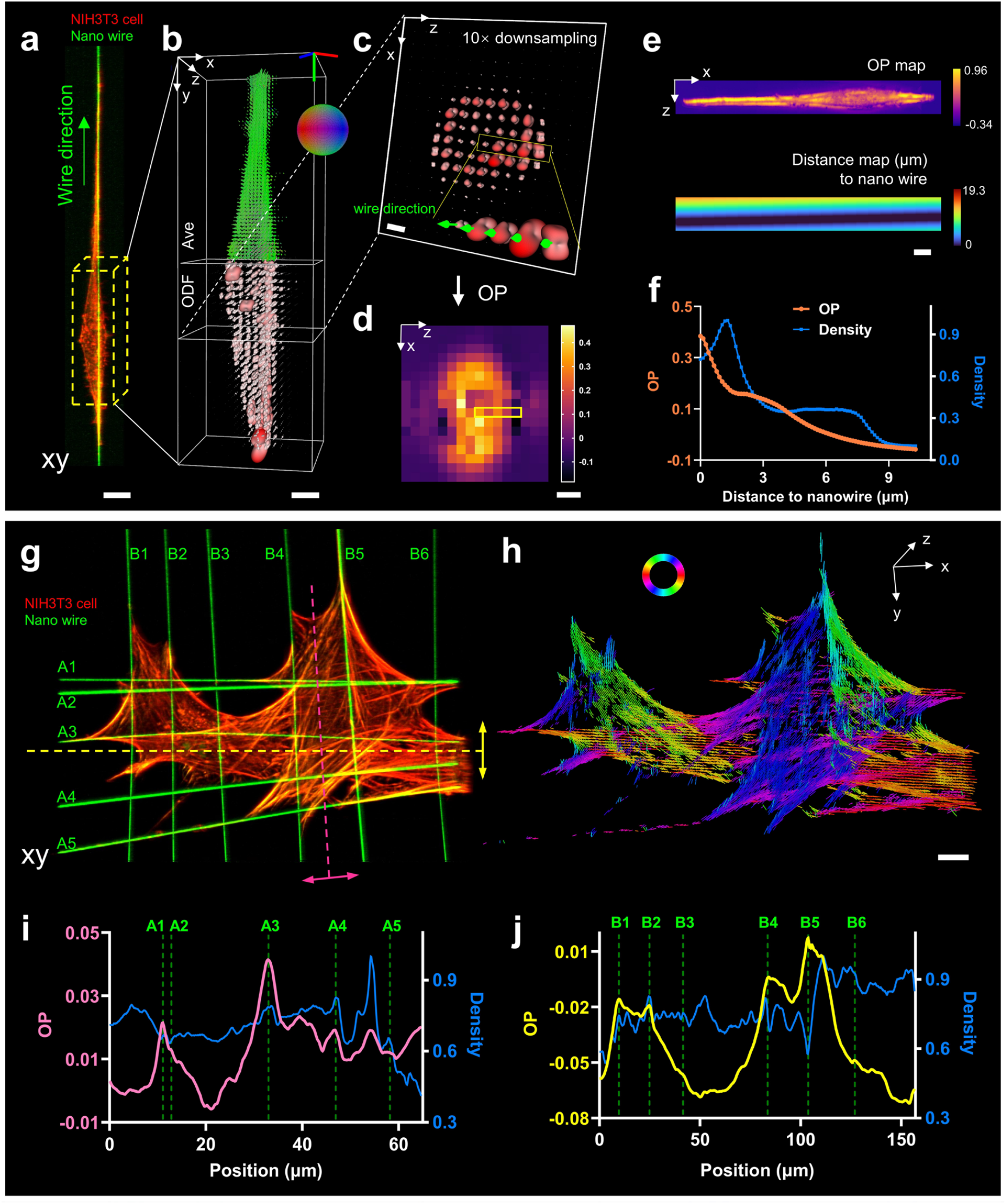
Revealing correlations between nanowire directions and the molecular orientation of actin. **a)** Maximum-intensity projection demonstrating the density map from reconstructions of a spindle-shaped NIH3T3 cell grown on single nanowire, labelled with Alexa Fluor 568 Phalloidin (red, polarized imaging with 561 nm excitation) and deconvolved results of nanowire (green, labBDP FL maleimide marker, non-polarized imaging with 488 nm excitation). The green arrow indicates the direction of nano wire. **b)** The 3D average orientation and ODF map reconstructed by eGRL of the yellow cubic region in **a**. The reconstruction has been downsampled 5-fold for visualization purposes. **c)** Magnified view of the slice marked in **b**, 10-fold downsampling for display purposes. In the higher-magnification view (inset), we compared each ODF with the wire direction (green arrows) for visually observing the degree to which the ODFs tend in this direction. **d)** Heat map showing the normalized Order Parameter (OP) over the region in **c** in the direction of the nanowire. The yellow rectangular region corresponds to the OP values of the yellow rectangular region in **c. e)** The entire OP map (top, 3D but shown in maximum-intensity projection) and the distance to the nanowire (bottom, 3D but shown in minimum-intensity projection). **f)** OP vs. the distance to the nanowire curve from the two maps in **e** (averaging OP values of all voxels with the same distance value), indicating that dipole orientations nearer the nanowire tend to align along the nanowire direction. **g)** Density maximum intensity projections showing another fixed NIH3T3 cell (in red color) grown on crosshatched nanowires (in green color) as for **a**. All the nanowires are numbered based on the approximate direction (A for horizontal wires, B for vertical ones). **h)** The peak orientation map of the actin in **g** reconstructed by eGRL, with a different colorbar/colormap encoded in 2D XY plane due to the narrow thickness of cell. **i)** OP profile along the pink dashed line (average within 5 μm of the line) in **g**, with the OP direction defined as the bidirectional pink arrow. The OP profile passes through wire A1-A5 and the corresponding wire positions are marked on the profile. Note the wires are mostly coincident with the local maxima on the OP curve. **j)** OP profile along the yellow dashed lines (average within 5 μm of the line) in **g**, with the OP direction defined as the bidirectional yellow arrow. The OP profile passes through wire B1-B6. As in **i**, the wires often coincide with the local maxima on the OP curve. Scale bars: **a** 10 μm, **b** 5 μm, **c** 2 μm, **d** 4 μm, **e, g, h** 10 μm. See also **Supplementary Figs. 14-16**.

Similarly, we observed such distance-dependent orientation when examining actin in cells grown on arrays of parallel or crossed nanofibers that had spindle, two-fiber and crosshatched morphologies (**Fig. 4g, h, Supplementary Fig. 15**). At the periphery of cells, the density and orientation maps revealed the arc-shaped actin filaments followed the cell shape between the nanowires with the orientations aligning tangentially to these structures, while there was no distinct orientational preference observed in most of the internal cellular regions (**Fig. 4g, h**). However, when examining those internal regions that intersected with several parallel nanowires, we found that peaks in the OP profiles were often coincident with the location of the nanowires (**Fig. 4i, j**), suggesting again a strong distance-dependence between nanowire influence and actin orientation.

Next, we examined the variation of OP along one nanofiber when it passed in the vicinity of fibers running in other directions. On crosshatched-shaped cells grown on intersecting nanofibers (**Supplementary Fig. 16a, b**), we considered regions surrounding several wires and drew the OP profiles in the direction of these wires (**Supplementary Fig. 16c, d**), finding that nanowires running in the orthogonal direction were coincident with the local minimum on the OP curves. The presence of extra nanowires appears to redirect the actin filaments towards their direction, resulting in an apparent competition between nanowires running in different directions which likely all guide actin alignment. We note that these observations would not be possible via intensity-based microscopy^46^ (**Fig. 4f, i, j**) or could be obscured in conventional PFM^47^ that relies on average orientation measurements. When considering average orientation, the tendency of molecules aligning with the target nanowire can be assessed only by their parallelism (how parallel the average molecular orientation is to the nanowire). Compared with a parallelism measure, the OP obtained from the 3D spatio-angular distributions present much higher contrast peaks or valleys, depending on the relative orientation of actin filaments and nanowires (**Supplementary Figs. 15, 16**). We noticed particularly strong OP signatures when considering actin in the vicinity of paired nanowires, when overall average orientations exhibit strong alignment with the direction of nanowires (**Supplementary Figs. 15a, b**).

### eGRL enables observation of spatio-angular dynamics in living cells

Given the success of our reconstruction method on fixed samples, we proceeded to apply it to live cells, where imaging is more challenging due to the necessity for rapid imaging, lower SNR in the data, and the size and computational burden associated with large time-lapse data.

First, we labeled actin filaments in a HeLa cell with Sir-Actin dye (640 nm excitation) and collected volumetric time-lapse images over 20 min at 1 min intervals between volumes, and processed them with our eGRL framework. Our reconstructions show that principal orientations (the maximum projection direction of ODF, comprehensively considering all the local peaks and demonstrating the overall alignment of ODF, **Supplementary Fig. 1i**) of SiR-Actin probes align perpendicular to the actin filaments (**Fig. 5a, b**). While the orientations in the flat regions of the thin cell are mostly in the plane of the coverslip (XY plane, **Fig. 5b** i, ii), we also found interesting exceptions in the vicinity of cell protrusions (**Fig. 5b** iii). Intriguingly, the protrusive orientations exhibited a gradually rotating pattern (roughly in the YZ plane) along the periphery of the protrusion (**Fig. 5c** i) yet remain perpendicular to actin filaments^48^. These orientations are not randomly distributed within the YZ plane, and we suspect this distribution may indicate ‘twisting’ within the slender actin bundles (**Fig. 5b** iii). The dynamics within the protrusion were further revealed by the orientation-encoded colors in time-lapse images (**Supplementary Fig. 17b**, right) as well as the x axis-projection orientation distribution along the contour of the protrusion (**Supplementary Fig. 17c, Supplementary Video 12**). These dynamics are not obvious in the density images (**Supplementary Fig. 17b**, left), demonstrating the potential of our method for observing 3D orientations that would be obscured in conventional imaging. Moreover, the significantly higher orientational changes in the actin filament of the protrusion, compared to actin elsewhere in the cell, can serve as an internal control to rule out the possibility that the observed dynamics are merely random fluctuations caused by photobleaching or noise (**Fig. 5c** ii). As a consistency check, we used eGLR reconstructions to compare the orientation of the actin probes Phalloidin^49^ (**Supplementary Fig. 18a, b**) and Sir-Actin (**Supplementary Fig. 18c, d**) in fixed and live HeLa cells. The Phalloidin probes generally aligned parallel to the direction of actin filaments, in contrast to Sir-Actin, which displayed perpendicular orientations especially at the cell edge and within looped actin structures in the middle of cells. These reconstructions are consistent with earlier observations^48^ and demonstrate the effectiveness of our method.

**Fig. 5.**
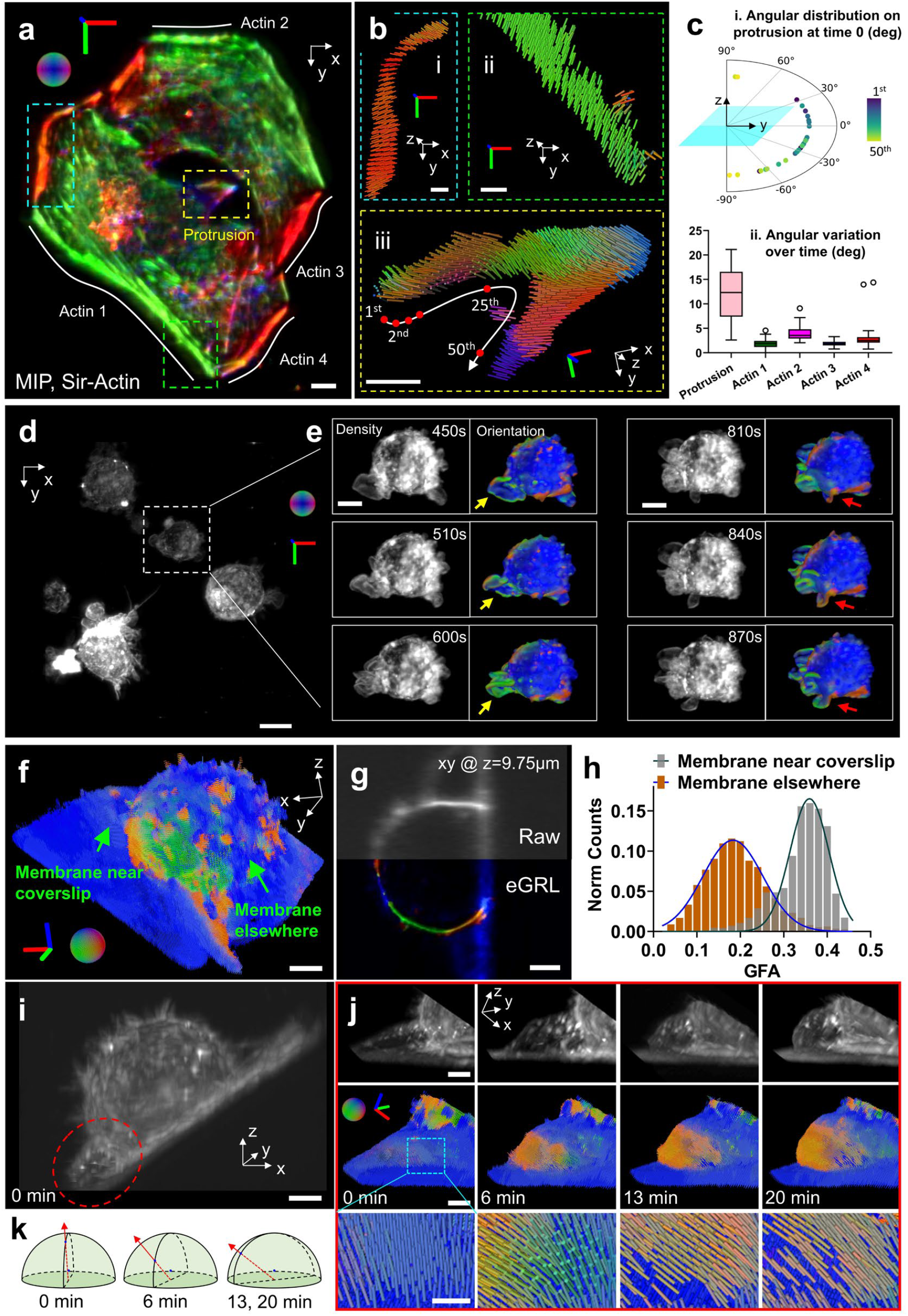
Observing spatio-angular dynamics in living cells. **a)** The orientation map of a live HeLa cell labelled with Sir-Actin and imaged with pol-diSPIM under 640 nm laser. **b)** Magnified views of the actin periphery and protrusion corresponding to the cyan, green and yellow dashed rectangular region in **a**. Note in protrusion region iii, the probes don’t align parallel to the coverglass, instead, they are organized in a pattern of continuous spatial rotation on the plane perpendicular to actin’s tangent direction. **c)** i, How principal orientation distributes along the actin filament of protrusion. 50 characteristic points were selected uniformly along the trail in **b** iii. The polar plots are shown for the inclination angle of the principal orientation relative to the x-y plane, with pseudo color to indicate the points’ order (1^st^ ∼ 50^th^) along the trail. ii, Time-lapse orientational dynamics in protrusion (yellow rectangle region in **a**) in contrast to measurements elsewhere in the cell (Actin 1∼4 in **a**). Around 40 points are counted along each region and the box plot is shown in the Tukey style (range from the first quartile to the third quartile, whiskers extending to 1.5 × interquartile range), with outliers marked by circles. Quantification is performed via angular variation, which is defined as the standard deviation of orientation changes over time. **d)** 3D density map of multiple FM1-43 labelled live macrophage cells. **e)** Example time series of the cell from the square region in **d**, comparing the density and principal orientation maps, demonstrating topological dynamics and orientation variation with the bending and folding of the pointed protruding membranes. **f)** The principal orientation map of a live U2OS cell labeled with FM1-43. **g)**, Comparison of raw data and the eGRL reconstruction colored by the principal orientation, in the lateral slice at z depth of 9.75 μm. **h)** Histogram of Generalized Fractional Anisotropy (GFA) at the initial time point, showing the higher GFA in the vicinity of the coverslip surface (coated with poly-L-Lysine) than elsewhere on the cell membrane, likely because the former constrains the membrane surface more than elsewhere in the cell. **i)** Density map of the U2OS cell at the initial time point where an emerging membrane protrusion is marked by the red ellipse. **j)** Time-lapse images of the principal orientation demonstrating the expansion of cell membrane (top row). Orientations initially perpendicular to the membrane move towards the membrane expansion direction (middle row, where the cyan regions are magnified in bottom row. Only part of the cell is shown for display simplicity). **k)** Schematic diagram of orientation tendency in bottom row of **j**. Scale bars: **a** 5 μm, **b** 2 μm, **d** 10 μm, **e** 3 μm, **f, g, i** 5 μm, **j** top and middle 3 μm, **j** bottom 1 μm. See also **Supplementary Figs. 17-23**.

Next, we applied our method to investigate spatio-angular distributions and dynamics in thicker live cells. We imaged a living macrophage cell labeled with FM1-43 every 30 s for 30 time points spanning 15 min (**Supplementary Fig. 19a**). We observed the collapse of membrane protrusions (**Supplementary Fig. 19b**, from 0 to 10 min) and its transition as the cell blebs (**Supplementary Fig. 19b**, 15 min), although FM1-43 orientations were maintained perpendicular to the membrane (**Supplementary Video 13**).

In addition, we captured a larger time-lapse dataset showing multiple living macrophage cells (**Fig. 5d**) imaged at the same rate and over the same time interval. During the spatio-angular reconstruction process, we noticed that this dataset (700 × 700 × 400 voxels × 6 modulations × 30 timepoints, roughly 91 × 91 × 52 μm^3^, ∼ 66 GB) required 1 TB RAM and thousands of hours to compute the reconstructions for all timepoints, even after the acceleration afforded by multi-threaded parallel computing. This bottleneck prompted us to refine our processing pipeline, whereby we cropped each volume into 320 subvolumes (each 100 × 100 × 100 voxels × 6 modulations), performed the eGRL prediction on each subvolume, and spatio-angularly stitched the results back together (**Methods**). This pipeline took roughly 5.1 hours on a single workstation equipped with a consumer-grade graphical processing unit (GPU) card, achieving an over 200-fold acceleration compared with direct processing of the entire volume. The eGRL reconstruction displayed rich orientation dynamics in membrane ruffles and cell protrusions that were impossible to observe in the accompanying density maps (**Fig. 5e**). Besides the orientational changes in lamellipodia that accompanied structural movements over time, we also captured patches of highly ordered membrane (**Supplementary Fig. 20**) that appeared to ‘traffick’ in a retrograde fashion on protrusive filopodia (**Supplementary Video 14**). These dynamic patches of ordered membrane could be structural movements of the membrane, either as twisting of the whole structure, or small lamellipodia-like ruffles as have been previously observed on filopodia^50^. Alternatively, these patches may represent retrograde trafficking^51^ of membrane microdomains in macrophage filopodia, which are cholesterol rich^52^, and may therefore differentially order FM1-43^53^. Further work colocalizing FM1-43 polarization with protein markers for lamellipodia-like ruffles or lipid-dependent clusters^50^ could distinguish these possibilities. In either case, the ability to easily resolve these dynamic structures in filopodia represents an improvement over other methods to observe membrane order, which require fluorescence lifetime imaging microscopy or ratiometric imaging of dyes such as Flipper-TR or Laurdan^54,55^.

To further validate our method’s ability to observe spatio-angular dynamics in live cells, we compared the sensitivity and orientation characteristics of different dyes. We examined samples labeled with two dyes, one with known sensitivity to polarized excitation and the other insensitive, by using dually labeled U2OS cells stained with FM1-43^10,56^ (sensitive, **Supplementary Fig. 21a**) and RFP-PH (insensitive, **Supplementary Fig. 21b**). The reconstructed orientation and GFA maps of polarization sensitive FM1-43 dye exhibited oriented and anisotropic ODFs, while those ODFs corresponding to polarization insensitive RFP-PH tended to exhibit more isotropic character (dispersed distribution without preferred orientation, **Supplementary Fig. 21c, d, Supplementary Video 15**). For the polarization sensitive FM1-43 dye, the spatio-angular distributions showed orientational selectivity consistent perpendicular to membranes of cellular structures with this dye.

We then analyzed the spatio-angular membrane dynamics in FM1-43 labeled U2OS cells. The cells were adhered on glass coverslips using poly-L-Lysine (PLL), and imaged with pol-diSPIM every 20 s over 60 timepoints spanning 20 minutes. The eGRL restoration highlighted the structure and orientation of the cell membrane (**Fig. 5f, g, Supplementary Video 16**), showing that FM1-43 fluorophores tend to be oriented perpendicular to the membrane surface. We used Generalized Fractional Anisotropy (GFA, see **Methods**) to quantify changes in membrane orientation near and far from the coverslip. We found that membranes displayed higher GFA near the coverslip than elsewhere on the cell (**Fig. 5h**, see also polarization-insensitive negative control in **Supplementary Fig. 21d**, excluding motion as a cause of differential GFA). Over time, the membrane originally adjacent to the coverslip (**Fig. 5i**) began to protrude (**Fig. 5j**, top row), showing accompanying changes (**Fig. 5j**, middle and bottom rows) in the principal orientation of molecules from perpendicular to the membrane surface towards the direction of overall membrane expansion (**Fig. 5k**).

## Discussion

Inspired by Richarson Lucy deconvolution, our eGRL framework incorporates statistical reconstruction methods into spatio-angularly coupled fluorescence microscopy, to improve estimation of 3D structure and orientation distributions compared to previous work relying on SVD^18^. eGRL achieves excellent noise suppression (**Supplementary Figs. 4, 8, Supplementary Video 3**), which helps to accurately estimate structure, orientation, and dynamics in biological samples. We devoted particular effort to minimizing the computational cost in the algorithm, as our two-stage mathematical improvement (angular domain transformation and iterative framework restructuring) and code-based GPU acceleration provide a 280-fold improvement in processing time relative to the underlying GRL algorithm, putting it on par with SVD reconstruction (eGRL 5.3s using 10 iterations vs. SVD 2.1s, tested on a patch with 100 × 100 × 100 voxels × 42 modulations both with GPU acceleration).

Our demonstrations in living cells were limited to relatively slow dynamics due to the need to capture multiple polarization modulations for full 3D polarized fluorescence imaging. With the acquisition setup and experimental conditions in this work, we find the eGRL reconstruction is robust to motion of at least the several hundred nm scale between polarization modulations, e.g, the reconstruction bias is limited to ∼5% in terms of PSIM when the motion is less than 1 μm (**Supplementary Fig. 22**), and is more robust still if the motion is random (**Supplementary Fig. 23**). Samples that move more rapidly would demand further improvements in imaging speed, which likely entails shorter exposure times or fewer polarized excitation cycles (potentially achievable via polarization splitting on the emission side). Recently, advances in deep learning have accelerated and enhanced spatial deconvolution, e.g., Richardson-Lucy network^34^ (RLN), whose structure is inspired by RLD but achieves superior performance, using single-view input to yield results comparable to dual-view deconvolution. We anticipate similarly transforming eGRL into a neural network that enables parameter-free setup, faster reconstruction speed, enhanced noise tolerance, and perhaps reduced polarization/view requirements, implying faster image acquisition. Additionally, we plan to explore universal methods for designing algorithms or networks that can maintain polarization relationships after separately processing images from each polarization modulation (which usually distorts the estimation as described in **Supplementary Fig. 11**), thereby providing greater flexibility for others to design preprocessing networks.

Finally, we anticipate further progress towards quantitative correlation between molecular spatio-angular indicators and biological parameters. Measurements on aligned fiber networks have recently been used to monitor cellular forces and contractility in spheroids^43^, which may pave the way for future developments on contactless force sensing through optical polarization.

We are also encouraged by new developments in genetically encoded, polarization-sensitive reporters which can be used to study actin filament assembly geometry and dynamics^57^. By integrating the growing family of such reporters with PFM and eGRL, we anticipate many insights into cellular function.

## Methods

### Spatio-angular modeling and reconstruction

#### Dipole PSF modeling

Generally, a polarized fluorescence microscope can be broken down into excitation and detection subsystems. We separately modeled these two systems to facilitate their combination, thereby enabling compatibility with a wide range of microscope types.

For polarized excitation from a single viewing direction, we split it into the illuminance PSF(**r**) (giving the distribution of illumination intensity) and an independent polarization excitation function 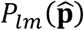:

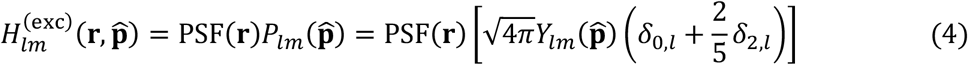

where the spherical harmonics function is indexed by degree *l* and order *m*, with |*m*| < *l*. And *δ*_*a*,*b*_ equals 1 when *a* = *b*, else equals 0. *Y*_*lm*_(**ŝ**) is the spherical harmonics base function.

For detection, we assume that the imaging system is laterally shift-invariant, so the detection dipole PSF can be modeled as:

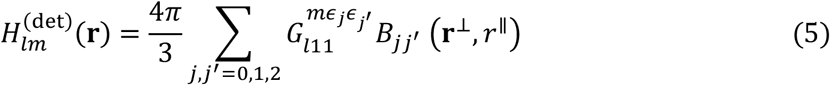

in which **r**^⊥^ denotes the lateral position while *r*^∥^ is the defocus position. *ϵ*_0_ = 1, *ϵ*_1_ = −1, *ϵ*_2_ = 0, and *G* is the real Gaunt coefficient. The irradiance 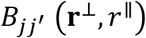 created on the detector is given by:

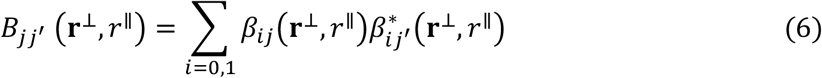

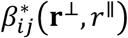 denotes the complex conjugate of *β*_*ij*_(**r**^⊥^, *r*^∥^) where *β*_*ij*_(**r**^⊥^, *r*^∥^) is the *i*th component of the electric field created at position **r**^⊥^ on the detector by the *j*th component of a dipole:

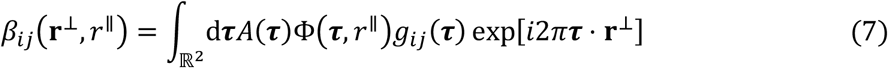

where *A*(***τ***) is the aplanatic apodization function with 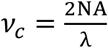:

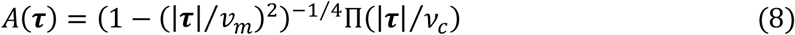

in which Π is a rectangle function defined as:

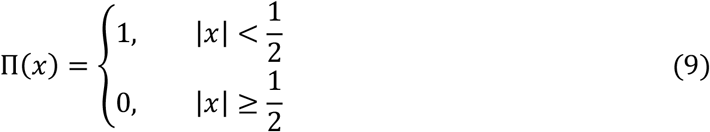

and Φ(***τ***, *r*^∥^) is used to encode the defocus phase with 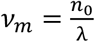:

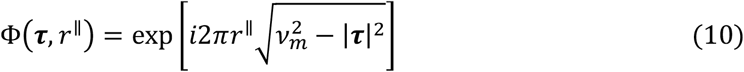

then *g*_*ij*_(***τ***) models the *i*th field components in the pupil plane created by the *j*th component of a dipole:

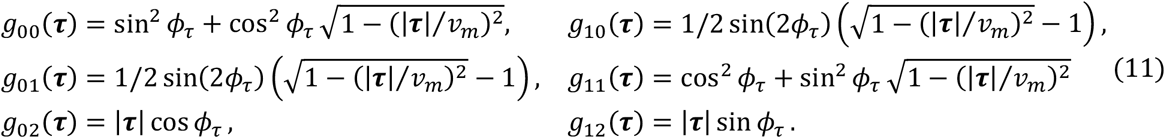

For a polarized wide-field microscope with uniformly polarized illumination and epi-fluorescence detection, we can directly combine illumination and detection via spherical harmonics multiplication:

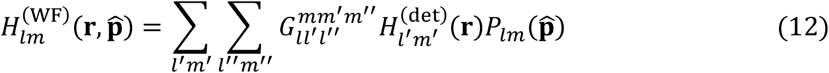

A polarized selective plane microscope with polarized light sheet excitation, enables optical sectioning. We assume that light sheet does not broaden appreciably along its propagation direction:

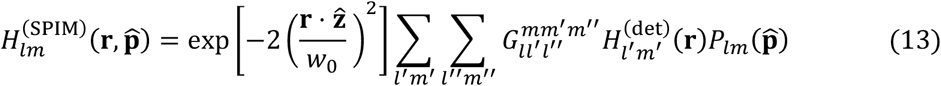

Further, pol-diSPIM adds another orthogonal view whose optical axis is along the x direction, resulting in improved lateral/axial resolution and larger angular bandwidth. The dipole PSF for the additional view is:

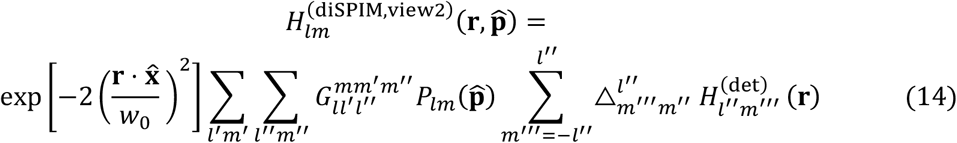

where △ represents the real Wigner D-matrices for the rotation of spherical harmonics coefficients, and the △ here (90° rotation across y axis) can be written as a matrix:

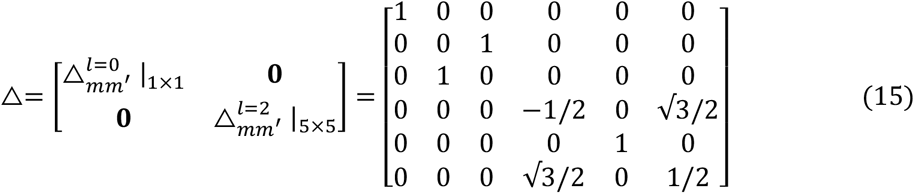

#### Iterative framework of Efficient Generalized Richardson-Lucy algorithm (eGRL)

Based on the spatio-angular imaging process and the typical noise characteristics of cameras, we derived the following iterative formula (see **Supplementary Note 1** for derivation) via maximum-likelihood expectation-maximization (MLEM) theory:

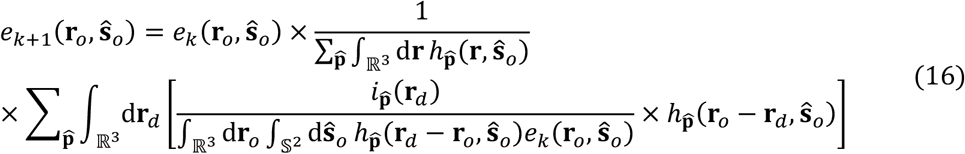

The process can be broken down into five steps of forward projection, division, back projection, normalization and update, and we have termed this the generalized Richardson-Lucy (GRL) algorithm. Note that 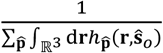 is referred to as object space sensitive constant used for normalizing the orientation channel, which is extremely important here to reinforce algorithm convergence but usually ignored in traditional RLD due to the pre-normalization of the PSF.

The initial GRL baseline provides accurate reconstructions but introduces a significant computational burden. By performing CPU multithreaded parallel computing on a computer workstation equipped with 2 Intel Xeon Platinum 8369B 2.90GHz CPU (totally 64 Cores / 128 Threads), GRL takes 278GB memory and 66 seconds per iteration to process an 80MB raw dataset with 10^6^ voxels and 42 polarization modulations, which might explain why iterative algorithms have not yet played a major role in such inverse problems. Therefore, we 1) approximated the orientation ensembles accurately with only a small number of spherical harmonic coefficients due to the angular band-limited nature of the microscope^17^, and accordingly developed methods to enable all variables, functions, and operations to be described in the spherical harmonic domain throughout the entire process, 2) optimized the backward projection strategy which further greatly reduces memory and time consumption (see **Supplementary Note 2** for details). With this two-step optimization, we obtain our final reconstruction framework:

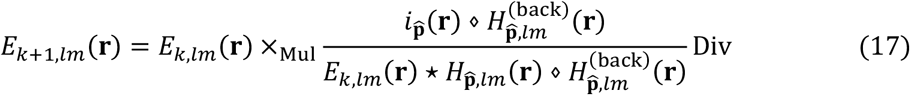

which is termed efficient generalized Richardson-Lucy (eGRL) algorithm, abbreviated as:

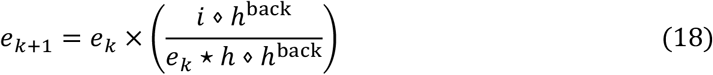

for clarity in the following descriptions.

eGRL consists of 2 precomputed steps and 3 steps in each iteration: FP+BP, DV, and Update. For the two (precomputed) steps of *i* ⋄ *h*^back^ and *h* ⋄ *h*^back^,

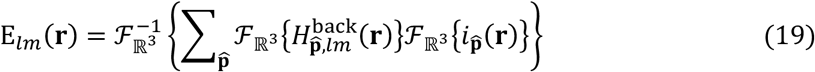

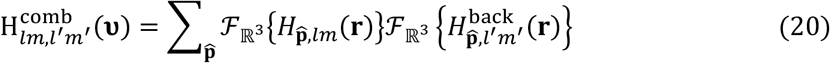

For the FP+BP step, with the combination of forward and back projection in *e*_*k*_ ⋆ *h* ⋄ *h*^back^, iterations no longer go through a ‘object->image->object’ process but a direct ‘object to object space’ transformation:

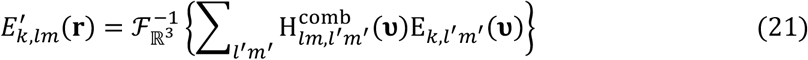

For the next DV step, we calculate the error map between the two projections *i* ⋄ *h*^back^ and *e*_*k*_ ⋆ *h* ⋄ *h*^back^, applying spherical harmonics division:

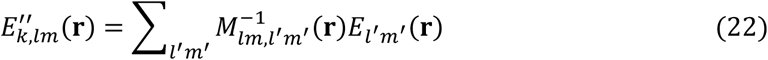

where:

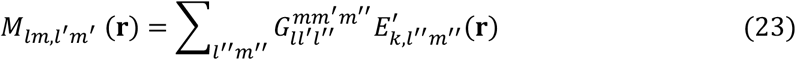

in which, 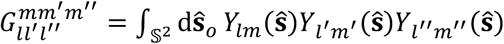 is the real Gaunt coefficient calculated by spherical harmonics function *Y*_*lm*_(**ŝ**). Note that 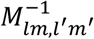 is the inverse of the matrix between *lm* and *l*^′^*m*^′^ channels.

For the final Update step, the k-th estimation is updated by the error map Eq. (22)and then we obtain the (k+1)-th estimate:

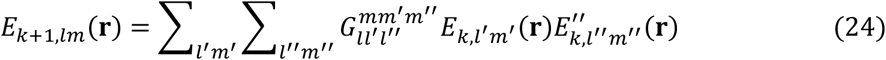

The entire eGRL iteration is given by:

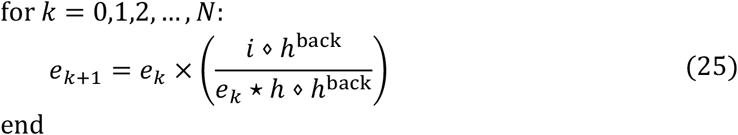

where the iteration number *N* is recommended to be set to 10-20 based on empirical observation. And the initial estimation is usually a uniform angular distribution while the density is set to the average of raw data 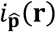 in polarization channels:

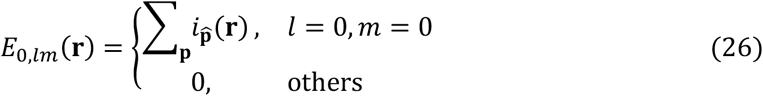

Moreover, we clarify that the baseline GRL is derived under the MLEM requirement of non-negativity for both the system matrix and the object. Similar to the Fourier-domain implementation of the RL algorithm, the optimized eGRL operating in the frequency and angular spectrum domains does not violate the non-negativity constraint in the spatial domain. In addition, eGRL with iterative framework modified is closer to Iterative Space Reconstruction Algorithm (ISRA, which minimizes a least-squares objective)^24^ than RL, however, our simulations show that GRL and eGRL performs similarly under Poisson noise (**Supplementary Fig. 4**).

For dual-view polarized microscopes, e.g. pol-diSPIM, eGRL can be extended by alternately considering polarization modulations in each view:

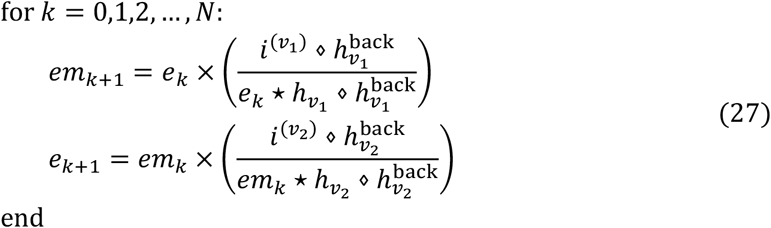

where the initial estimation is given by the average of images in polarization and view channels:

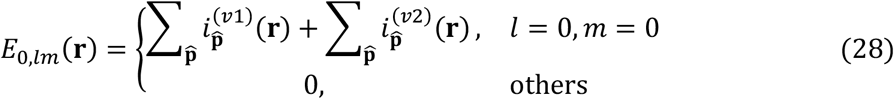

We recommend 10-iteration alternating dual-view eGRL deconvolution as a good rule of thumb in pol-diSPIM to balance processing time and spatio-angular resolution.

### Comparing methods

#### The ablated eGRL-p (spatial and angular decoupling)

For a series of polarization images without spatial blurring, each independent pixel can provide an angular estimate based on the pixel-level intensity under different polarization modulations. But for optical images blurred with point spread function, we developed the eGRL scheme for spatio-angular joint estimation. To verify the superiority of this coupling, we designed an ablation experiment of splitting the density and orientation estimation with the following steps:

1. Density estimation. We apply RL deconvolution to each polarization channel and obtain a series of polarization images with spatial blurring recovered.
2. Orientation estimation. We hypothesize that pixels in the deconvolved images are independent, so these can be reconstructed to give angular estimates based on the pixel-level intensity under different polarization modulations. We term this pixel-wise angular estimation method derived from eGRL as eGRL-p:

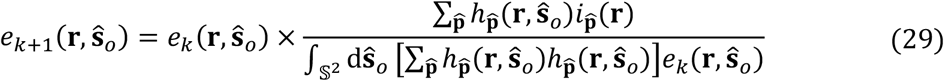

Further, we perform calculations in the spherical harmonics domain:

For precomputation steps:

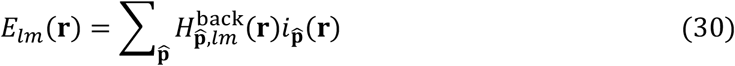

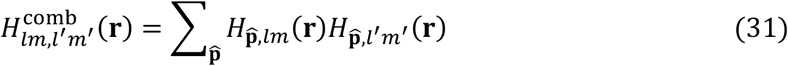

For the FP+BP step:

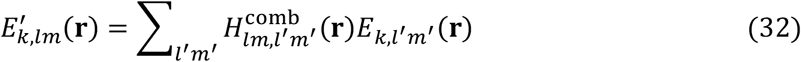

For the DV step:

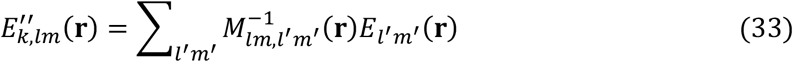

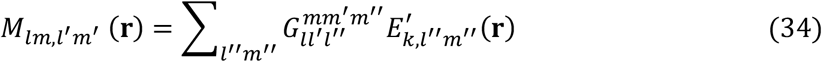

For the final update step:

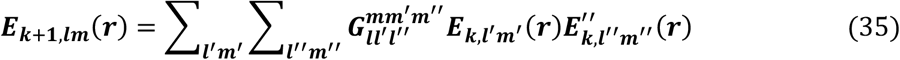

#### Analytical solution based on singular value decomposition (SVD)

The forward imaging process of polarized fluorescence microscope, as described in Eq. (1), can also be written in frequency and spherical harmonics domain as:

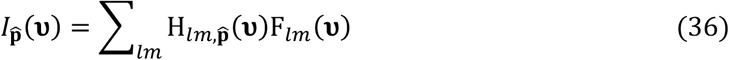

Then rewrite *R*-rank H_*lm*_ in terms of its SVD:

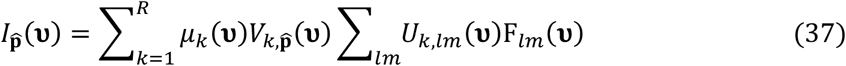

a Tikhonov pseudoinverse can be directly applied for estimation:

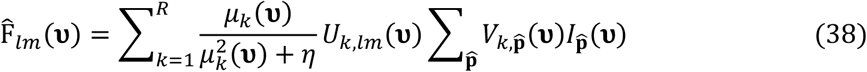

For dual-view polarization imaging, SVD requires us to concatenate the dipole PSFs of two views in polarization channel, and perform the same operation for the images. Then execute the original process:

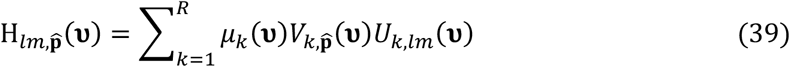

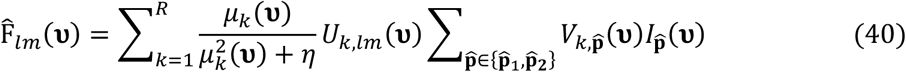

The regularization parameter *η* for simulation data is optimized by searching from 10^−8^ to 10^0^ in a step of 10^0.1^. The optimal value is determined by evaluating the reconstruction quality using PSIM indicators compared with GT. For experimental pol-diSPIM data, *η* is manually set to ∼ 0.1 empirically according to the visual reconstruction noise and prior biological information.

### Experimental data configuration

#### Polarized dual-view light sheet microscope (pol-diSPIM)

The pol-diSPIM system has been described in our previous paper^18^. It is built on an asymmetric dual-view light sheet geometry with a pair of water-immersion objectives: a 25×, 1.1 NA lens, and a 28.6×, 0.67 NA lens. The system is equipped with 3 lasers (488 nm, 561 nm, and 640 nm) for multicolor imaging. The dual-view optics are designed for 50× and 28.6× magnification, respectively, with object-space pixel sizes of 130 nm and 227 nm. While capable of operating as a conventional dual-view light sheet microscope^27^, the system offers additional functions to perform polarization modulations in the excitation paths by: (1) a liquid crystal (LC) module in each excitation path to customize the polarization orientation of the laser beam, and (2) a MEMS mirror to steer the beam propagation and tilt the illumination light sheet (**Supplementary Fig. 9**). The LC modules and MEMS mirrors together enable the system to explore a large set of possible polarization modulations (**Supplementary Table 1**).

#### Pol-diSPIM data acquisition and pre-processing

Data acquisition. The pol-diSPIM system has three degrees of freedom that can be used to estimate the orientation of the imaged molecules: transverse illumination polarizations, illumination tilt angles, and imaging views. We modulate the illumination beam’s transverse polarization with an LC module and modulate the tilt of the illumination beams with the MEMS mirror in either path.

We acquired volumetric data by scanning the sample stage through a stationary light sheet, acquiring fluorescence images with 15–50 ms exposures and 1 μm per stage step for each image. We acquired image volumes in both views before we changed the polarization/tilt state and acquired the next pair of volumes from both views. For multicolor imaging, all views, polarizations, and tilts for one color were acquired, followed by all views, polarizations, and tilts for the next color until all colors were acquired. In summary, we acquired datasets with as many as eight dimensions, and each dimension was collected in a loop in the following order from fastest to slowest: camera frame, stage scan positions, views, polarization/tilts, colors, and time points.

Data pre-processing. A) Deskewing. Since volumetric data were acquired by scanning the sample stage, raw images were deskewed, then cropped to save storage and computational expense. B) Registration. After deskewing, the two view images were interpolated and upsampled to an isotropic voxel size of 130 nm × 130 nm × 130 nm. Then View B (1.1 NA objective illumination and 0.67 NA objective detection) images were rotated by −90 degrees about the y-axis so that they were coarsely aligned to View A (0.67 NA objective illumination and 1.1 NA objective detection) images. To estimate a registration transformation, we averaged the volumes acquired under different illumination conditions (polarizations, tilts) into a single volume for each view, then estimated the 12-dimensional affine transformation that maximized the cross correlation of the two volumes. Finally, we applied the estimated transformation to all of the raw data volumes acquired in view B. All optimizations and transformations were performed with GPU-based 3D affine registration routines. c) Calibration. The use and realignment of the microscope may cause measurable drift of the illumination states, so we acquire the calibration data from a spatially and angularly uniform sample, an auto-fluorescent plastic slide (Chroma 92001) for the average intensity 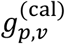 under different polarization *p* and view *v*. Then, due to the cos^2^ relationship between polarization angle and intensity, we calculate the phase shift of polarization by fitting this curve to obtain the deviation between the actual polarization angle and the program set. After getting the actual polarization angle, we generated the system dipole PSF and simulated the ideal average data 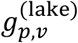 for imaging fluorescent lake (uniform intensity, isotropy ODF) under the same instrument configuration. The raw data 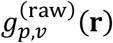 could be corrected by the following form:

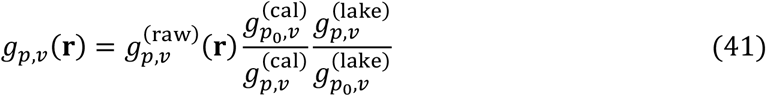

#### Computer hardware

All the simulation, PSF generation, and reconstruction algorithms (SVD, eGRL-P, eGRL) were implemented with Python version 3.9 in Windows 10 Pro 22H2 operating system.

The tests were performed with CPU multithreads parallel computing for acceleration (using the Joblib 1.1.0 package) or GPU-based acceleration (using the Cupy 8.3.0 package), on a computer workstation equipped with 1 TB memory, 2 Intel Xeon Platinum 8369B 2.90GHz CPU (totally 64 Cores / 128 Threads), and one Nvidia GeForce GTX 4090 GPU with 24 GB memory.

#### GPU-based postprocessing pipeline for large polarization data

We developed a postprocessing pipeline that can reconstruct the registered large datasets imaged with pol-diSPIM, applied to all experimental data in this paper. Such datasets themselves may only occupy a few to tens of gigabytes, which is only a moderate burden for intensity deconvolution, but requires dozens of times more memory and runtime to support spatio-angular reconstruction. For example, a ∼2.2 GB dataset (300×300×300 voxels × 42 polarization modulations, 16-bit float) corresponds to a ∼127 GB dipole OTF matrix (300×300×300 voxels × 42 polarization modulations × 15 spherical harmonics coefficients, 64-bit complex), with a peak memory usage of ∼283GB and a ∼120 s time cost per iteration.

First, cropping for reconstruction. The raw data (assuming the shape is L×W×H voxels × P polarization modulations) is cropped sequentially along the x, y, z axes to a series of sub-volumes and then sent to the GPU-based reconstruction module. The transfer of data between the CPU and GPU memory is time-consuming. To avoid excessive transfer when possible, the shape of each sub-volume should be carefully preset. For our 24 GB memory GPU device and P=42, we set it to 100×100×100×42 with a 10% redundancy for each boundary, ensuring that any single block cost just ∼0.28 s per iteration accelerated by GPU (previously ∼10.1 s per iteration by 128-thread parallel computing).

Then, stitching for fusion. The reconstructed blocks (each one has a 100×100×100 voxels ×15 spherical harmonics coefficients) are placed back to the original position. They are fused by performing linear blending between the overlapping regions. We create a weight mask, multiply the blocks by the weight mask, and then sum the resulting weighted blocks together. The fusion process ensures that there are no obvious boundaries or edge artifacts at the stitching edges.

The 300×300×300×42 data (cropped into 48 sub-volumes) will just require a ∼2 min runtime for 10 iterations (with another ∼8.5s for stitching), in comparison to the previous ∼80 minutes. This pipeline can also be used as a plugin in other preprocessing (cropping, registration) frameworks, e.g., our earlier pipeline for large cleared-tissue data imaged with diSPIM^58^.

### Spatio-angular analysis methods

#### Subset maps of ODFs

Assume that we have a spatio-angular distribution *F*_*l*,*m*_(**r**) or *f*(**r, ŝ**), then density map is an orientation-independent spatial statistic, giving the estimated number of fluorophores at each spatial point:

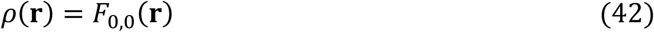

Peak orientation maps represent the most probable direction for fluorophores in each voxel:

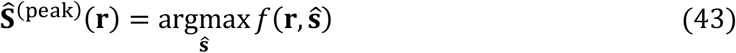

Principal orientation is the maximum projection direction of ODF, comprehensively considering all the local peaks and demonstrating the overall alignment of ODF. The principal orientation map is given by:

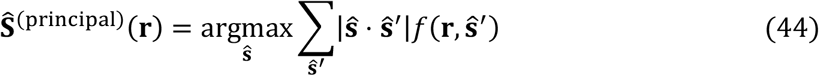

Generalized fractional anisotropy is a biological metric for one single ODF, indicating how oriented the dipoles are. The GFA map of the sample can be computed as:

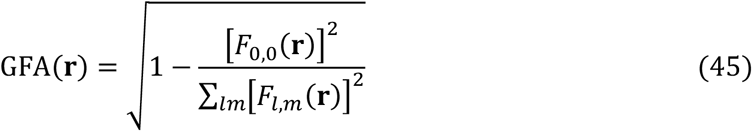

Furthermore, Order parameter is how oriented dipoles are with respect to a special direction 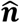. The entire OP map can be computed as:

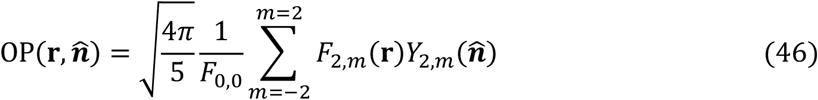

#### Quantitative analysis

For all simulation datasets, we take the whole density map to evaluate the SSIM on eGRL outputs and ground truths with a Python package (scikit-image, v0.18.3).

Structural similarity index measure (SSIM) for density images is defined as:

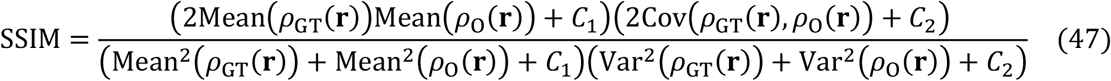

where GT and O are the density maps of ground truth and eGRL output; and *C*_1_ and *C*_2_ are small constants that prevent the denominator from becoming zero.

Orientation-distribution-function similarity index measure (OSIM) is a new index we define to measure the average spatial difference of the shape of ODFs over the entire volume, where **r** and **ŝ** are the discrete spatial points and orientations:

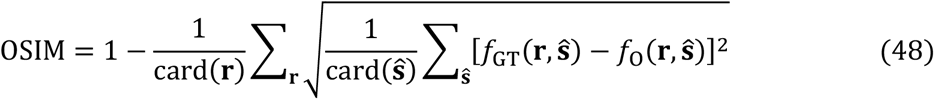

where *f*_GT_ and *f*_O_ are the spatio-angular distribution of ground truth and eGRL output; card(**r**) and card(**ŝ**) are the number of elements in **r** and **ŝ**.

Orientation-distribution-function normalized cross-correlation (ONCC) is a new index we define to measure the spatial-average similarity on the shape of ODFs computed over the entire volume:

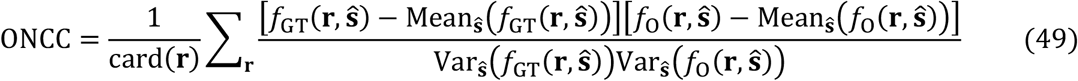

Peak-orientation similarity index measure (PSIM) is a new index we define to measure the spatial-average difference of the direction of peak orientations over the entire volume:

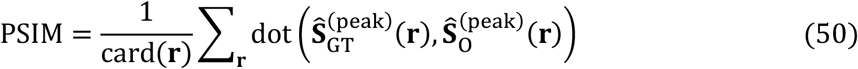

#### Simulation of phantom objects

To evaluate the quality and performance of eGRL (relative to other reconstruction algorithms), we generated four types of 3D phantoms with well-designed angular distributions in Python.

Single shell phantom in a 64×64×64 volume. The initial shell has a 1-voxel thickness, 30-voxel radius, and uniform density distribution, which is then blurred by a Gaussian filter with sigma=2 for smoothing the boundary. ODFs are attached to each voxel with the same GFA but varying dipole moments parallel to the radial direction.

Multiple solid spheres are randomly located in a 128×128×128 volume. The spheres were generated with random central density (500-900 counts, decaying under a Gaussian function from center to edge) and random radius (2-6 voxels). Peak orientations at each voxel are parallel to the radial direction of its sphere.

Three single-helix phantoms in a 64×22×22 or a 64×64×64 volume, sampled at 130nm isotropic voxel resolution. They rotated along the x, y, z axes separately, but all have a 20×20×20 bounding box and are placed equidistant along the x axis. Each of them has a uniform density (with 600nm radius and 1000nm pitch) but a monotonically varying polarization distribution. Each dipole moment is parallel to the tangential direction of the helix.

Two double-helixes phantoms in a 128×128×128 volume, sampled with 130 nm isotropic voxels. Inside each phantom, two helices are arranged like a DNA structure with a specific relative distance between them. Each of them has a uniform density (with 600 nm radius and 1000 nm pitch) but varying polarization distribution (GFA increases from bottom to top). The two phantoms separately have a 554 nm and 478 nm inner space, 7020 nm and 7280 nm pitch, and the same 600 nm radius. Also, each dipole moment is parallel to the tangential direction of the helix.

Typically noise-free data *S* were synthetically generated from phantoms and then corrupted with Poisson noise *N*_*p*_. The SNR (dB) of simulated noisy raw data can be computed as:

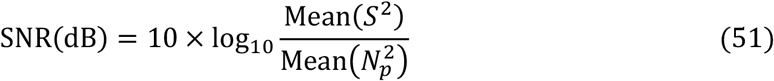

where Mean(. ) is used to compute the mean value of the volumes.

#### Mimic polarized beads dataset based on non-polarized beads images

100 nm yellow green beads were captured using an asymmetric diSPIM with a pair of 1.1NA and 0.67NA objectives with no polarization modulation, noted as bead_*v*_(**r**), with *v* indicating each view.

In calibration, the average response of fluorescent lake (uniform intensity, isotropy ODF) to polarization state *p and view v* is calculated to 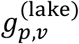. We assume that fluorophores in the beads are randomly oriented, so there’s merely a global intensity change between the images under polarized and non-polarized illumination. Hence, we scaled the overall intensity of non-polarized beads images by the factor calculated from calibration fluorescent lake, to obtain synthetic beads images that would mimic the measurement from polarization *p* and view *v*:

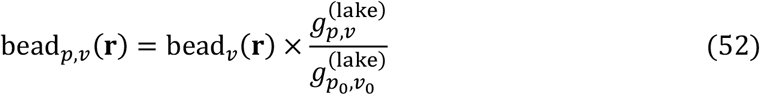

### Sample preparation

#### Bead samples

Glass coverslips (24 mm × 60 mm, #1.5, Electron Microscopy Sciences, 63793-01) were cleaned with clean water and coated with 0.1% poly(l-lysine) (Sigma-Aldrich, P8920) for 10 minutes. 100 nm diameter yellow-green beads (Thermo Fisher Scientific, F8803) were diluted ∼ 10^5^ -fold, and 20 µL added to the coverslip. After 10 minutes, the coverslip was washed three times with clean water before imaging. Beads were used to obtain measured estimates of the system PSF, which in turn were used to guide the generation of theoretical PSFs.

#### Giant unilamellar vesicles samples

We prepared giant unilamellar vesicles (GUVs) via electroformation^59,60^. We coated a coverslip with 20 µL cBSA, waited for ∼ 15 minutes at room temperature for it to dry into a thin layer, then washed three times with distilled water. We mixed 2 µL of FM1-43 (ThermoFisher, a membrane crossing dye with a dipole transition moment oriented normal to the membrane^61^) and 40 µL of GUV solution in a 1.5 mL tube, transferred the solution to the cBSA coated coverslip, and waited for ∼20 minutes for GUVs to settle. Finally, we placed the coverslip in the imaging chamber, filled it with sucrose solution, and waited ∼ 12 hours, covered with a thin film to reduce evaporation, before imaging.

#### Fixed plant xylem samples

Xylem cells were prepared by inducing tobacco (Nicotiana tabacum) BY-2 cells to differentiate into tracheary elements, as described by Yamaguchi et al.^62^. Briefly, cells were cultured with standard methods for BY-2^63^. A stable cell line was generated in which a transcription factor (VND7), driven by an inducible promoter (dexamethasone), had been integrated into the genome. Four days after adding 1 µM dexamethasone to the culture, cells were collected, stained for 30 minutes with 0.02% fast scarlet in growth medium, rinsed in growth medium, adhered to poly-L-lysine coated coverslips, and air dried. Fast scarlet binds cellulose in an oriented manner^64^.

#### Fixed U2OS cells with labelled actin

U2OS cells (American Type Culture Collection, HTB-96) were cultured in DMEM media (Lonza, 12-604F) supplemented with 10% FBS (Thermo Fisher Scientific, A4766801) at 37^°^C and 5% CO2 on coverslips. Cells were fixed by 2% paraformaldehyde (Electron Microscopy Sciences, 15711) in 1× PBS at room temperature for 15 minutes and rinsed three times with 1× PBS. Cells were incubated with Alexa Fluor 488 phalloidin (Invitrogen, A12379, 1:50 dilution in 1× PBS) for 1 hour at room temperature and rinsed three times with 1× PBS before imaging.

#### Fiber network manufacturing

Polystyrene fibers were manufactured using the non-electrospinning spinneret-based tunable engineered parameters (STEP) platform as previously reported^65,66^. Polystyrene of two different molecular weights (Agilent, Mw = 15 × 10^6^ g/mol and Polystyrene Standard, Mw = 2.5 × 10^6^ g/mol) was dissolved in xylene (Carolina Chemicals) to form polymeric solutions at 5% (w/w). Additionally, 20 µL of 1 mg/mL of BDP FL Maleimide dye (Lumiprobe) was added to the polymer solutions to get fluorescent fibers. Fibers were spun on hollow 5 × 5 mm metal scaffolds. The first layer of fibers deposited were large diameter fibers ∼ 2 µm (Mw = 15 × 10^6^ g/mol) followed by an orthogonal layer of 200 nm (Mw = 2.5 × 10^6^ g/mol) fibers with spacing varying from 7 to 20 µm to achieve a variety of cell shapes (elongated on single fibers and parallel-shaped cells on two or more fibers)^67-69^. Additionally, crosshatch networks of 200 nm fiber diameters were also prepared with spacing varying from 7 to 20 µm^70,71^ to achieve polygonal and kiteshaped cells on multiple fibers. The fiber networks were fused at junctions using a custom-built chemical fusing chamber.

#### Cell culture and seeding on fiber networks

3T3 mouse fibroblasts (ATCC) were grown in Dulbecco’s modified Eagle’s medium (Corning) supplemented with 10% fetal bovine serum (Corning) in T25 flasks (Thermo Scientific). The cells were grown in an incubator kept at 37°C and 5% CO2. The nanofiber network scaffolds were tacked on a cover glass (VWR, 24 × 60 mm No. 1.5) with the help of high-vacuum grease (Dow Corning). Next, the scaffolds were sterilized with 70% ethanol for 10 minutes followed by Phosphate Buffer Solution (PBS) washes (two times). Next, the scaffold was coated with 4 µg/mL bovine fibronectin (Sigma Aldrich) in PBS for at least one hour to promote cell adhesion. Cells were then seeded onto the scaffolds with a seeding density of 300,000 cells/mL and were allowed to spread onto the fibers for a few hours followed by the addition of 3 mL of media. Cells were allowed to further spread for an additional 24 hours before fixation.

#### Immunostaining cells on fiber networks

NIH3T3 cells were fixed with 4% paraformaldehyde in PBS (Santa Cruz Chemicals) for 15 minutes. The cells were then washed with PBS twice and then permeabilized with 0.1% Triton X-100 solution. Following two PBS washes, the cells were blocked with 5% goat serum (Fisher Scientific) for 30 minutes. Next, conjugated antibody Alexa Fluor 568 Phalloidin (1:100, Thermo Fisher) diluted in antibody dilution buffer was added to the cells. After one hour, the cells were washed with PBS (3×, 5 minutes each). The sample was then covered in 2 mL of Live Cell Imaging Media (Thermo Fisher) for imaging.

#### Fixed HeLa cells with labeled actin

Hela cells (American Type Culture Collection, HTB-96) were cultured in DMEM media (Lonza, 12-604F) supplemented with 10% FBS (Thermo Fisher Scientific, A4766801) at 37 °C and 5% CO2 on coverslips. Cells were fixed by 2% paraformaldehyde (Electron Microscopy Sciences, 15711) in 1× PBS at room temperature for 15 minutes, permeabilized with 0.2% Triton-X 100 (Sigma, 648463) in 1x PBS for 5 minutes and rinsed three times with 1× PBS. Cells were incubated with Alexa Fluor 488 phalloidin (Invitrogen, A12379, 1:50 dilution in 1× PBS) for 1 hour at room temperature and rinsed three times with 1× PBS before imaging in 1X live imaging solution (Thermofisher, A59688DJ).

#### Live HeLa cells with labeled actin

Live Hela cells (American Type Culture Collection, HTB-96) were incubated with DMEM medium (Lonza, 12-604F) containing 10% FBS (Thermo Fisher Scientific, A4766801) and 500nM of SiR-Actin (Cytoskeleton, CY-SC001) for 2 hours at 37 °C with 5% CO2. After labeling, cells were incubated in 1x live cell imaging solution for imaging.

#### Live Macrophage cells with labeled membrane

RAW264.7 mouse macrophages were maintained in RPMI 1640 (ATCC modification, Gibco, A1049101) with 10 % heat-inactivated FBS (R&D Systems, S12550H) at 37°C with 5% CO2. For imaging, cells were stimulated with 100 ng/ml PMA (Sigma, P1585) in RPMI without phenol red (ThermoFisher, 11835030) supplemented with 0.5 mM ascorbic acid (Sigma, A4544) to minimize photoxicity due to increased oxidative stress. Cells were labeled with 5 mM FM1-43 at room temperature for 2 minutes and rinsed with imaging media and immediately imaged in imaging media.

#### Live U2OS cells with labeled membrane

Live U2OS cells expressing RFP-PH were incubated with pre-chilled Hank’s balanced salt solution (HBSS) containing 5 µg/mL FM1-43 (Thermofisher, T3163) on ice for 2 minutes and rinsed by 1x live cell imaging solution (Thermofisher, A59688DJ) 3 times. Cells were incubated in the 1x live cell imaging solution for imaging.

## Data and code availability

The source code of this manuscript is available at https: https://github.com/ejunyuliu/PFM-tool. The data that support the findings of this study are included in **Figs. 1-5, Supplementary Figs. 1–23** and **Supplementary Videos 1–16**. Representative data to test the code are publicly available at https: https://doi.org/10.5281/zenodo.17033756; other datasets are available from the corresponding author upon request.

## Acknowledgments

The instrumentation and data acquisition were performed at the National Institute of Biomedical Imaging and Bioengineering, within the intramural program of the US National Institutes of Health. We thank Abhishek Kumar and Rudolf Oldenbourg for their assistance in building the pol-diSPIM system, and Harshad Vishwasrao for his assistance in maintaining the pol-diSPIM system. This research was supported by the National Natural Science Foundation of China (62427807, 62475232, 62405269, 6240030418), the Talent Program of Zhejiang Province (2021R51004), the Leading Innovative and Entrepreneur Team Introduction Program of Zhejiang (2024R01001), the Postdoctoral Fellowship Program of CPSF (GZB20230626), and the Howard Hughes Medical Institute (HHMI). This article is subject to HHMI’s Open Access to Publications policy. HHMI laboratory heads have previously granted a non-exclusive CC BY 4.0 license to the public and a sub-licensable license to HHMI in their research articles. Pursuant to those licenses, the author-accepted manuscript of this article can be made freely available under a CC BY 4.0 license immediately upon publication.

## Author contributions

Conceived project: H.S., H.L., M.G. Implemented algorithms: J.L., T.C, P.L-R., H.L., M.G. Wrote software: J.L., T.C, M.W, Y.L., M.G. Designed experiments: J.L., T.C, Y.W., T.I.B., R.F., V.S., A.N., S.M., P.L-R., H.S., H.L., M.G. Prepared samples: A.A., Y.S., J.C., T.I.B., V.J., R.F., M.G. Performed experiments: J.L., A.A., Y.S., Y.W., J.C., R.F., M.G. Performed data analysis: J.L., T.C, Y.L., M.W., P.X., H.Y., W.Z., M.G. Wrote manuscript: J.L., H.L. and M.G., with advice from all authors. Provided biological insight and advice: A.A., T.I.B., R.F., V.S., A.N., S.M. Supervised research: V.S., A.N., S.M., P.L-R., H.S., H.L., M.G.

