## Supplementary File for "Observing biological spatio-angular structures and dynamics with statistical image reconstruction and polarized fluorescence microscopy"

### Supplementary Video Captions

**Supplementary Video 1, Rendering methods for clearer visualization of spatio-angular objects.** First, we synthesize a spatio-angular shell phantom consisting of dumbbell shaped ODFs oriented along the radial direction at each surface voxel. Then, volumetric (left column), sliced (middle column) and fly-around (right column) ODF/peak orientation maps are rendered under different thresholds and down-sampling ratios, where the orientations in the peak orientation maps are represented by both the directions and the pseudo colors of cylinders. See also **Supplementary Fig. 1a**.

**Supplementary Video 2, Demonstration of spatio-angular reconstruction process on synthetic phantom with eGRL.** Two parallel lines in 3D space with two different kinds of ODFs in each line (1$^{st}$ column, dumbbell-shaped in top line and x-shaped in bottom line) are imaged with polarized diSPIM. Here only a transverse XY plane through the object is shown under one specific polarization modulation. The spatial and angular distribution are jointly reconstructed by eGRL, ODFs in the two lines (2$^{nd}$ column) and line profiles through the middle of the image (3$^{rd}$ and 4$^{th}$ column) are also shown and updated after each iteration count. See also **Fig. 1b**.

**Supplementary Video 3, Performance of eGRL vs. SVD on synthetic data with different noise levels.** A synthetic shell phantom is imaged with polarized dual-view light sheet microscopy and the raw data are corrupted with different levels of Poisson noise, then separately reconstructed by eGRL (10 iterations) and SVD (0.01 regularization parameter). Note eGRL reconstruction is robust to noise while SVD is quite sensitive to noise. See also **Supplementary Fig. 8**.

**Supplementary Video 4, Bias errors caused by regularization parameter in SVD reconstruction on simulated data.** Synthetic shell phantom with molecular orientations perpendicular to the surface (left). SVD reconstructions at different regularization parameters $\eta$ ('eta') values are visualized in the peak orientations (middle) and the radar chart (right) counts the x, y and z proportion of all peak orientations. Increasing the regularization parameter results in structural distortion and orientations biased towards the y-axis. See also **Fig. 2g**.

**Supplementary Video 5, Experimental GUV labelled with FM1-43, showing raw data and reconstruction process.** Raw images with 18 polarization modulations (9 volumes for each view) were acquired with pol-diSPIM (left column), highlighting local differences in intensity in different subregions of the same structure. Taking the average of all raw data as the initial estimate, eGRL iterations jointly estimate the peak orientation (middle column) and density (right column) distributions. All the density images are shown from the same middle slice (z = 10.5 µm) selected from the 3D volume. See also **Fig. 3a**.

**Supplementary Video 6, Characterizing orientational preference of FM1-43 anchored to GUV**

**membrane.** As in **Supplementary Video 5**, but shown in slices. Each layer along the y-axis is visualized by slicing (left column), showing the ODF (middle column) and peak (right column) orientations aligned perpendicular to the shell. See also **Fig. 3a**, **b**.

**Supplementary Video 7, Fly-around demonstration of orientation distribution in GUV samples.** As in **Supplementary Video 5**, but showing the reconstructions of the whole GUV dataset ($1^{st}$ column) by eGRL vs. SVD estimation, and 3 representative ROIs ($2^{nd}$ – $4^{th}$ column) for higher magnification observation. A color-coded sphere is visualized at the corner and synchronously rotated to indicate which orientation a specific color represents. See also **Fig. 3a**, **b**.

**Supplementary Video 8, eGRL provides better structural contrast and more accurate orientation distribution than SVD when applied to fixed U2OS samples labelled with Alexa Fluor 488 phalloidin.** eGRL and SVD results are shown in terms of density (left column), ODF (middle column) and peak orientation (right column) maps. See also **Fig. 3c-e**.

**Supplementary Video 9, eGRL still performs well with reduced polarization measurements on tobacco xylem cells labelled with Pontamine fast scarlet.** Raw images were captured with 18, 8 and 6 polarization modulations and separately reconstructed by both eGRL and SVD. Note the downsampling in polarization encoding shows that eGRL is robust to the accurate orientation details. See also **Fig. 3h.**

**Supplementary Video 10, eGRL performs well with reduced polarization measurements on fixed U2OS labelled with Alexa Fluor 488 phalloidin**. Raw images were captured on fixed U2OS cells under 18, 8 and 6 polarization modulations and separately reconstructed by both eGRL (top) and SVD (bottom). See also **Supplementary Fig. 13**.

**Supplementary Video 11, Visualizing Xylem cells labeled with Pontamine fast scarlet.** Orientation map of Xylem cell was reconstructed with eGRL and SVD. We also show slices taken along the y axis of the ODF and orientation map. See also **Fig. 3h**.

**Supplementary Video 12, eGRL retrieves angular rotational-distribution and dynamics of SiR Actin in live HeLa cells, that are obscured in intensity images.** Density map (left) and maximum-intensity-projection pseudo-colored by orientation (middle). The polar charts (right) indicate the orientation distribution along the trails on peripheral (yellow border) and protrusion (green border) regions respectively represented by 50 and 25 selected points in the density map. Low fluctuations along peripheral region serve as a control to assess orientation dynamics caused by non-cellular changes, e.g. photobleaching. Dynamics along protrusion demonstrate small changes that are obscured in the intensity images, but noticeable in the orientation distribution. See also **Fig. 5a-c**.

**Supplementary Video 13, Membrane transitions, as assessed via morphology and orientation, in a single live macrophage cell labeled with FM1-43.** Time-lapse maximum intensity projection of density (left) and orientation distribution (middle), revealing the transition from membrane protrusion to expanding bleb (magnified in right column). See also **Supplementary Fig. 19**.

**Supplementary Video 14, eGRL restores spatio-angular dynamics on membrane protrusions in living macrophages.** First, fly-around density and orientation maps at initial time point to characterize the orientational alignment of FM1-43 probe on live macrophage samples. Then, z-slices of dual-view raw data from pol-diSPIM are compared with estimated and orientation distribution. Last, we show time-lapse maximum intensity projections of reconstructed density maps that are pseudo-colored by orientation. See also **Fig. 5d-e.**

**Supplementary Video 15, Dual-color spatio-angular imaging of live U2OS cells labeled by FM1-43 and RFP-PH dyes.** Time-lapse maximum intensity projection of dual-color raw data (1$^{st}$ column), reconstructed orientation (2$^{nd}$ and 3$^{rd}$ columns) and GFA (4$^{th}$ and 5$^{th}$ columns) maps in FM1-43 and RFP-PH channels. Note GFA extracted from spatio-angular reconstructions by eGRL provides a quantitative measure of molecular distribution anisotropy beyond principal-orientation representations. See also **Supplementary Fig. 21.**

**Supplementary Video 16, eGRL reconstructs spatio-angular dynamics on live U2OS cell membrane labeled by FM1-43.** First, successive image planes of pol-diSPIM raw data and eGRL reconstructed spatio-angular distribution at time point 0. Then, rendering of time-lapse volumetric density and orientation map, showing a higher magnification view of the cell substrate to highlight the orientational alignment that accompanies the membrane expansion. See also **Fig. 5f-k**.

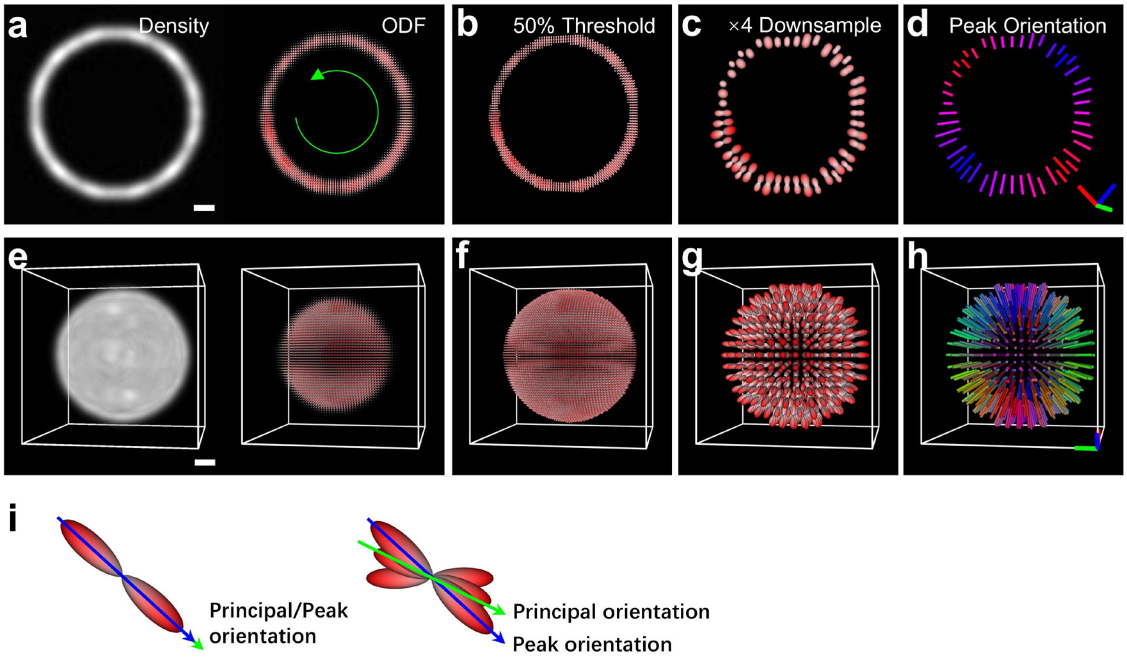

**Supplementary Fig. 1, Different representations of the spatio-angular distribution. a)** the density and the corresponding ODF map of a ring phantom with continuously decreasing polarization (i.e., decreasing Generalized Fractional Anisotropy (GFA, **Methods**) and increasing isotropy) along the orientation described by the green circle. Note the density distribution (**a**, left) is not uniform along the ring and the ODFs are densely displayed to show this density fluctuation. **b)** ODF map with low intensity removed to facilitate display (Threshold = 0.5*maximum density). **c)** and additional 4 times spatial downsampling on the image in **b** to further sparsen ODFs for clearer display. **d)** the corresponding peak orientation map, in which the oriented cylinders marked by different colors demonstrate the most probable direction (ODF has the largest value in this direction) for fluorophores, and the length of cylinders indicates the degree of polarization. **e)** a shell phantom with uniform orientation (ODFs at all voxels are oriented in the radial direction) and density distributions. **f-h)** same as **b-d** but corresponding to the shell phantom in **e**. **i)** Principal orientation vs. Peak orientation on the ODFs with single/multiple local peaks. The principal orientation is the maximum projection direction of ODF (also the direction with maximal Order Parameter), comprehensively considering all the local peaks and demonstrating the overall alignment of ODF. It is more robust to noise than the peak orientation. Scale bars: **a-h** 1 µm.

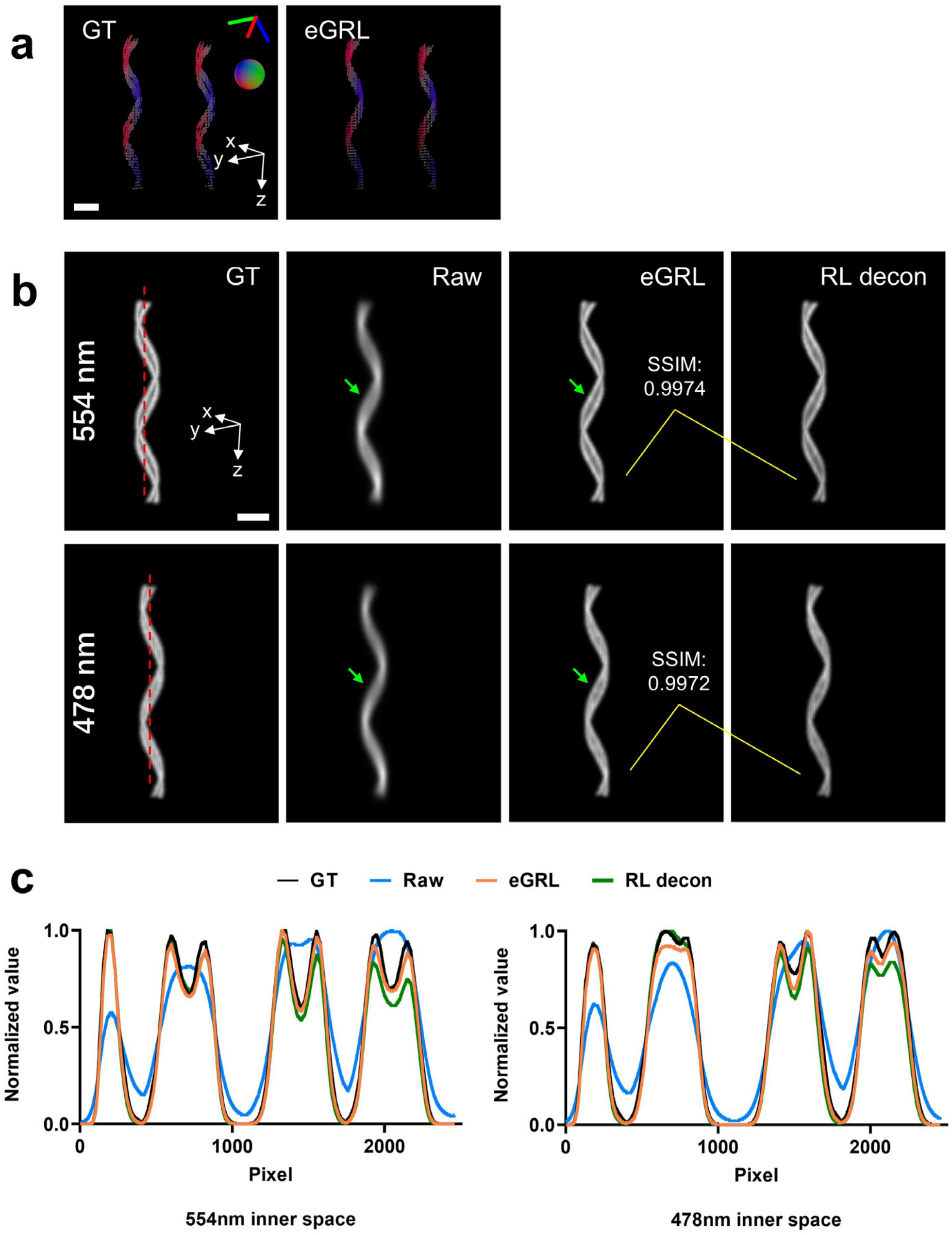

**Supplementary Fig. 2, eGRL produces high fidelity estimates of oriented phantom structures. a)** Orientation distributions of double-helix phantoms and reconstructions produced by eGRL (see also **Fig. 1e**). **b)** Density maps of the double-helix phantoms (from left to right): ground truth (GT), raw data (average of all polarization measurements), the eGRL reconstruction (20 iterations), and the deconvolved image (RL deconvolution from the averaged raw data, 20 iterations). **c)** Line profiles along the red lines in **b)**. Note eGRL provides similar results as RL deconvolution and can distinguish the closely separated helices (indicated by green arrows) which are hardly resolved in the raw data. Scale bars: **a** 5 µm, **b** 2 µm.

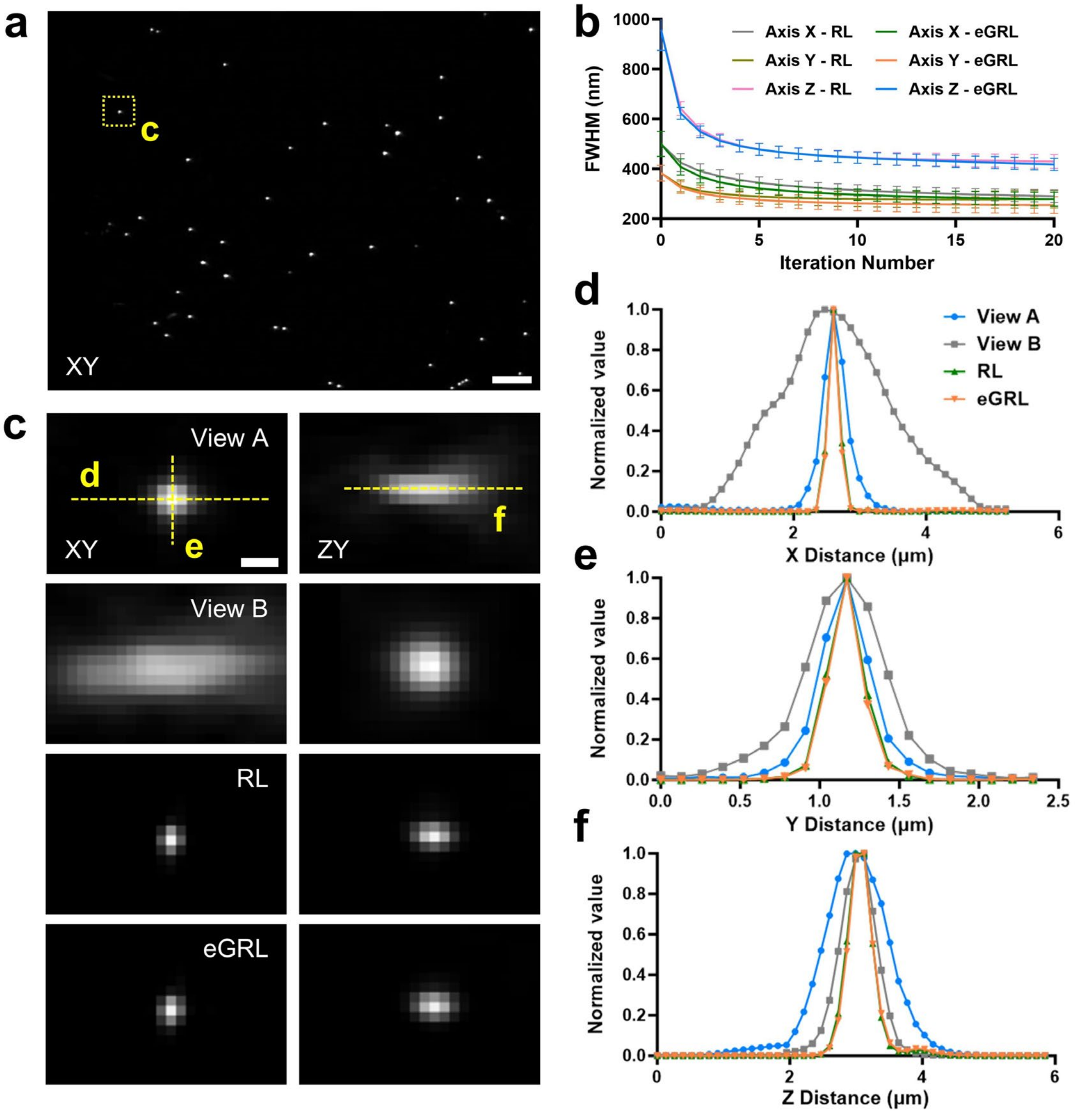

**Supplementary Fig. 3, Comparative evaluation of the deconvolution ability of eGRL and RL using images of 100 nm beads. a)** 100 nm yellow green beads were captured using an asymmetric diSPIM with a pair of 1.1NA and 0.67NA objectives (with no polarization modulation). Maximum-intensity projection bead images are shown. If the fluorophores in the beads are randomly oriented (isotropic response to polarization), we would expect merely a global intensity change between the images under polarized and non-polarized illumination. Hence, we scaled the overall intensity of non-polarized beads images to obtain synthetic beads images that would mimic the measurement from 18 polarization modulations (Scheme 1, see **Supplementary Table 2**). eGRL was performed on this synthetic dataset, while the RL result was obtained by joint deconvolution of the two views after averaging all image volumes in each view. **b)** Full width at half maximum (FWHM) values derived from n = 10 beads in **a)**, standard deviations as well as mean values are shown. RL results were obtained by joint deconvolution of the polarization-averaged images from each view. **c)** Image planes from the center of one of the raw polarization volumetric datasets as the outlined region in **a**, showing data from view A and B, RL deconvolution results of polarization-averaged raw data from each view, and eGRL reconstructions outcome when using raw data consisting of all polarization modulations and views. **d-f)** Comparison of axial and lateral line profiles from **c)**. Scale bars: **a** 5 μm, **c** 0.5 μm.

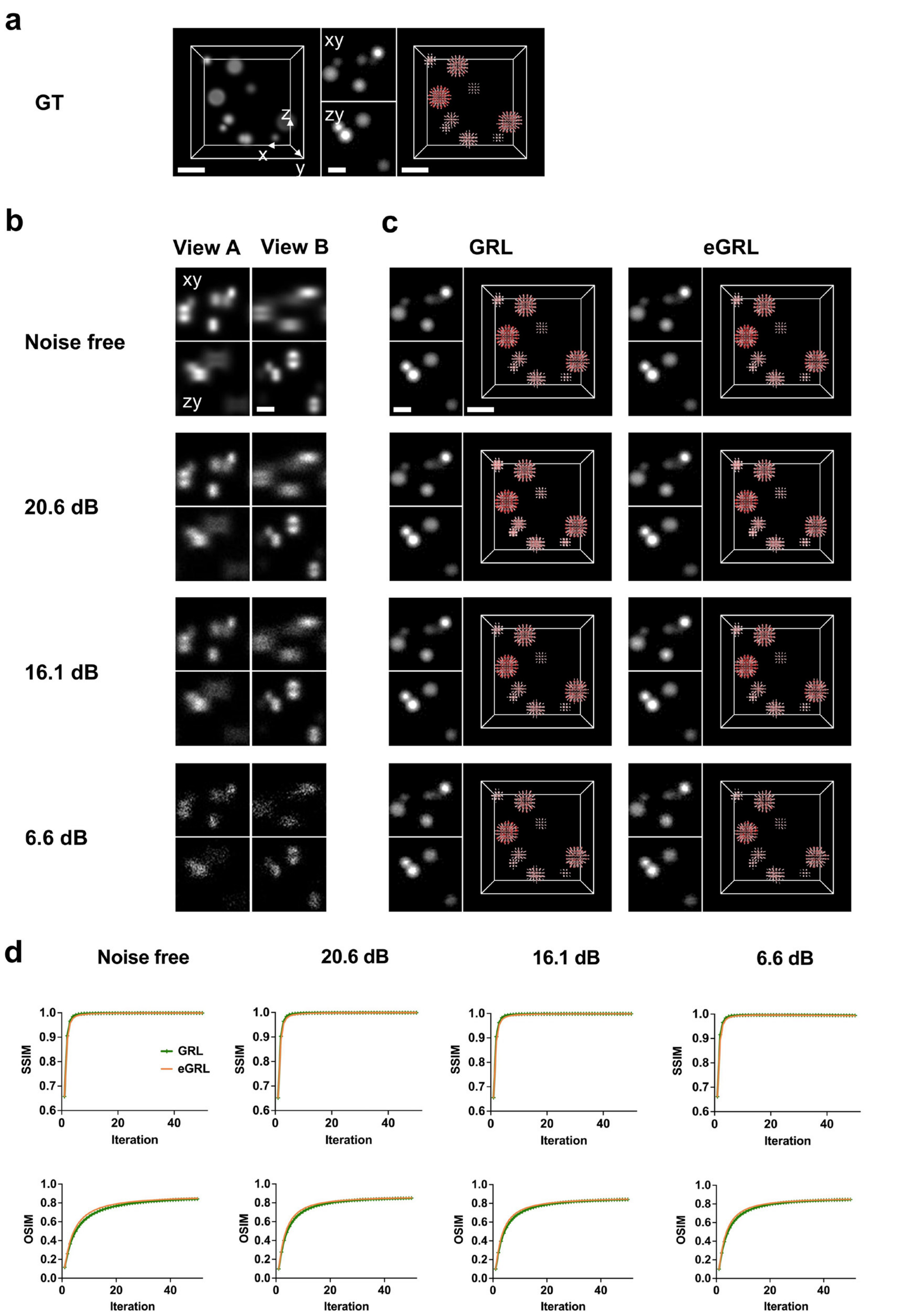

**Supplementary Fig. 4, eGRL and GRL provide similar image quality as assayed on synthetic data. a)** Synthetic spheres with random sizes and positions, showing density maps (left), lateral and axial slices (middle), and the ODF map (right). **b)** Raw data simulated from sample in **a** by pol-diSPIM dipole PSFs with different noise levels, showing the xy (top) and zy (bottom) slices from the two orthogonal views. **c)** The lateral (top left), axial (bottom left) slices, and the ODF maps (right) from the GRL and eGRL results, reconstructed from the raw data in **b** with corresponding SNR. **d)** SSIM and OSIM (ODF similarity index measure. See **Methods**) analysis of the ODF maps between the reconstructions and ground truth. Note extreme similarity between eGRL and GRL. Scale bars: **a-d** 2 μm.

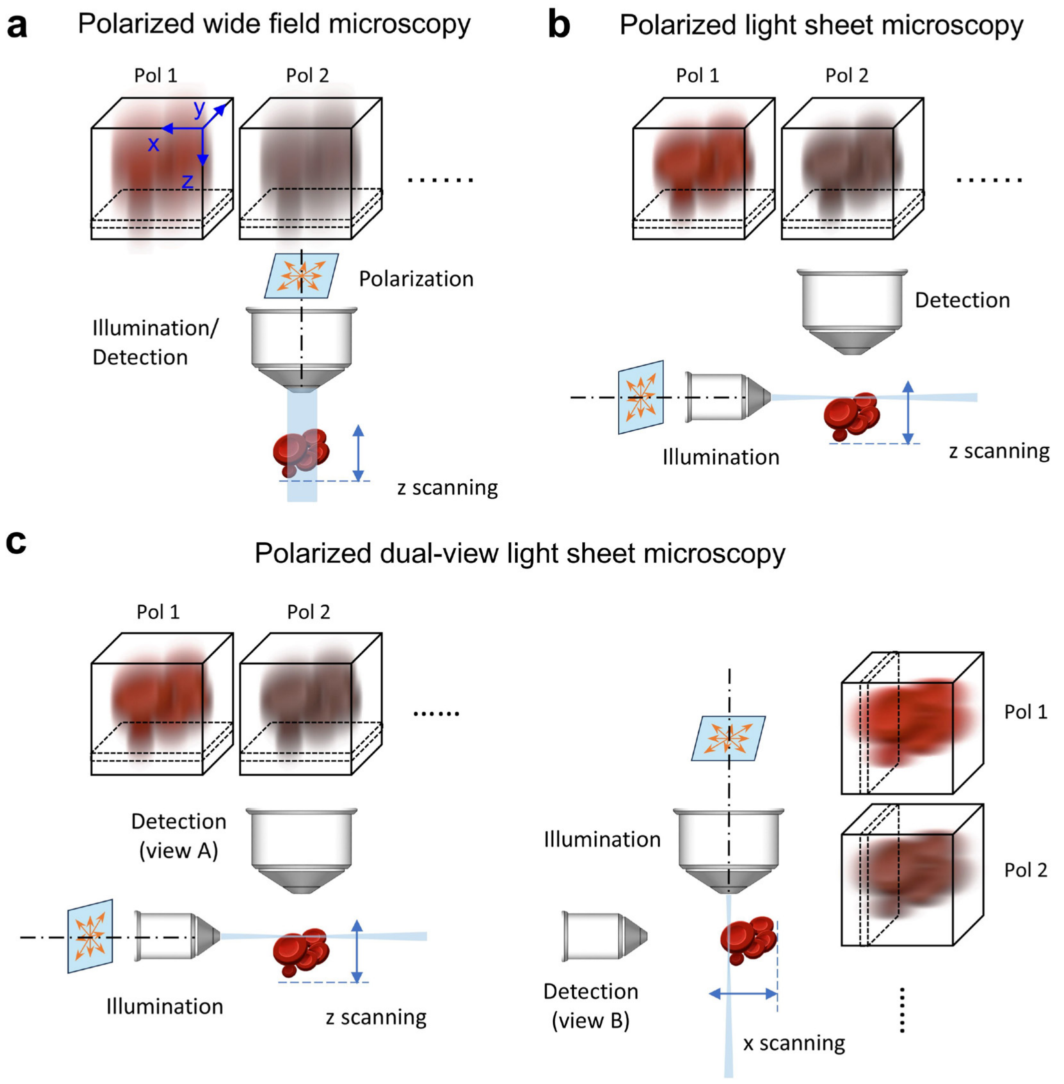

**Supplementary Fig. 5, Schematic of different configurations of polarized fluorescence microscopy used in simulations. a)** Volumetric measurements under varying polarization modulations are modelled based on wide field microscopy. The polarized illumination and corresponding detection are through the same objective. Dashed rectangles in volumes refer to how the slices are stacked during axial scanning. Orange double-headed arrows indicate different polarization directions perpendicular to the illumination axis. **b)** Same as in **a** but configured with light sheet microscopy. Polarized light sheet illumination and corresponding detection are via two orthogonal 1.1NA and 0.67NA objectives. **c)** In polarized dual-view light sheet microscopy, the dual-view imaging process is modelled from the two orthogonal directions by swapping the detection and illumination paths.

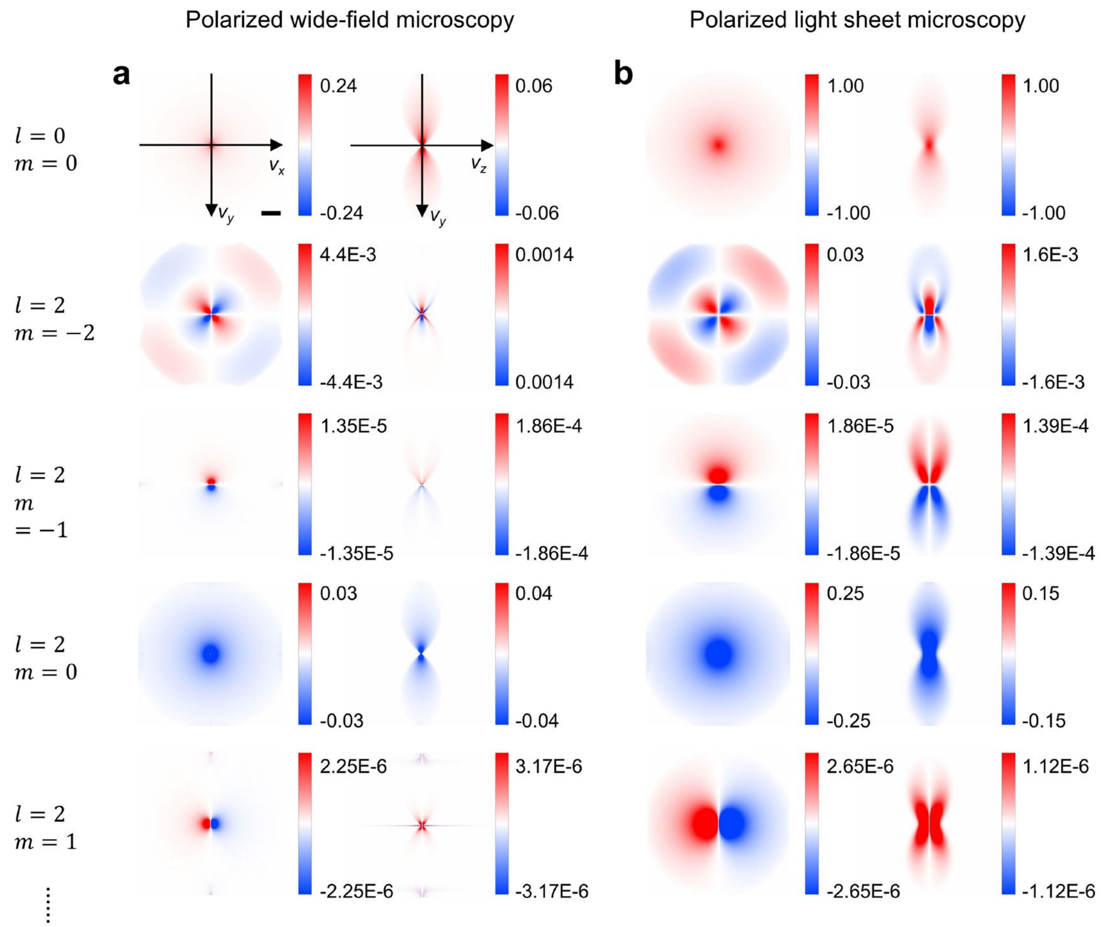

**Supplementary Fig. 6, Dipole OTFs of polarized wide-field microscopy and polarized light sheet microscopy. a)** The first five spherical harmonic orders of the real part of the dipole OTF in polarized wide-field microscopy with objective NA=1.1. **b)** Part of the dipole PSF in a polarized light sheet microscopy with objective NA=1.1 and light sheet waist thickness of FWHM=2000 nm. Note this configuration also can be considered as one of the two views in the polarized diSPIM. Scale bar: **a, b** 0.02 $\mu m^{-1}$.

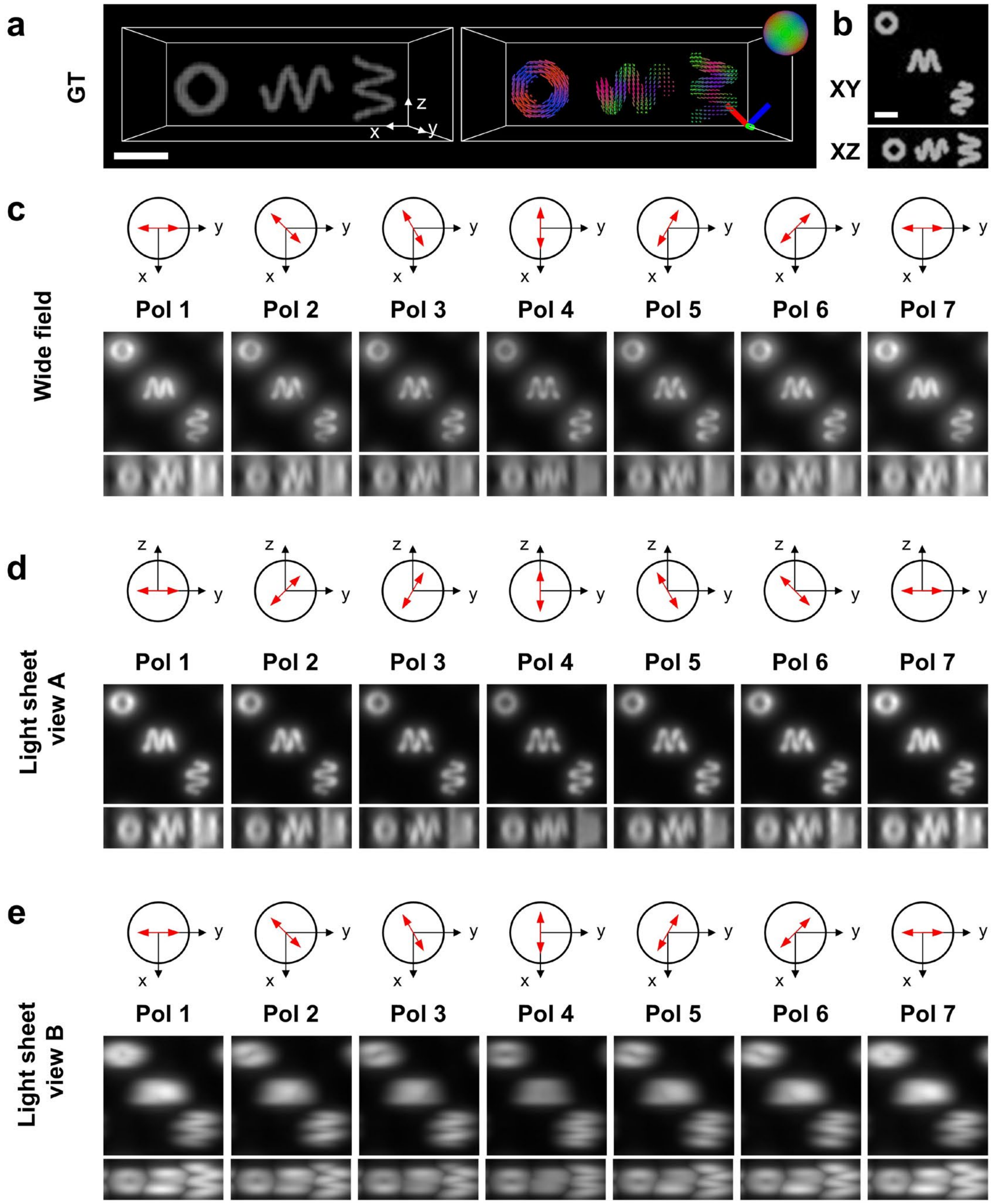

**Supplementary Fig. 7, Synthetic noise-free raw data from polarized wide-field microscopy and pol-diSPIM. a)** Density and orientation distributions of a phantom consisting of three helices (see also **Fig. 2a**). **b)** Maximum-intensity projections of density map of the phantom. **c-e)** Noise-free raw data simulated from spatio-angular imaging process in wide-field microscopy and dual-view light sheet microscopy (diSPIM). Illumination polarization direction and maximum-intensity projections of the raw density image with each polarization modulation are shown. Scale bars: 1 µm.

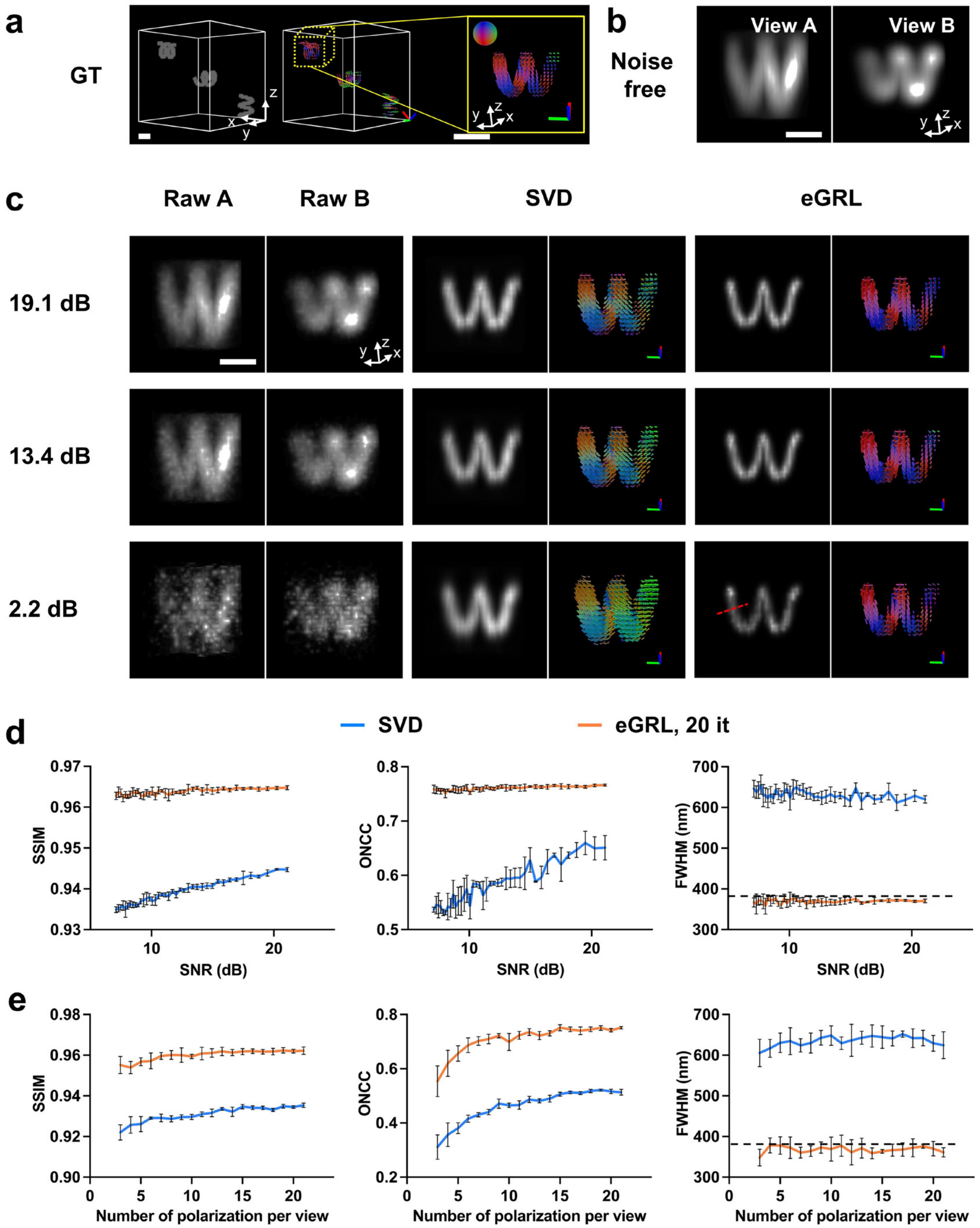

**Supplementary Fig. 8, Performance of eGRL and SVD on synthetic data with different noise levels and different number of polarization modulations. a)** The same three helix phantoms as in **Supplementary Fig. 7** shown in density map (left), peak orientation map (middle), and an additional high magnification view (right) of the yellow dashed rectangle region. **b)** 3D rendering of Y-axis-polarized measurement, which is one of the raw datasets synthetically generated from two orthogonal views in pol-diSPIM, corresponding to the yellow region in **a)**. **c)** Synthetic data in **b** are contaminated with different levels of Poisson noise, and reconstructed by eGRL and SVD algorithms. **d-e)** Evaluation metrics (SSIM, ONCC, and FWHM) vs. input data SNR and number of polarization modulations curves are plotted for the comparison of eGRL and SVD. FWHMs are fitted from profiles along the dashed red line in bottom right **c**. Scale bars: **a** 1 µm, **b, c** 2 µm.

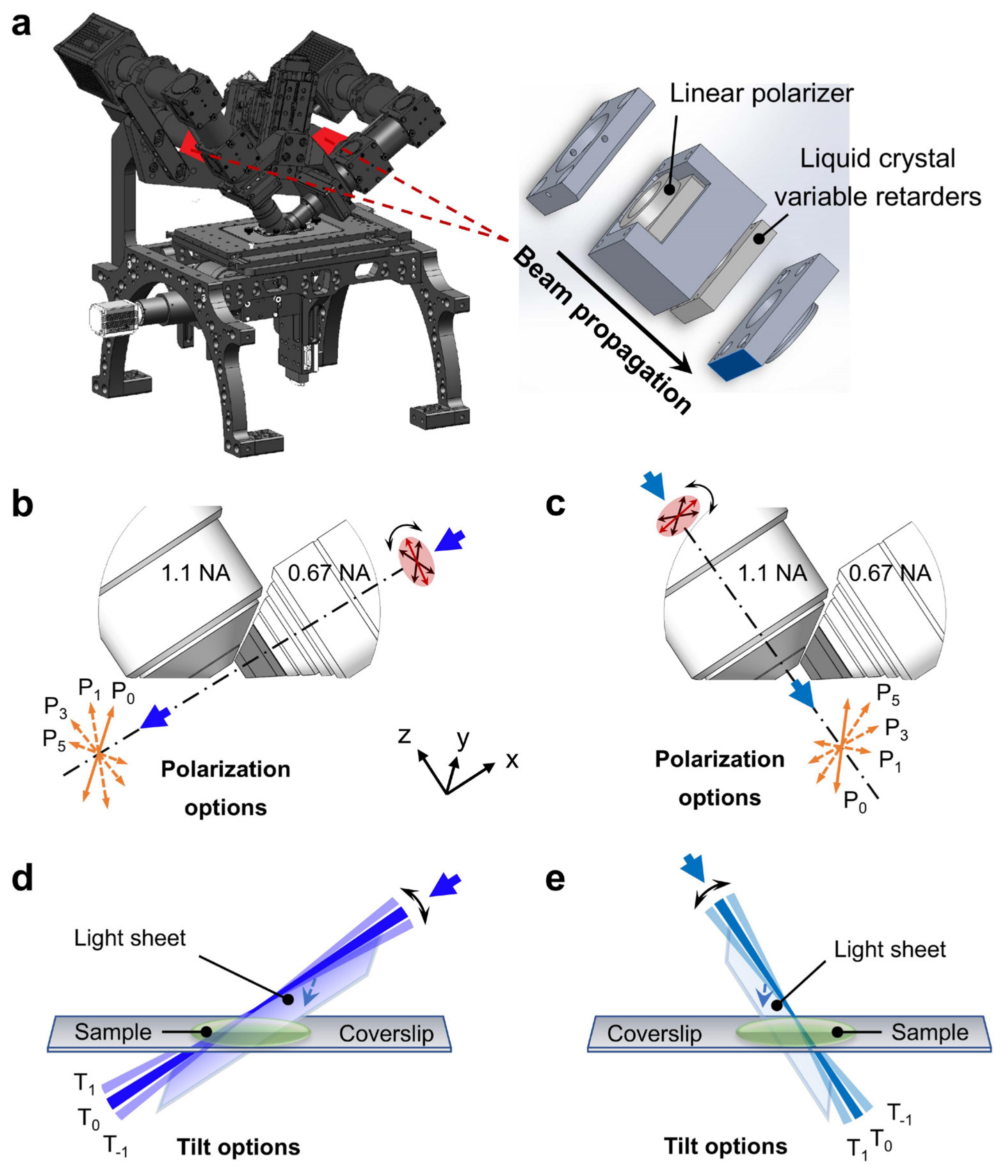

**Supplementary Fig. 9, Instrumentation and imaging schemes for polarized dual-view light sheet microscopy (pol-diSPIM). a)** Overview of the microscope with the liquid crystal (LC) modules (red color in each arm) detailed in right inset. **b)** Polarization modulation for 0.67NA objective illumination and 1.1NA objective detection (View A). A polarized Gaussian beam is selected from a pool of 6 polarization orientations indicated by the orange double-headed arrows (perpendicular to direction of beam propagation) and indexed from 0 to 5, with angles to y-axis 0 (i.e., parallel to the y-axis), 45, 60, 90 (parallel to the z-axis), 120, and 135 degrees. $P_2$ (60 degrees) and $P_4$ (120 degrees) are omitted in the figure for clarity. **c)** Similarly, polarization orientations for 1.1NA objective illumination and 0.67NA objective detection (View B) are also configured with angles to y-axis at 0 (parallel to the y-axis), 45, 60, 90 (parallel to the x-axis), 120, and 135 degrees. **d)** Tilt modulation for 0.67NA objective illumination and 1.1NA objective detection (View A). Three tilt options, $T_1$, $T_0$, and $T_{-1}$, are provided with $T_0$ parallel or close to the low-NA objective optic axis (x-axis). **e)** Similar to **d**, three tilt options for 1.1NA objective illumination and 0.67NA objective detection (View B) are provided with $T_0$ parallel or close to the high-NA objective optic axis (z-axis). The relevant angles are also reported in **Supplementary Table 1**.

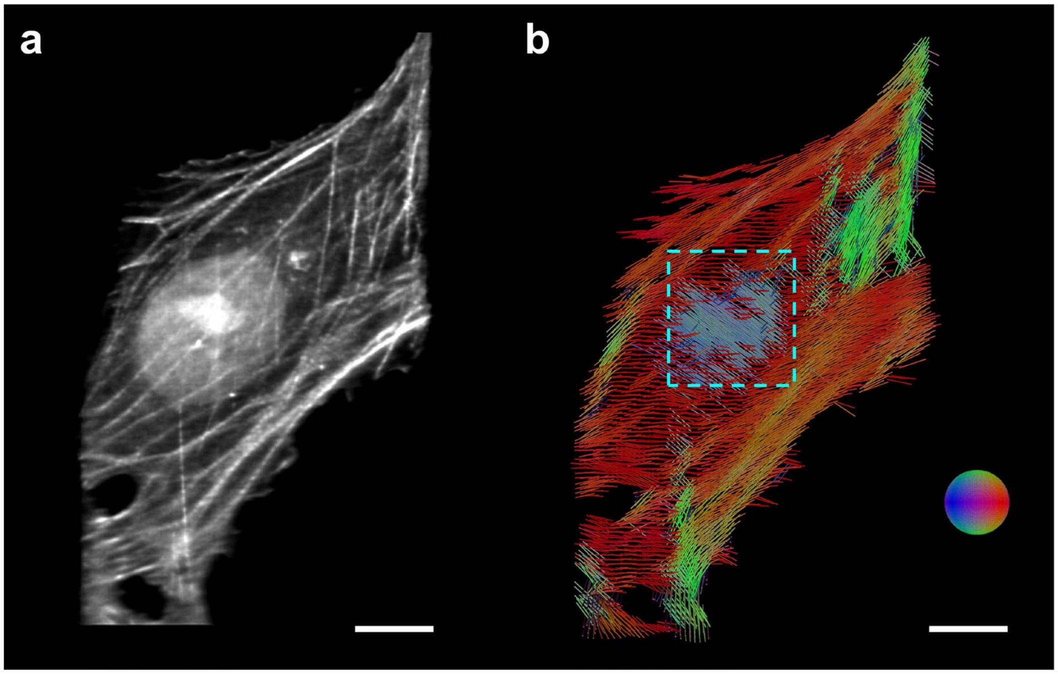

**Supplementary Fig. 10 Differential orientation distribution of fluorophores close to nucleus versus elsewhere in U2OS cell. a)** Density map (shown in average-intensity projection) of U2OS cell labeled with Alexa Fluor 488 phalloidin (see also **Fig. 3c**) reconstructed by eGRL. b) Orientation map shown in averaged projection. The outlined region highlights distinct orientation distribution near the nucleus compared to surrounding areas. Scale bars: **a**, **b** 10 µm.

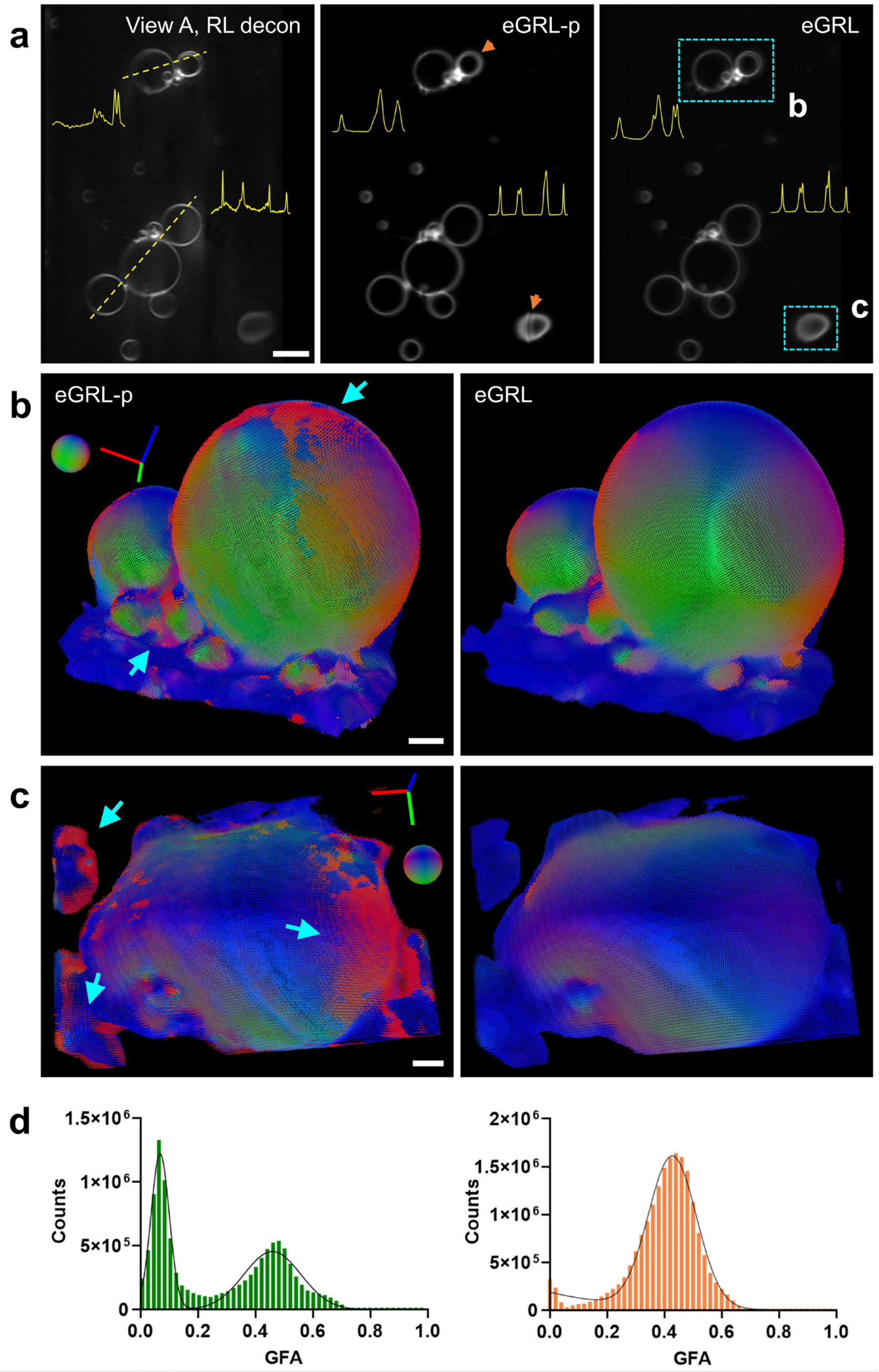

**Supplementary Fig. 11, Performance of eGRL versus its ablated version eGRL-p on GUV samples. a)** Single density slices chosen from reconstruction results of FM1-43 labelled GUVs: RL deconvolution of single view raw image (left), eGRL-p reconstruction based on the RL deconvolved images (middle), and joint angular/spatial eGRL reconstruction (right), with 10 iterations for all cases. The curves at the background delineated the intensity profiles along nearest dashed lines. Orange arrows highlight blurred detail and artifacts in the density map produced by eGRL-p. **b, c)** Peak orientation maps respectively derived from eGRL-p and eGRL, in two ROIs marked by the cyan dashed rectangle regions in **a**. Numerous errors in eGRL-p reconstruction are highlighted by cyan arrows where the anticipated normal-orientations are distorted in the sample. **d)** GFA histograms constructed from the entire GUV dataset. Note that histogram exhibits an additional peak near zero whereas it is typically expected to show a single peak around 0.5, as reported by Chandler, T. et al (2025) [1]. This result indicates that eGRL-p reconstruction errors result in a tendency to lower GFA distributions. Scale bars: **a** 10 µm, **b**, **c** 2 µm.

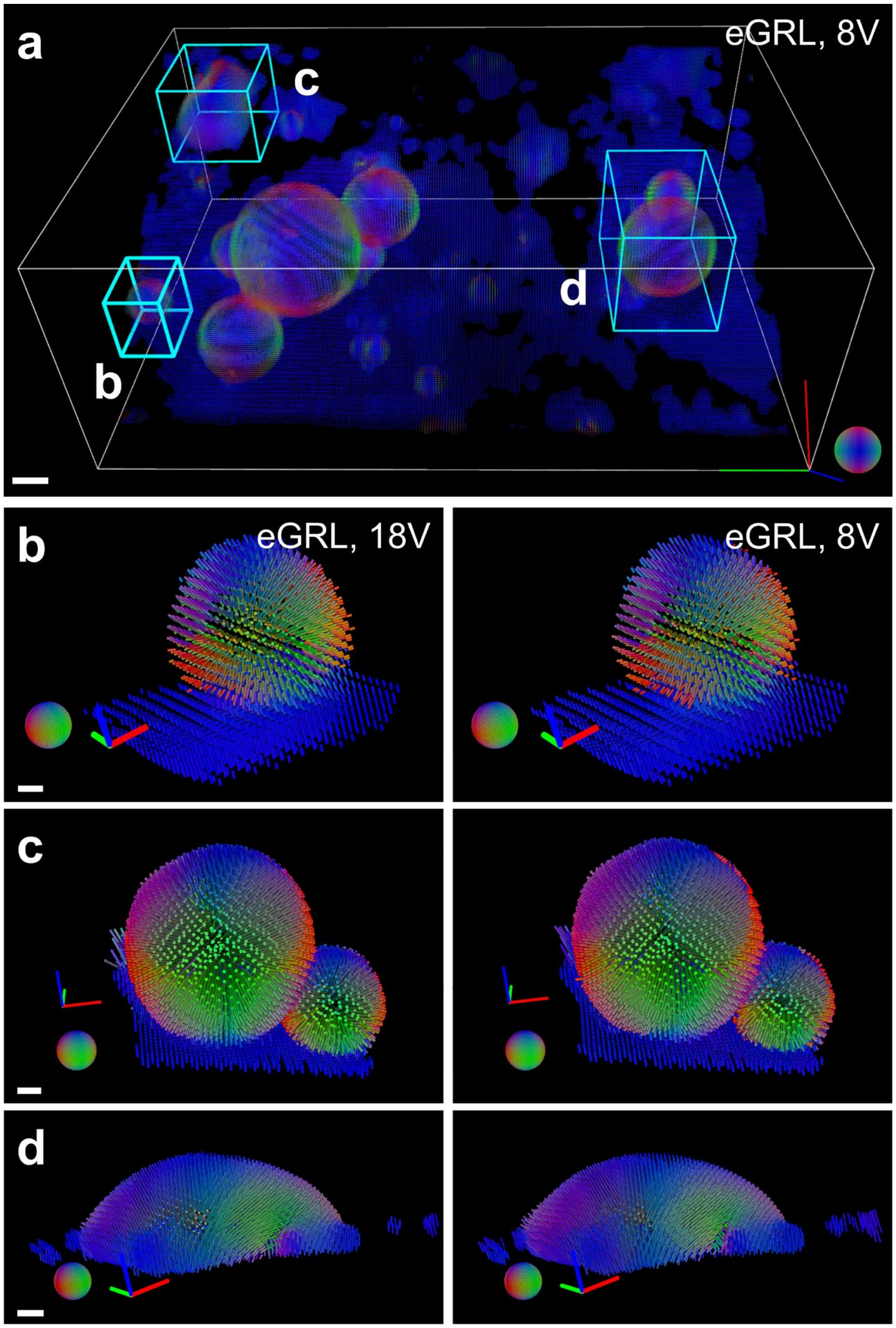

**Supplementary Fig. 12, Performance of eGRL with a reduced number of polarization modulations on GUV sample. a)** eGRL reconstruction of FM1-43 labelled GUVs used only 8 polarization modulations (Scheme 2, 8V. See **Supplementary Table 2**), by contrast to the results obtained using a full complement of 18 modulations (Scheme 1, 18V. See **Supplementary Table 2**) as shown in **Fig. 2a**. **b-d)** Comparisons of eGRL reconstructions using both 18 and 8 polarization modulations across the 3 cyan-labelled cubic regions specified in **a**. Scale bars: **a** 5 µm, **b** 1 µm, **c**, **d** 2 µm.

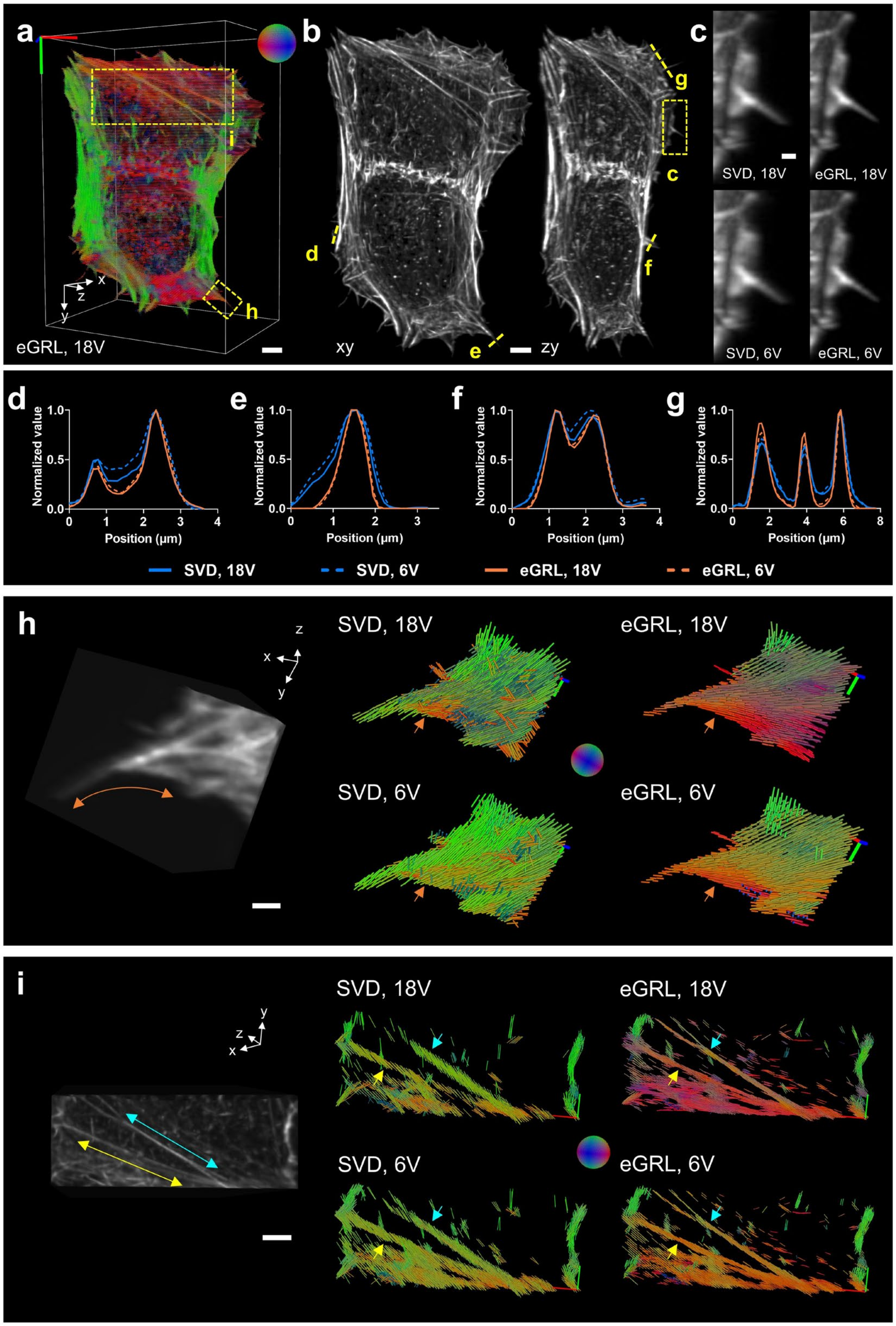

**Supplementary Fig. 13, Performance of eGRL and SVD under fewer polarization modulations. a)** Peak orientations in eGRL reconstruction of a fixed U2OScell labeled with Alexa Fluor 488 phalloidin with 18 polarization modulations. **b)** Lateral (left) and axial (right) maximum-intensity projections of the density map from **a)**. **c)** Higher-magnification views of the rectangular region in **b)** reconstructed by eGRL and SVD with 18 (top) and 6 (bottom) polarization modulations. **d-g)** Comparison of profiles along the four dashed lines in **b)**. **h, i)** Density map (from eGRL result, 18 polarization modulations) and corresponding reconstructions of the two rectangular regions in **a)** with 18 (top) and 6 (bottom) polarization modulations. Note orientations highlighted by arrows should align with the direction of actin fibers indicated by bidirectional arrows with corresponding color. Scale bars: **a, b** 5 µm, **c** 1 µm, **h** 2 µm, **i** 4 µm.

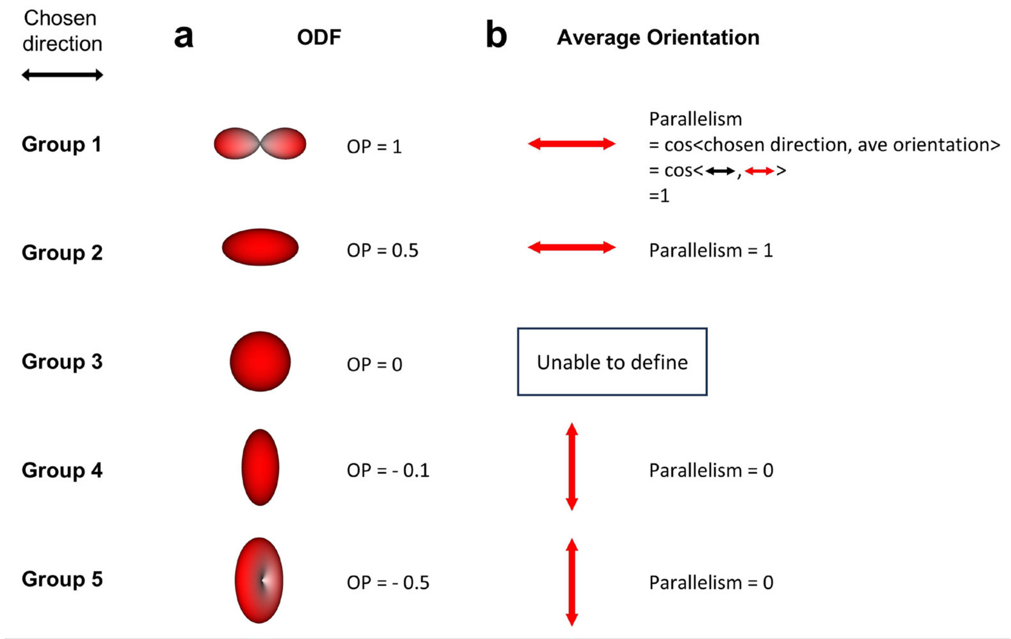

**Supplementary Fig. 14, ODFs and OPs provides a useful assessment of dipoles' preference towards a chosen direction. a)** The ODF shapes of five different groups of dipoles and the corresponding order parameter (OP) values with respect to the direction shown at upper left (labeled as 'chosen direction'). **b)** The average orientations of dipoles in **a)**, which are often used as the output from a traditional polarization microscope. We also compute the corresponding parallelism index between the average orientation vector and the chosen direction. Note the average orientation or parallelism alone cannot distinguish groups 1 and 2 (or groups 4 and 5), demonstrating the usefulness of ODFs and OPs in these applications.

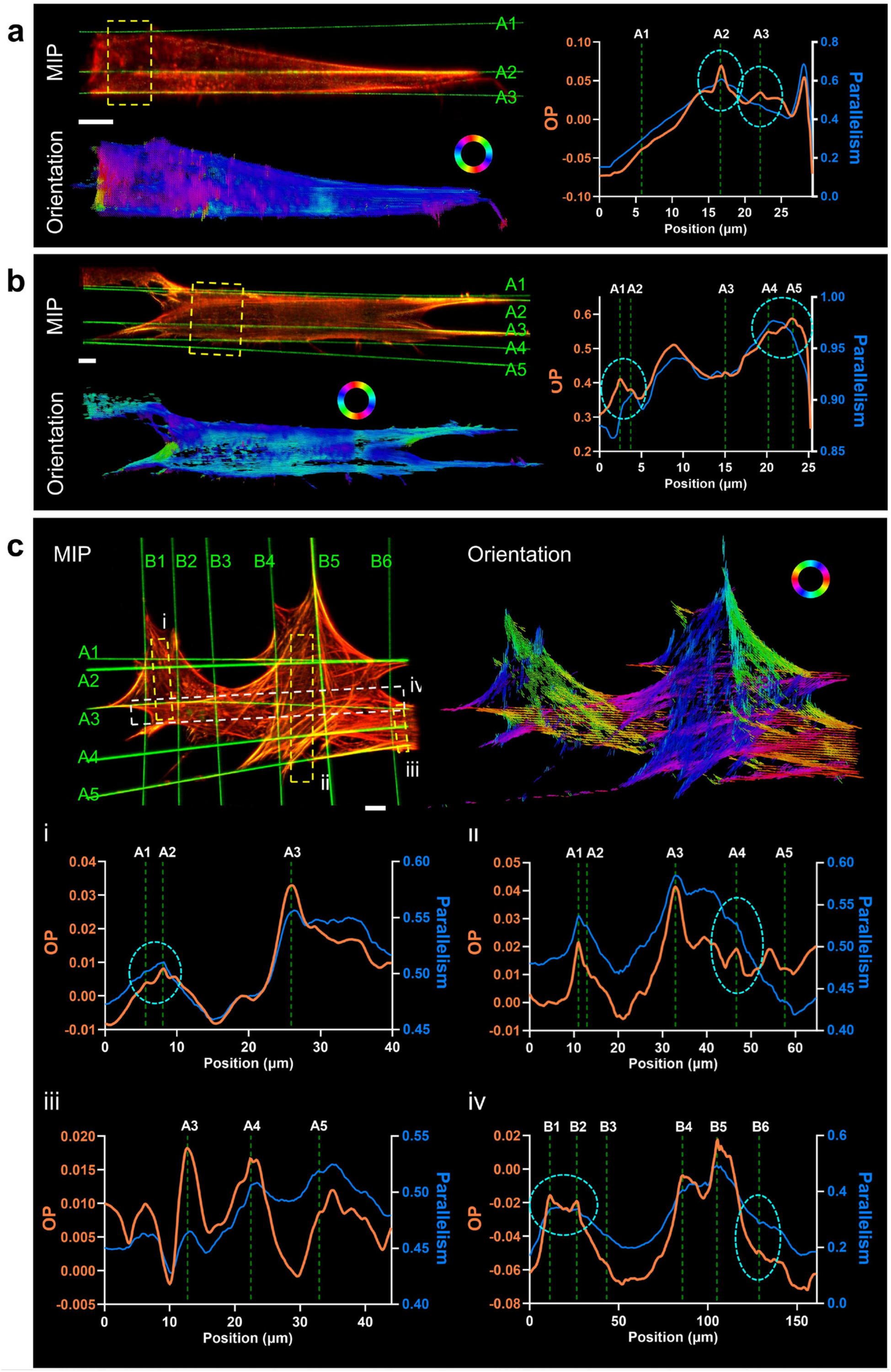

**Supplementary Fig. 15, Spatio-angular reconstructions show correlations between the direction of actin molecules and the direction of nanowires upon which the cells are grown. a)** A spindle shaped fixed NIH3T3 cell labeled with Alexa Fluor 568 Phalloidin (red) grown on multiple nanowires (green), showing lateral maximum intensity projection of density map (top) and average orientation map (bottom, shown by a colormap encoded in 2D space). Nanowires are labeled with green characters A1-A3. Right shows the comparison of orientational analysis in the dashed rectangle region, including OP analysis (orange curve) derived from ODFs and parallelism indicator (blue curve) based on previously available average orientations. Here the average orientations are calculated from the ODFs, and parallelism describes the dot product between vectors of dipole orientation and nanowire's direction, another measure of the degree of alignment of dipoles to nanowires. The position axis in this chart corresponds to the long side of the dashed rectangle at left. **b)** Twin cells grown on multiple nanowires in roughly parallel directions. Similar representations and analysis are demonstrated as in **a**. **c)** Extended analysis of the crosshatched actin fibers grown on crisscrossed nanowires in **Fig. 3g**. Note that our spatio-angular imaging and ODF reconstruction demonstrate explicit patterns of dipole preference, which are not discerned in the intensity images or the average orientation detection by traditional PFMs (see especially those comparisons outlined by cyan dashed circles). Scale bars: **a - c** 10 μm.

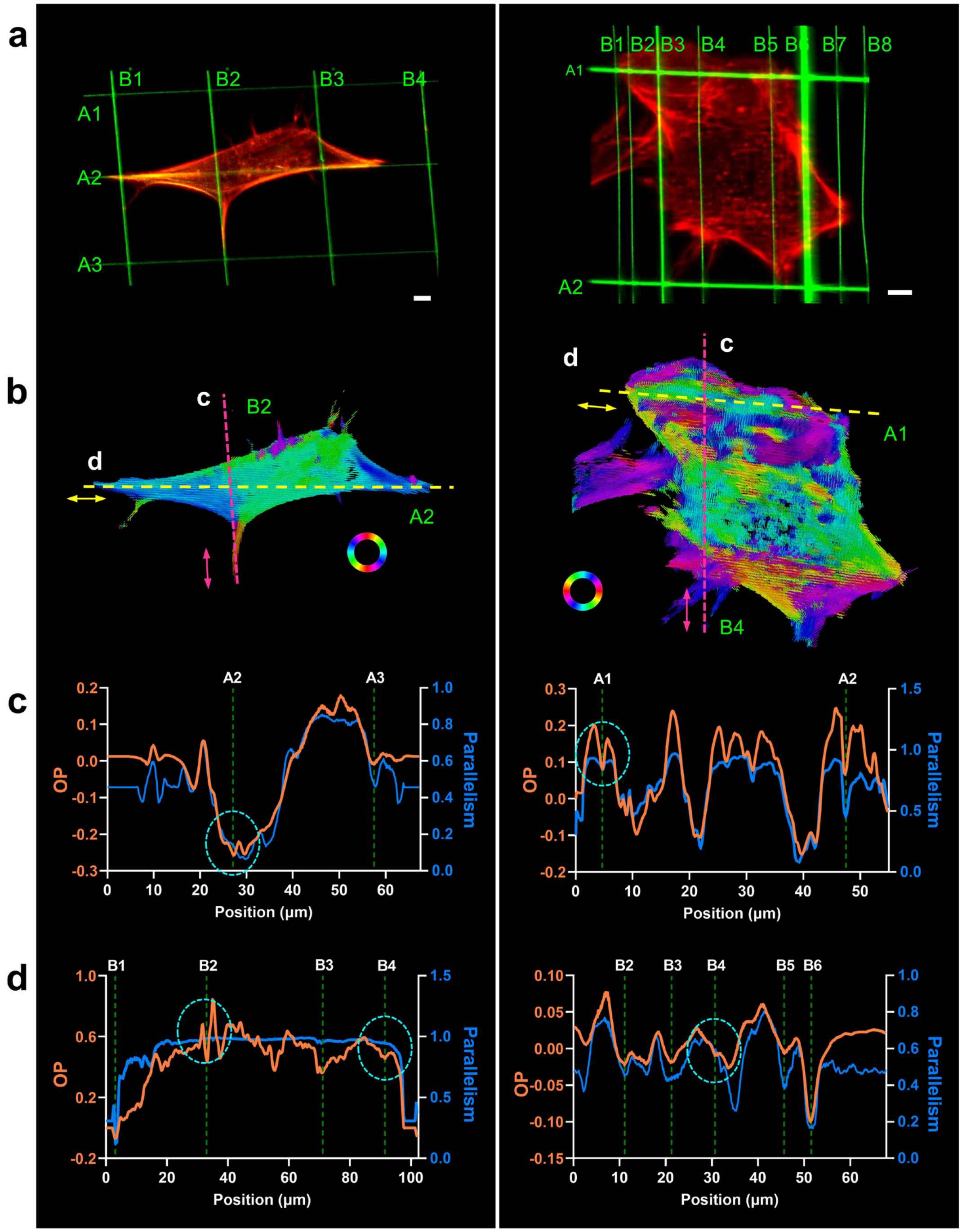

**Supplementary Fig. 16, Examining the change in OP and parallelism in the vicinity of multiple wires.** **a)** Two additional fixed NIH3T3 cells labeled with Alexa Fluor 568 Phalloidin (red) on crosshatched fibers (green). **b)** The peak orientation map of the actin fibers in **a** reconstructed by eGRL, shown by a colormap encoded in 2D space. **c)** OP profile along the vertical nanowires indicated by the pink dashed lines in **b** (average within 1.5 µm of the line), with the OP direction defined by the bidirectional pink arrow. The OP profiles pass through horizontal nanowires and the corresponding wire positions are marked on the profiles. Note the nanowires are mostly collocated with the local minima on the OP curves. **d)** OP profiles along horizontal nanowires indicated by the yellow dashed lines in **b** (average within 1.5 µm of the line), with the OP directions as the bidirectional yellow arrows. The OP profiles pass through vertical nanowires. Same as in **c**, the nanowires are mostly collocated with the local minima on the OP curves. Note these analyses from OP suggest nanowires guide dipoles to align in the tendency of nanowires' directions, which is not always obvious in terms of the parallelism values (the blue curves). See also **Fig. 3**. Scale bars: **a**, **b** 5 µm.

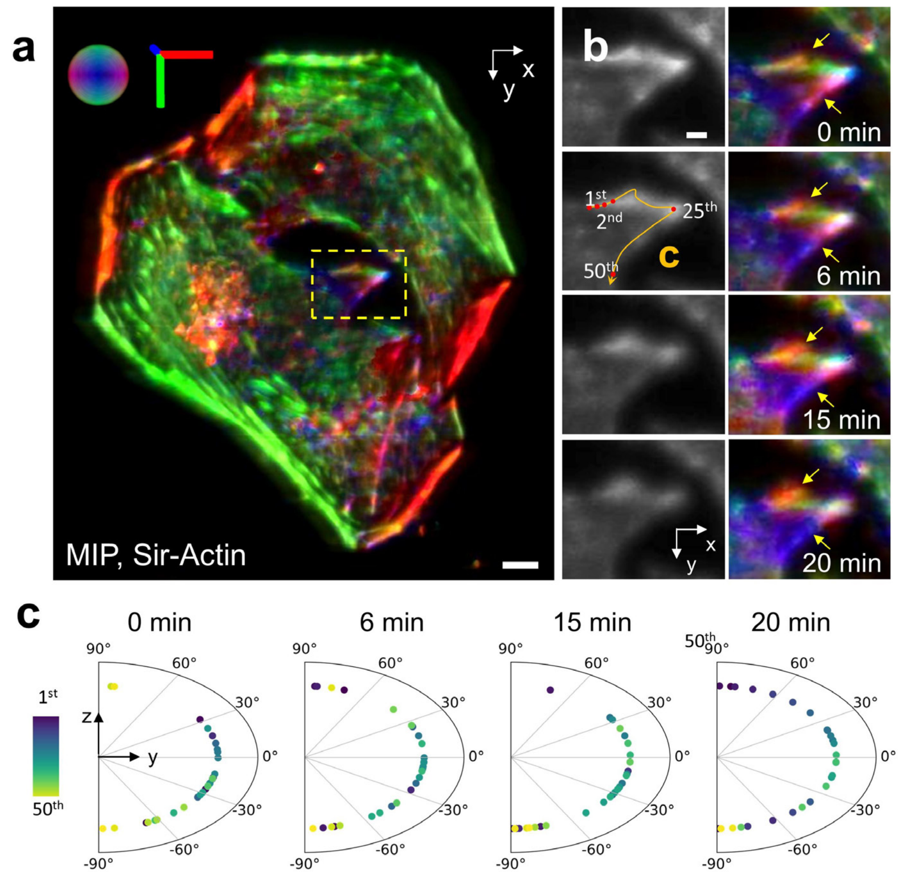

**Supplementary Fig. 17, Orientational distribution and dynamics on live HeLa cell protrusion. a)** The principal orientation map of a live HeLa cell labelled with Sir-Actin and excited at 640 nm. The bottom left panel shows a magnified view of the actin protrusion corresponding to the yellow dashed rectangular region. Note the probes don't align parallel to the coverglass, instead, they are organized in a pattern of continuous spatial rotation in the plane perpendicular to the actin filaments. **b)** A dynamic sequence of the same region outlined by the yellow dashed rectangle in **a**. The arrows point out small changes that are hard to observe in the density map but are noticeable in the orientation distribution. **c)** How principal orientation distributes along the actin filament of protrusion. 50 characteristic points were selected uniformly along the trail in **b**. The polar plots are shown for the inclination angle of the principal orientation in the zy plane, with pseudo color indicating the points' order ($1^{st}$ ~ $50^{th}$) along the trail. Scale bars: **a** 5 µm, **b** 2 µm.

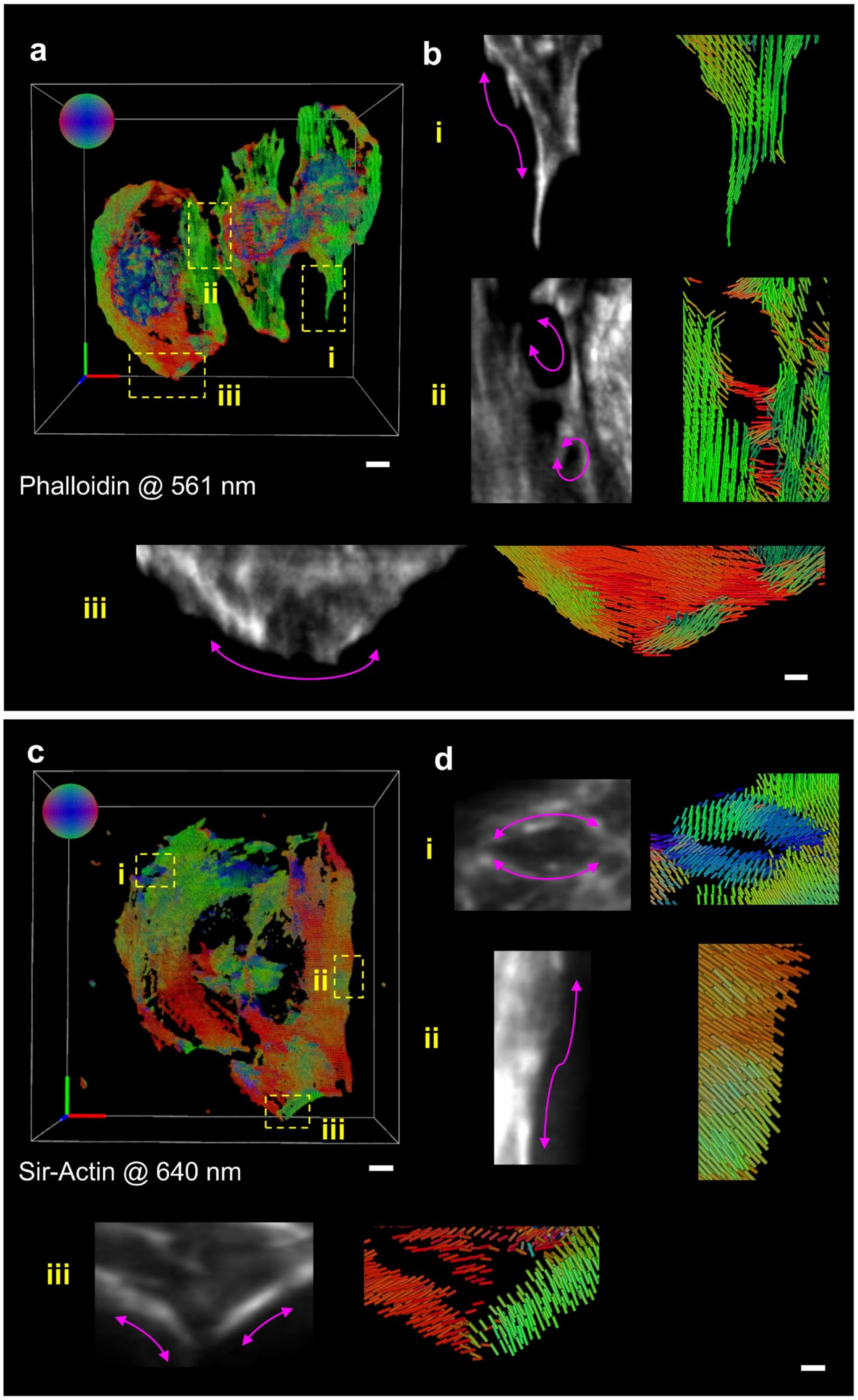

**Supplementary Fig. 18, Comparison of the overall orientation distribution pattern of actin filaments in fixed and live HeLa cells labelled with different dyes. a)** A fixed HeLa cell labeled with Alexa Fluor 568 Phalloidin and imaged under 561 nm excitation. **b)** Higher-magnification views of the orientation map corresponding to the dashed rectangle regions in **a**, highlighting that the orientations align parallel to the direction of actin filaments (profiles shown by magenta arrows). **c)** A live HeLa cell labeled with Sir-Actin and imaged under 640 nm excitation. **d)** Higher-magnification views of the orientation map corresponding to the dashed rectangle regions in **c**, illustrating predominantly perpendicular orientations of Sir-Actin label relative to actin filaments (profiles shown by magenta arrows), in stark contrast to the images in **a**. Additionally, representative ODF maps corresponding to the cyan dashed regions are shown for an in-depth comparison. Scale bars: **a** 5 µm, **b** 1 µm, **c** 5 µm, **d** 1 µm.

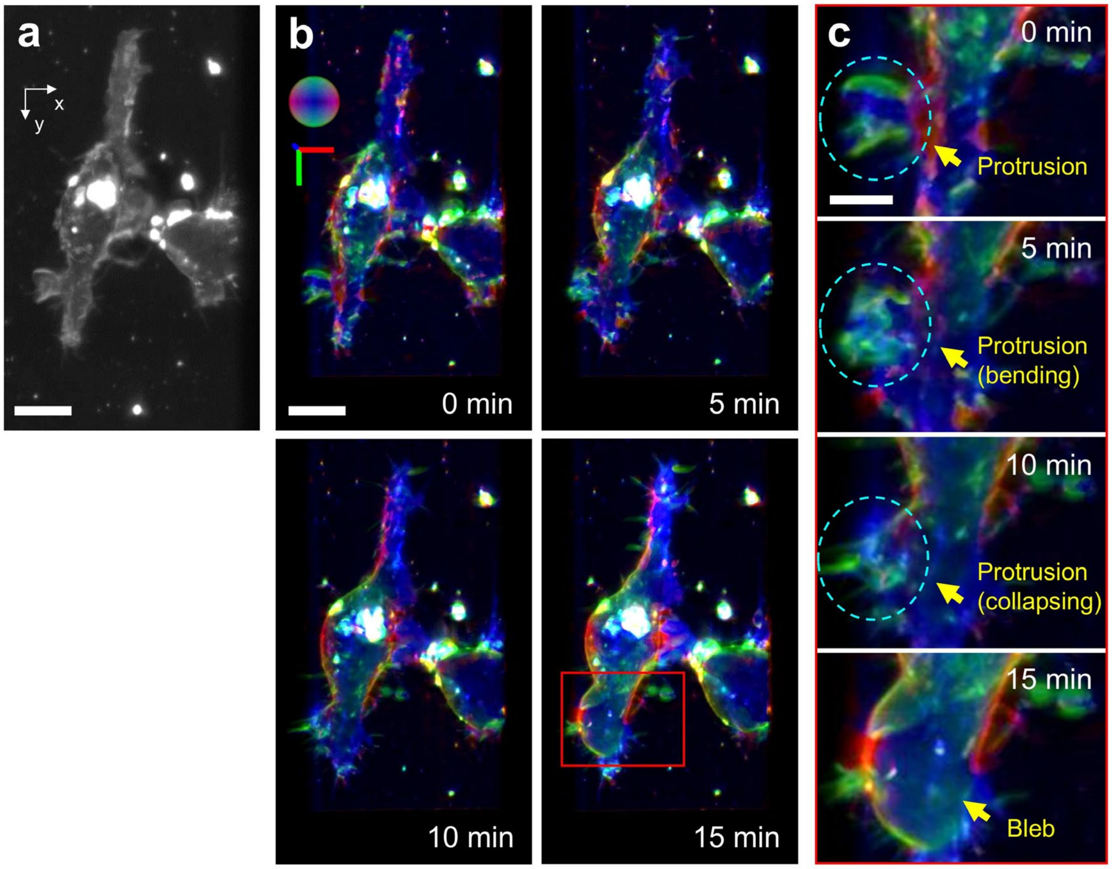

**Supplementary Fig. 19, Membrane transition in terms of morphology and orientation in live macrophage cell labeled with FM1-43. a)** Maximum intensity projection of 3D density map of FM1-43 labelled live macrophage cells. **b)** Time-lapse maximum-intensity-projection images pseudo colored by orientation distribution. **c**) Magnifications of regions outlined with a red rectangle in **b**, yellow arrows distinctly highlight the collapse of membrane protrusions and transition to a bleb, along with orientation echoing the structure. Scale bars: **a**, **b** 10 µm, **c** 5 µm.

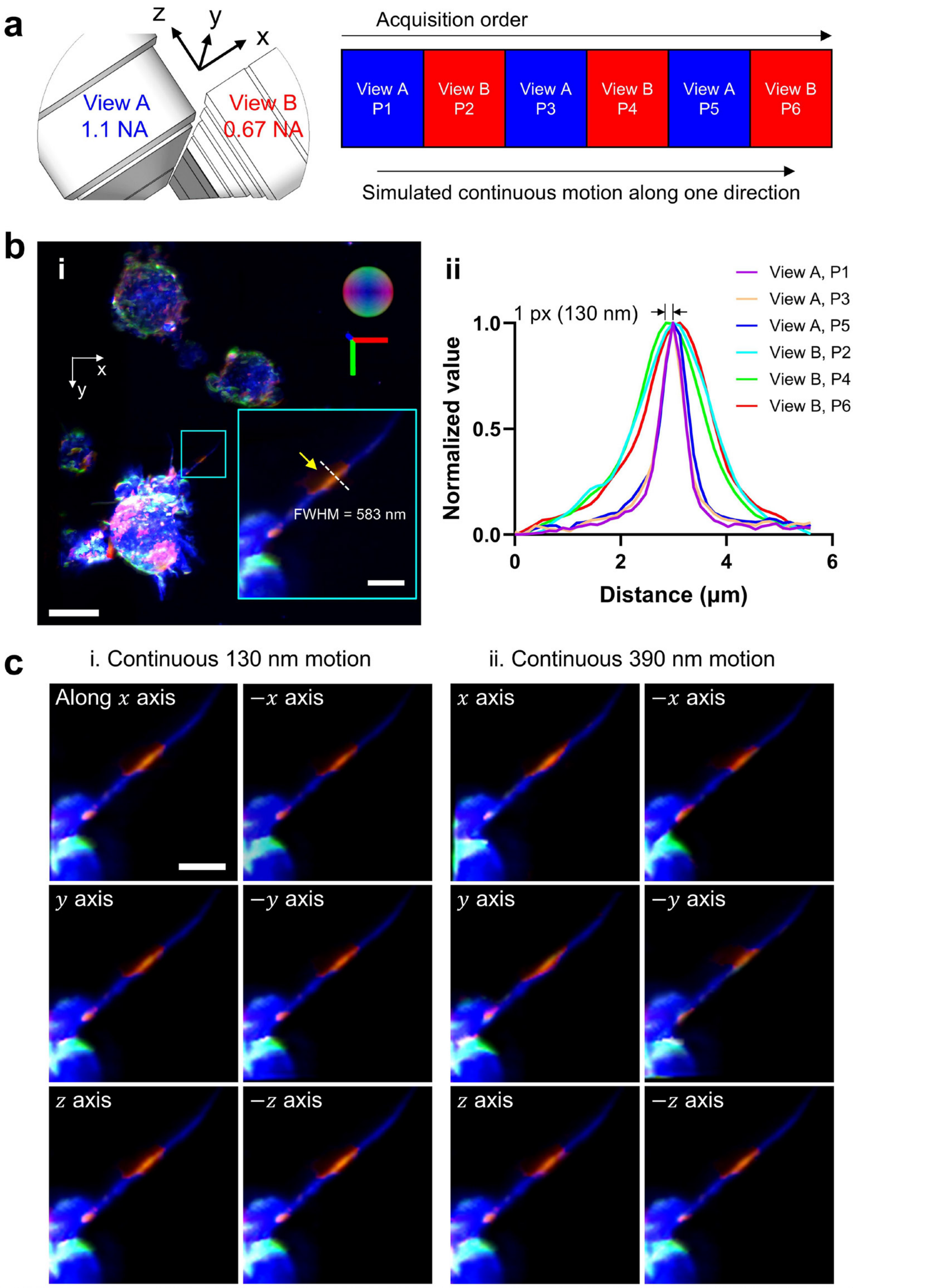

**Supplementary Fig. 20 , Excluding motion artifacts from analysis of macrophage membrane protrusions. a)** Pol-diSPIM acquisition configuration, in which we acquired image volumes in both views (but with different polarizations) before we acquired the next pair of volumes from both views, i.e., two views are acquired in an alternating duty cycle. **b)** i, Maximum intensity projection of multiple FM1-43 labelled live macrophage cells colored by principal orientations at each voxel. Higher magnification view shows the orientation distribution along the membrane protrusion, highlighting a unique orientation patch with the yellow arrow. ii, along each yellow line in **a** i, we draw the profiles extracted from all polarization/view images to show negligible motion during sequential acquisitions under different polarizations/views modulations. Each profile has been normalized to 0 ~ 1. Note considering the acquisition order of polarizations and views, we attribute the 1-pixel (130nm) distance between peaks to polarization modulation rather than cellular motion. **c)** Spatio-angular reconstruction from data with synthetic motion between adjacent volumes by artificially adding 1-pixel (130 nm) shift (**c** i) or 3-pixel (390 nm) shift (**c** ii) successively to each raw image volumes. In the case of 1-pixel shift caused by synthetic motion (**c** i), our cross-validation demonstrates that eGRL can still recover similar results despite motion spanning several pixels over entire 6 raw volume acquisition. Reconstructions remain consistent even with 3-pixel shift caused by synthetic motion (**c** ii). Note the unique patch remains regardless of the additional synthetic motion. Scale bars: **b** 10 µm, magnified inset in **b** 2 µm, **c** 2 µm.

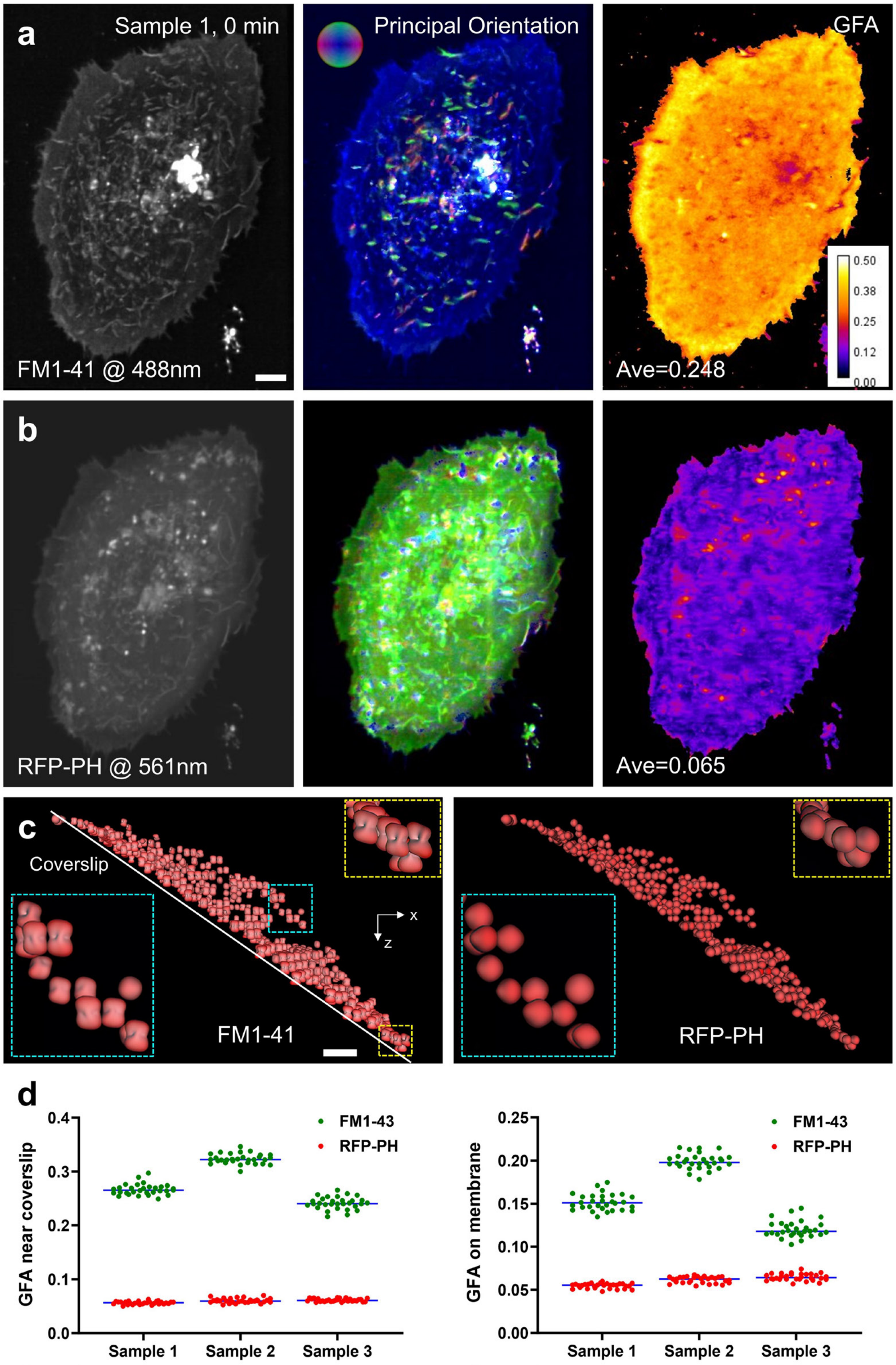

**Supplementary Fig. 21, Polarization imaging of membrane dyes sensitive and insensitive to polarization modulations. a, b)** A live U2OS cell, labeled with FM1-43 (sensitive to polarization modulation) and RFP-PH (insensitive to polarization modulation), was imaged in dual-color channels with 488 nm and 561 nm excitation, left to right showing the density, Principal Orientation, and GFA maps. **c),** Comparison of ODF maps and the magnified shape of individual ODFs for the visualization of probe ensembles. **d)** The GFA statistics of regions on top membrane vs. near coverslip from 3 time-lapse datasets of live U2OS cells with FM1-43 and RFP-PH labeled. For each sample or dye, we calculated the average GFA value for each time point, with GFA values pooled from all the time points represented by points in these graphs. Note polarization-sensitive FM1-43 dye shows anisotropic behavior, with more polarized ODFs and enhanced GFA values over the polarization-insensitive RFP-PH. In addition, FM1-43 exhibits a contrasting GFA level between coverslip and elsewhere while the polarization-insensitive RFP-PH dye shows similar GFA in regions on membrane and coverslip, preliminarily ruling out the spatial motion as the source of lower GFA on membrane elsewhere than coverslip, echoing the observations in **Fig. 5h**. Scale bars: 5 µm.

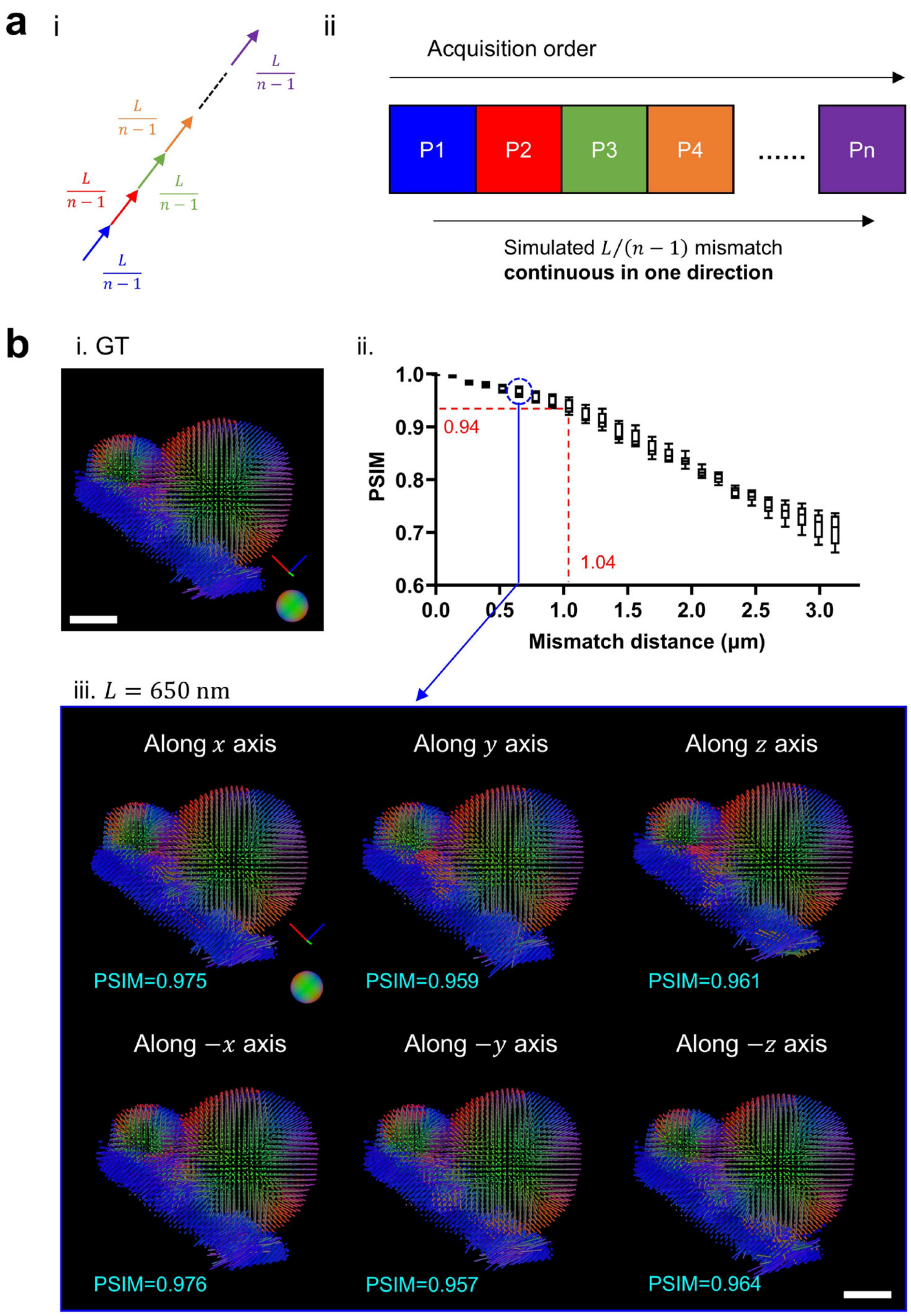

**Supplementary Fig. 22, Evaluating the maximum mismatch distance that is tolerable in eGRL reconstruction on GUV dataset. a)** i, We assume that, between each pair of two temporally adjacent volumes, the object always moves a certain distance $L/(n-1)$ in one certain direction, a total Euclidean distance of $L$ during the entire acquisition. ii, in the sequence of volumetric polarization measurements ($P_1$~$P_n$), we implemented the same shift for each volume. **b)** i. In the GUV dataset, we applied this synthetic motion along one axis. ii, box plot of PSIM values under varying levels of induced motion $L$. The experiment was repeated times along different axes including x, -x, y, -y, z, and -z. Specifically, PSIM suffers a ~5% drop when introducing ~1 μm of continuous motion in one direction.

iii, Orientation map of eGRL if 650 nm synthetic motion is added along indicated axes. Scale bars: **b** 5 µm.

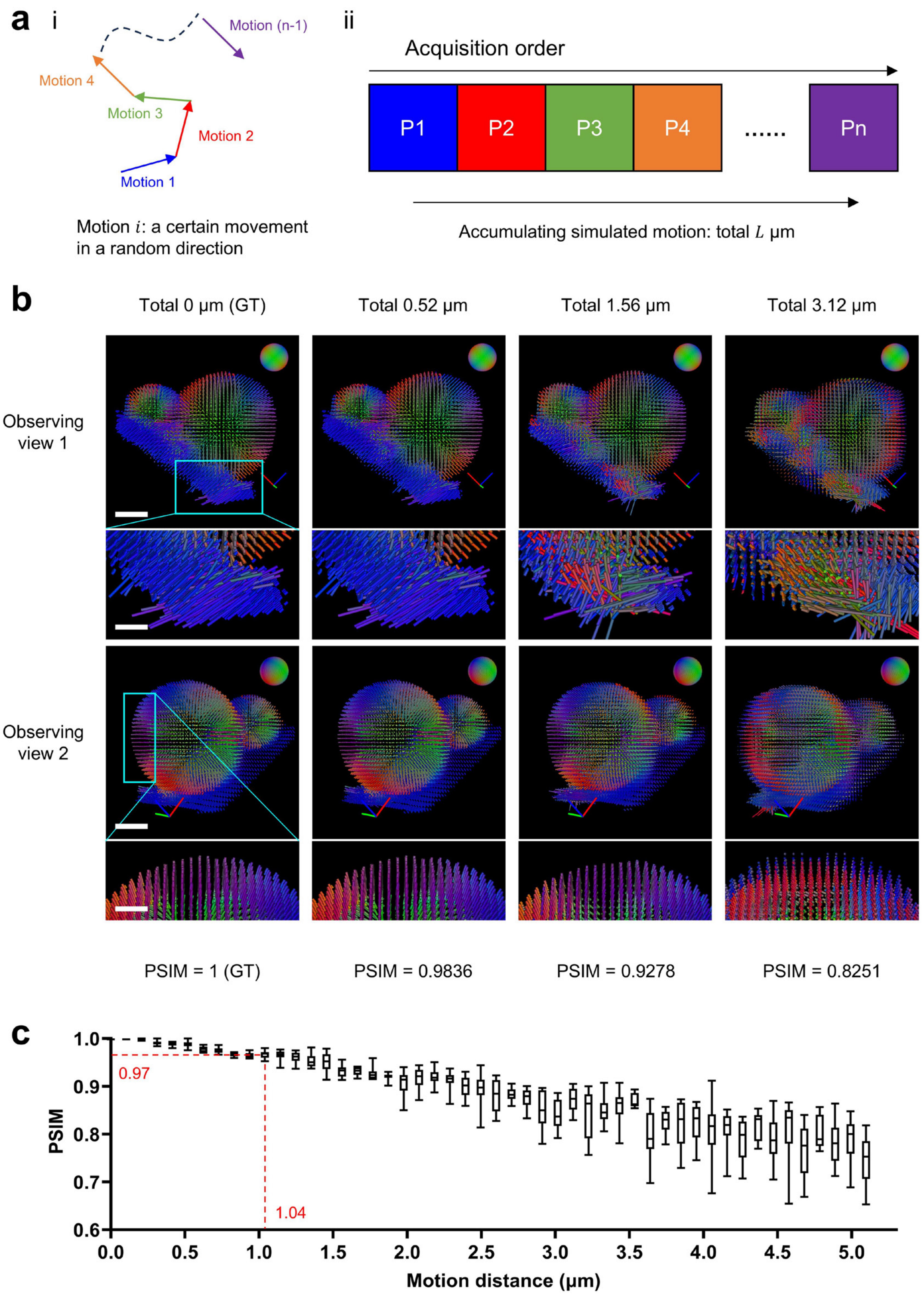

**Supplementary Fig. 23, Assessing motion effects on eGRL reconstruction by artificially introducing random movements in GUV dataset. a)** i, Another case which assumes, between volumes captured under two temporally adjacent polarization modulations, the object moves a certain distance in a random direction. ii, in the sequence of volumetric polarization measurements ($P_1$~$P_n$), we shifted each volume randomly with the same distance $L/(n-1)$ compared with the previous volume. In other words, the volume captured under the $i^{th}$ modulation experiences $L$ drift compared with the volume at the first modulation. **b)** We applied this synthetic motion at indicated $L$ to the fixed GUV dataset with 8 polarization modulations and then performed eGRL reconstruction, and show two views of the eGRL reconstructed orientation maps, including higher magnification insets and PSIM to quantify how the reconstructions are distorted by different levels of motion. Note the orientation distributions are well maintained under random motion of ~1.56 µm, showing eGRL reconstruction can tolerate slight motion. **c)** Box plot of PSIM values under varying levels of added motion. The experiment was repeated 10 times for each condition. eGRL results maintain a high fidelity (PSIM = 0.97) even when ~1 µm random movement is induced. Scale bars: **b** 5 µm, magnified inset in **b** 2.5 µm.

### Supplementary Tables

**Supplementary Table 1,** Beam polarization and tilt configurations in pol-diSPIM.

| Polarization Settings | $P_0$ | $P_1$ | $P_2$ | $P_3$ | $P_4$ | $P_5$ |
|---|---|---|---|---|---|---|
| Angle (deg, to Y axis) | 0 | 45 | 60 | 90 | 120 | 135 |
| Tilt Settings | View A (1.1-NA det, 0.67-NA exc) | | | View B (0.67-NA det, 1.1-NA exc) | | |
| | $T_{-1}$ | $T_0$ | $T_1$ | $T_{-1}$ | $T_0$ | $T_1$ |
| Angle (deg, to Z or X axis) | -5.3 | 0.2 | 9.2 | -6.5 | 0.2 | 4.8 |

See schematic sketch in **Supplementary. Fig. 9**.

**Supplementary Table 2,** Excitation modulation schemes with different polarization and tilt combinations in pol-diSPIM.

| | View A | View B |
|---|---|---|
| Scheme 0:<br>full combinations,<br>42 volumes (42V) | $T_0$ with $P_0$, $P_1$, $P_2$, $P_3$, $P_4$, $P_5$, $P_0$<br>$T_1$ with $P_0$, $P_1$, $P_2$, $P_3$, $P_4$, $P_5$, $P_0$<br>$T_{-1}$ with $P_0$, $P_1$, $P_2$, $P_3$, $P_4$, $P_5$, $P_0$ | $T_0$ with $P_0$, $P_1$, $P_2$, $P_3$, $P_4$, $P_5$, $P_0$<br>$T_1$ with $P_0$, $P_1$, $P_2$, $P_3$, $P_4$, $P_5$, $P_0$<br>$T_{-1}$ with $P_0$, $P_1$, $P_2$, $P_3$, $P_4$, $P_5$, $P_0$ |
| Scheme 1:<br>18 volumes (18V) | $T_0$ with $P_0$, $P_2$, $P_4$<br>$T_1$ with $P_0$, $P_2$, $P_4$<br>$T_{-1}$ with $P_0$, $P_2$, $P_4$ | $T_0$ with $P_0$, $P_2$, $P_4$<br>$T_1$ with $P_0$, $P_2$, $P_4$<br>$T_{-1}$ with $P_0$, $P_2$, $P_4$ |
| Scheme 2:<br>8 volumes (8V) | $T_1$ - $P_2$<br>$T_{-1}$ - $P_0$<br>$T_{-1}$ - $P_2$<br>$T_{-1}$ - $P_4$ | $T_0$ - $P_2$<br>$T_1$ - $P_0$<br>$T_{-1}$ - $P_0$<br>$T_{-1}$ - $P_4$ |
| Scheme 3:<br>6 volumes (6V) | $T_1$ - $P_0$<br>$T_1$ - $P_2$<br>$T_1$ - $P_4$ | $T_0$ - $P_2$<br>$T_1$ - $P_4$<br>$T_{-1}$ - $P_4$ |

**Supplementary Table 3,** Data acquisition and processing details for experimental data of fixed cells.

<table>
<tr><td colspan="2" rowspan="2">Samples</td><td colspan="2" rowspan="2">Fixed GUV<br>Membrane</td><td colspan="3" rowspan="2">Fixed U2OS<br>Actin</td><td colspan="2" rowspan="2">Fixed tobacco<br>xylem<br>Actin</td><td colspan="6">NIH3T3<br>Nanowire/Actin</td><td rowspan="2">Fixed HeLa<br>Actin</td></tr>
<tr><td colspan="2">Spindle</td><td>Two-fiber</td><td colspan="3">Cross-hatched</td></tr>
<tr><td colspan="2">Figures/Videos</td><td colspan="2">Fig. 3a, b,<br>Sup. Fig. 11,<br>12, 22, 23,<br>Sup. Video 5-7</td><td>Fig. 3c-e,<br>Sup. Fig. 10,<br>Sup. Video 8</td><td colspan="2">Sup. Fig. 13,<br>Sup. Video 10</td><td colspan="2">Fig. 3f-j,<br>Sup. Video 9, 11</td><td>Fig. 4a-f</td><td>Sup. Fig. 15a</td><td>Sup. Fig. 15b</td><td>Fig. 4g-j,<br>Sup. Fig. 15c</td><td>Sup. Fig. 16<br>left</td><td>Sup. Fig. 16<br>right</td><td>Sup. Fig. 18a,<br>b</td></tr>
<tr><td colspan="2">Fluorescence Label</td><td colspan="2">FM1-43</td><td colspan="3">Alexa Fluor 488 phalloidin</td><td colspan="2">Pontamine fast<br>scarlet</td><td colspan="6">labBDP FL maleimide,<br>Alexa Fluor 568 Phalloidin</td><td>Phalloidin</td></tr>
<tr><td colspan="2">Color number</td><td colspan="2">1</td><td colspan="3">1</td><td colspan="2">1</td><td colspan="6">2</td><td>1</td></tr>
<tr><td rowspan="8">Acquisition</td><td>Polarization num</td><td colspan="2">42</td><td colspan="3">42</td><td colspan="2">42</td><td colspan="6">6</td><td>42</td></tr>
<tr><td>Excitation</td><td colspan="2">488</td><td colspan="3">488</td><td colspan="2">561</td><td colspan="6">488, 561</td><td>561</td></tr>
<tr><td>Step size × Slices</td><td colspan="2">1 µm × 65</td><td>0.8 µm × 80</td><td colspan="2">1 µm × 70</td><td colspan="2">1 µm × 100</td><td>1 µm × 50</td><td>1 µm × 55</td><td>1 µm × 80</td><td>1 µm × 198</td><td>1 µm × 100</td><td>1 µm × 90</td><td>1 µm × 110</td></tr>
<tr><td>Exposure time/slice</td><td colspan="2">15 ms</td><td>30 ms</td><td colspan="2">30 ms</td><td colspan="2">50 ms</td><td>50 ms</td><td>50 ms</td><td>50 ms</td><td>50 ms</td><td>50 ms</td><td>50 ms</td><td>20 ms</td></tr>
<tr><td>Acquisition time / time point</td><td colspan="2">41.0 s</td><td>100.8 s</td><td colspan="2">88.2 s</td><td colspan="2">210 s</td><td>15 s</td><td>16.5 s</td><td>24 s</td><td>59.4 s</td><td>30 s</td><td>27 s</td><td>92.4 s</td></tr>
<tr><td>Time interval</td><td colspan="2">--</td><td>--</td><td colspan="2">--</td><td colspan="2">--</td><td>--</td><td>--</td><td>--</td><td>--</td><td>--</td><td>--</td><td>--</td></tr>
<tr><td>Total time points</td><td colspan="2">1</td><td>1</td><td colspan="2">1</td><td colspan="2">1</td><td>1</td><td>1</td><td>1</td><td>1</td><td>1</td><td>1</td><td>1</td></tr>
<tr><td>Total time</td><td colspan="2">40.95 s</td><td>100.8 s</td><td colspan="2">88.2 s</td><td colspan="2">210 s</td><td>15 s</td><td>16.5 s</td><td>24 s</td><td>59.4 s</td><td>30 s</td><td>27 s</td><td>92.4 s</td></tr>
<tr><td colspan="2">Image size (each volume after cropping and interpolation)</td><td colspan="2">512×760×275</td><td>464×640×338</td><td colspan="2">440×640×296</td><td colspan="2">408×316×270</td><td>170×1226×156</td><td>202×910×207</td><td>298×1430×237</td><td>1348×1232×807</td><td>490×796×372</td><td>502×444×355</td><td>592×566×465</td></tr>
<tr><td colspan="2">Total data size</td><td colspan="2">4.2 G voxels,<br>8.4 GB, 16 bit</td><td>3.9 G voxels,<br>7.8 GB, 16 bit</td><td colspan="2">3.3 G voxels,<br>6.5 GB, 16 bit</td><td colspan="2">1.4 G voxels, 2.7<br>GB, 16 bit</td><td>186 M voxels, 372<br>MB, 16 bit</td><td>218 M voxels, 436<br>MB, 16 bit</td><td>578 M voxels, 1.13<br>GB, 16 bit</td><td>7.5 G voxels,<br>15 GB, 16 bit</td><td>830 M voxels,<br>1.62 GB, 16 bit</td><td>453 M voxels,<br>906 MB, 16 bit</td><td>6 G voxels,<br>12 GB, 16 bit</td></tr>
<tr><td colspan="2">Polarization used</td><td>18</td><td>8</td><td>18</td><td>18</td><td>6</td><td>18</td><td>8</td><td>6</td><td>6</td><td>6</td><td>6</td><td>6</td><td>6</td><td>18</td></tr>
<tr><td colspan="2">Reconstruction time</td><td>8 min</td><td>7 min</td><td>7 min</td><td>6 min</td><td>5 min</td><td>3 min</td><td>3 min</td><td>2 min</td><td>3 min</td><td>6 min</td><td>71 min</td><td>9 min</td><td>6 min</td><td>10 min</td></tr>
</table>

**Supplementary Table 4,** Data acquisition and processing details for experimental data of live cells

| Samples | | Live HeLa Actin | | Live Macrophage Membrane | | Live U2OS Membrane | | | |
|---|---|---|---|---|---|---|---|---|---|
| Figures/Videos | | Fig. 5a-c, Sup. Fig. 17, Sup. Video 12 | Sup. Fig. 18c, d | Fig. 5d, e, Sup. Fig. 20, Sup. Video 14 | Sup. Fig. 19, Sup. Video 13 | Fig. 5f-k, Sup. Video 16 | Sup. Fig. 21a-c, Sup. Video 15 bottom | Sup. Fig. 21d Sample 2, Sup. Video 15 top | Sup. Fig. 21d Sample 3 |
| Fluorescence Label | | Sir-Actin | | FM1-43 | | FM1-43 | FM1-43, RFP-PH | | |
| Color number | | 1 | | 1 | | 1 | 2 | | |
| Acquisition | Polarization num | 8 | | 6 | 8 | 6 | 6 | | |
| | Excitation | 640 | | 488 | | 488 | 488, 561 | | |
| | Step size × Slices | 1 μm × 100 | 1 μm × 90 | 1 μm × 120 | 1 μm × 48 | 1 μm × 65 | 1 μm × 70 | | |
| | Exposure time/slice | 20 ms | 20 ms | 20 ms | 20 ms | 10 ms | 10 ms | | |
| | Acquisition time / time point | 16 s | 14.4 s | 14.4 s | 7.7 s | 3.9 s | 4.2 s | | |
| | Time interval | 60 s | 60 s | 30 s | 30 s | 20 s | 20 s | | |
| | Total time points | 30 | 30 | 30 | 30 | 60 | 60 | | |
| | Total time | 30 min | 30 min | 15 min | 15 min | 20 min | 20 min | | |
| Image size (each volume after cropping and interpolation) | | 543×648×405 | 572×580×380 | 700×700×400 | 340×565×203 | 300×300×215 | 360×480×275 | 360×360×296 | 360×460×279 |
| Total data size | | 31.8 G voxels, 63.6 GB, 16 bit | 28.1 G voxels, 56.2 GB, 16 bit | 32.8 G voxels, 65.6 GB, 16 bit | 8.7 G voxels, 17.4 GB, 16 bit | 6.5 G voxel, 13.0 GB, 16 bit | 31.8 G voxels, 63.7 GB, 16 bit | 25.6 G voxels, 51.2 GB, 16 bit | 30.8 G voxels, 61.6 GB, 16 bit |
| Polarization used | | 8 | 8 | 6 | 8 | 6 | 6 | 6 | 6 |
| Reconstruction time | | 280 min | 245 min | 304 min | 84 min | 91 min | 228 min | 190 min | 228 min |

### Supplementary Notes

#### Supplementary Note 1, Details of GRL scheme

##### 1.1 The general framework of GRL compared with traditional RL

1.1.1 Traditional imaging process and RL framework

Traditionally in fluorescence microscopy, fluorophores are treated as monopoles, and polarization is rarely considered. If the density of fluorophores at location $\mathbf{r}_o$ is $f(\mathbf{r}_o)$, and the microscope has an emission point spread function (PSF) $h(\mathbf{r})$, the image recorded by the camera can be written as:

$$g(\mathbf{r}_d) = \int_{\mathbb{R}^3} \mathrm{d}\mathbf{r}_o \, h(\mathbf{r}_d - \mathbf{r}_o) f(\mathbf{r}_o) \tag{1}$$

where $\mathbf{r}_d$ is the 3D coordinate vector in the image space and $\mathbf{r}_o$ is the same in the object space.

To partially reverse degradation introduced by blurring and Poisson noise, the Richardson-Lucy (RL) algorithm is widely used. The iterative form of this maximum-likelihood expectation-maximization (MLEM) algorithm has the following expression:

$$e_{k+1} = e_k \times \frac{1}{c} \times \left(\frac{i}{e_k * h} * h^{\mathrm{back}}\right) \tag{2}$$

where $e_k$ is the $k$-th estimate of the desired object image, $e_{k+1}$ is the $(k+1)$-th estimate. $i$ is the measured image, $h$ the forward projector, $h^{\mathrm{back}}$ the back projector, and $*$ denotes the convolution. The PSF is typically used for $h$, and $h^{\mathrm{back}}$ is traditionally matched to $h$ as its transpose by flipping the PSF. $c$ is called the vector of object-space sensitivity constants[2]:

$$c = \int_{\mathbb{R}^3} \mathrm{d}\mathbf{r} \, h(\mathbf{r}) \tag{3}$$

However, the monopole assumption is an approximation, and here we more accurately model fluorophores as dipoles. Fluorescent dipoles can be represented as a spatial and angular distribution $f(\mathbf{r}_o, \mathbf{s}_o)$, describing the density of dipoles oriented in the direction $\hat{\mathbf{s}}_o \in \mathbb{S}^2$ and at location $\mathbf{r}_o \in \mathbb{R}^3$.

1.1.2 Spatio-angular imaging process and GRL framework

Using volume imaging polarized fluorescence microscopy (PFM), we can obtain a series of 3D *image* stacks from different detection views with different polarization modulations. We collect several intensity measurements $i_{\hat{\mathbf{p}}}(\mathbf{r}_d)$ with excitation polarization $\hat{\mathbf{p}}$, describing the resulting imaging process as[3]:

$$g_{\hat{\mathbf{p}}}(\mathbf{r}_d) = \int_{\mathbb{R}^3} \mathrm{d}\mathbf{r}_o \int_{\mathbb{S}^2} \mathrm{d}\hat{\mathbf{s}}_o\, h_{\hat{\mathbf{p}}}(\mathbf{r}_d - \mathbf{r}_o, \hat{\mathbf{s}}_o) f(\mathbf{r}_o, \hat{\mathbf{s}}_o) \tag{4}$$

where $\hat{\mathbf{p}}$ is treated as a discrete variable, and $h_{\hat{\mathbf{p}}}$ is the dipole point spread function (PSF) with dipole polarization taken into consideration and determined by the imaging system and the $\hat{\mathbf{p}}$ excitation property of the fluorophore. It's worth noting here that, $g_{\hat{\mathbf{p}}}(\mathbf{r}_d)$, $h_{\hat{\mathbf{p}}}(\mathbf{r}_d - \mathbf{r}_o, \hat{\mathbf{s}}_o)$ and $f(\mathbf{r}_o, \hat{\mathbf{s}}_o)$ descriptions in the spatial and spherical domain are natural nonnegative since the density of dipoles in a certain direction cannot be negative.

Eq. (4) is an extension of Eq. (1) and a more generalized expression of the imaging process that maps the object space to the image space under the dipole assumption, but is no longer a standard convolution. So we define this imaging process, or forward projection, by the operator $\star$ as:

$$f \star h = \int_{\mathbb{R}^3} \mathrm{d}\mathbf{r}_o \int_{\mathbb{S}^2} \mathrm{d}\hat{\mathbf{s}}_o\, h_{\hat{\mathbf{p}}}(\mathbf{r}_d - \mathbf{r}_o, \hat{\mathbf{s}}_o) f(\mathbf{r}_o, \hat{\mathbf{s}}_o) \tag{5}$$

and correspondingly, we define the back projection from the image space to the object space by the operator $\diamond$ as:

$$g \diamond h^{\mathrm{back}} = \sum\nolimits_{\hat{\mathbf{p}}} \int_{\mathbb{R}^3} \mathrm{d}\mathbf{r}_d\, h_{\hat{\mathbf{p}}}^{\mathrm{back}}(\mathbf{r}_o - \mathbf{r}_d, \hat{\mathbf{s}}_o) g_{\hat{\mathbf{p}}}(\mathbf{r}_d) \tag{6}$$

Like the monopole case, to restore the spatial and angular distribution of fluorescent dipoles $f(\mathbf{r}_o, \hat{\mathbf{s}}_o)$, we can use a RL-like algorithm but need to incorporate two additional dimensions, the polarization $\hat{\mathbf{p}}$ and the orientation $\hat{\mathbf{s}}_o$ to adjust the convolution process. We term this as the generalized Richardson-Lucy algorithm (GRL) with the following form:

$$e_{k+1} = e_k \times \frac{1}{c} \times \left( \frac{i}{e_k \star h} \diamond h^{\mathrm{back}} \right) \tag{7}$$

in which, the vector of object-space sensitivity constants is defined as:

$$c(\hat{\mathbf{s}}_o) = \sum_{\hat{\mathbf{p}}} \int_{\mathbb{R}^3} \mathrm{d}\mathbf{r}\, h_{\hat{\mathbf{p}}}(\mathbf{r}, \hat{\mathbf{s}}_o) \quad (8)$$

Unlike traditional RL where we do the summation of the scalar value $h(\mathbf{r})$ for intensity normalization $h/\mathrm{sum}(h)$, GRL requires the vector of object space sensitivity constant Eq. (8) to balance the mapping between multiple dimensions, especially the extra polarization and angular channel (**Figure SN1**). Otherwise, for example in the forward projection, we need to map from all $\hat{\mathbf{p}}$ to each $\hat{\mathbf{s}}_o$, since normalization is carried out for the whole $h$, the difference of magnitude between different $\hat{\mathbf{s}}_o$ will become larger as we iterate, so the iteration won't converge.

##### 1.2 GRL framework implements

Here we split the GRL iteration process in Eq. (7) into several steps as follows:

Step 1, obtain the forward estimation $i'_{\hat{\mathbf{p}}}(\mathbf{r}_d)$ by

$$i'_{\hat{\mathbf{p}}}(\mathbf{r}_d) = \int_{\mathbb{R}^3} \mathrm{d}\mathbf{r}_o \int_{\mathbb{S}^2} \mathrm{d}\hat{\mathbf{s}}_o\, h_{\hat{\mathbf{p}}}(\mathbf{r}_d - \mathbf{r}_o, \hat{\mathbf{s}}_o) e_k(\mathbf{r}_o, \hat{\mathbf{s}}_o) \quad (9)$$

Step 2, obtain the forward correction ratio $i''_{\hat{\mathbf{p}}}(\mathbf{r}_d)$ by

$$i''_{\hat{\mathbf{p}}}(\mathbf{r}_d) = \frac{i_{\hat{\mathbf{p}}}(\mathbf{r}_d)}{i'_{\hat{\mathbf{p}}}(\mathbf{r}_d)} \quad (10)$$

Step 3, obtain the back-projecting update matrix $e'_k(\mathbf{r}_o, \hat{\mathbf{s}}_o)$ by

$$e'_k(\mathbf{r}_o, \hat{\mathbf{s}}_o) = \sum_{\hat{\mathbf{p}}} \int_{\mathbb{R}^3} \mathrm{d}\mathbf{r}_d\, h^{\mathrm{back}}_{\hat{\mathbf{p}}}(\mathbf{r}_o - \mathbf{r}_d, \hat{\mathbf{s}}_o) i''_{\hat{\mathbf{p}}}(\mathbf{r}_d) \quad (11)$$

Step 4, obtain the updated estimation $e''_k(\mathbf{r}_o, \hat{\mathbf{s}}_o)$ by

$$e''_k(\mathbf{r}_o, \hat{\mathbf{s}}_o) = e_k(\mathbf{r}_o, \hat{\mathbf{s}}_o) \times e'_k(\mathbf{r}_o, \hat{\mathbf{s}}_o) \quad (12)$$

Step 5, obtain the latest estimation $e_{k+1}(\mathbf{r}_o, \hat{\mathbf{s}}_o)$ by sensitivity correction:

$$e_{k+1}(\mathbf{r}_o, \hat{\mathbf{s}}_o) = \frac{e''_k(\mathbf{r}_o, \hat{\mathbf{s}}_o)}{c(\hat{\mathbf{s}}_o)} \quad (13)$$

and the initial estimation is usually obtained by:

$$e_0(\mathbf{r}_o, \hat{\mathbf{s}}_o) = \sum_{\hat{\mathbf{p}}} i_{\hat{\mathbf{p}}}(\mathbf{r}_o) \quad (14)$$

In addition, Eq. (9), (11) contain convolution operations in the spatial domain ($\mathbb{R}^3$) that can be calculated in the Fourier domain more rapidly, hence the entire backbone structure of our proposed GRL algorithm is the following:

**Algorithm S1: Generalized Richardson-Lucy algorithm**

Input: $K$, $h_{\hat{\mathbf{p}}}(\mathbf{r},\hat{\mathbf{s}}_o)$, $h_{\hat{\mathbf{p}}}^{\text{back}}(\mathbf{r},\hat{\mathbf{s}}_o)$, $i_{\hat{\mathbf{p}}}(\mathbf{r})$

Output: $e_K(\mathbf{r},\hat{\mathbf{s}}_o)$

---

$e_0(\mathbf{r}_o,\hat{\mathbf{s}}_o) = \sum_{\hat{\mathbf{p}}} i_{\hat{\mathbf{p}}}(\mathbf{r}_o)$

$c(\hat{\mathbf{s}}_o) = \sum_{\hat{\mathbf{p}}} \int_{\mathbb{R}^3} \mathrm{d}\mathbf{r}\, h_{\hat{\mathbf{p}}}(\mathbf{r},\hat{\mathbf{s}}_o)$

For $k = 0, 1, 2, \dots, K-1$:

$E_k(\boldsymbol{\upsilon},\hat{\mathbf{s}}_o) = \mathcal{F}_{\mathbb{R}^3}\{e_k(\mathbf{r},\hat{\mathbf{s}}_o)\}$

$I'_{\hat{\mathbf{p}}}(\boldsymbol{\upsilon}) = \int_{\mathbb{S}^2} \mathrm{d}\hat{\mathbf{s}}_o H_{\hat{\mathbf{p}}}(\boldsymbol{\upsilon},\hat{\mathbf{s}}_o) E_k(\boldsymbol{\upsilon},\hat{\mathbf{s}}_o)$

$i'_{\hat{\mathbf{p}}}(\mathbf{r}) = \mathcal{F}^{-1}_{\mathbb{R}^3}\{I'_{\hat{\mathbf{p}}}(\boldsymbol{\upsilon})\}$

$i''_{\hat{\mathbf{p}}}(\mathbf{r}) = \dfrac{i_{\hat{\mathbf{p}}}(\mathbf{r})}{i'_{\hat{\mathbf{p}}}(\mathbf{r})}$

$I''_{\hat{\mathbf{p}}}(\boldsymbol{\upsilon}) = \mathcal{F}_{\mathbb{R}^3}\{i''_{\hat{\mathbf{p}}}(\mathbf{r})\}$

$E'_k(\boldsymbol{\upsilon},\hat{\mathbf{s}}_o) = \sum_{\hat{\mathbf{p}}} H_{\hat{\mathbf{p}}}(\boldsymbol{\upsilon},\hat{\mathbf{s}}_o) I''_{\hat{\mathbf{p}}}(\boldsymbol{\upsilon})$

$e'_k(\mathbf{r},\hat{\mathbf{s}}_o) = \mathcal{F}^{-1}_{\mathbb{R}^3}\{E'_k(\boldsymbol{\upsilon},\hat{\mathbf{s}}_o)\}$

$e''_k(\mathbf{r},\hat{\mathbf{s}}_o) = e_k(\mathbf{r},\hat{\mathbf{s}}_o) e'_k(\mathbf{r},\hat{\mathbf{s}}_o)$

$e_{k+1}(\mathbf{r},\hat{\mathbf{s}}_o) = \dfrac{e''_k(\mathbf{r}_o,\hat{\mathbf{s}}_o)}{c(\hat{\mathbf{s}}_o)}$

End

---

#### 1.3 Mathematical derivation of GRL

##### 1.3.1 Likelihood function

Let $f$ denote the sample and $i$ denote the image collected from camera. If $i$ has been obtained, we can achieve the estimation of $f$ by maximizing the conditional probability $P(f|i)$. Based on Bayes' theorem, we have:

$$P(f|i) = \frac{P(i|f)P(f)}{P(i)} \tag{15}$$

Since $P(i)$ is independent of $f$, it can be ignored, and we just need to maximize $P(i|f)P(f)$. This is equivalent to maximizing its natural logarithm $K(i|f)$:

$$K(i|f) = \ln[P(i|f)P(f)] = \ln P(i|f) + \gamma \ln P(f) \tag{16}$$

in which, $\ln P(i|f)$ denotes the likelihood function of sample, $\ln P(f)$ and $\gamma$ are the regularization term and a parameter. As $g$ is the ideal forward projection of $f$ and we set the regularization term to 0, maximizing $K(i|f)$ is equivalent to maximizing:

$$L(i|g) = \ln P(i|g) \tag{17}$$

Under the assumption of Poisson noise, the conditional probability density function for an individual pixel at $\mathbf{r}_d$ and polarization $\hat{\mathbf{p}}$ is given by:

$$P\left(i_{\hat{\mathbf{p}}}(\mathbf{r}_d)\middle|g_{\hat{\mathbf{p}}}(\mathbf{r}_d)\right) = \frac{g_{\hat{\mathbf{p}}}(\mathbf{r}_d)^{i_{\hat{\mathbf{p}}}(\mathbf{r}_d)}}{i_{\hat{\mathbf{p}}}(\mathbf{r}_d)!} e^{-g_{\hat{\mathbf{p}}}(\mathbf{r}_d)} \tag{18}$$

Assuming that pixels are independent of each other, we have:

$$P(i|g) = \prod_{\mathbf{r}_d,\hat{\mathbf{p}}} P\left(i_{\hat{\mathbf{p}}}(\mathbf{r}_d)\middle|g_{\hat{\mathbf{p}}}(\mathbf{r}_d)\right) = \prod_{\mathbf{r}_d,\hat{\mathbf{p}}} \frac{g_{\hat{\mathbf{p}}}(\mathbf{r}_d)^{i_{\hat{\mathbf{p}}}(\mathbf{r}_d)}}{i_{\hat{\mathbf{p}}}(\mathbf{r}_d)!} e^{-g_{\hat{\mathbf{p}}}(\mathbf{r}_d)} \tag{19}$$

Then we obtain:

$$L(i|g) = \ln\big(P(i|g)\big) = \sum_{\hat{\mathbf{p}}} \int_{\mathbb{R}^3} \mathrm{d}\mathbf{r}_d \left[-g_{\hat{\mathbf{p}}}(\mathbf{r}_d) + i_{\hat{\mathbf{p}}}(\mathbf{r}_d)\ln\left(g_{\hat{\mathbf{p}}}(\mathbf{r}_d)\right) - \ln\big(i_{\hat{\mathbf{p}}}(\mathbf{r}_d)!\big)\right] \tag{20}$$

1.3.2 Maximize the likelihood function

Estimating the density of dipoles oriented in the direction $\hat{\mathbf{s}}_o$ and at location $\mathbf{r}_o$ in sample $f$ is equivalent to maximizing the likelihood function:

$$f(\mathbf{r}_o, \hat{\mathbf{s}}_o) = \underset{f(\mathbf{r}_o,\hat{\mathbf{s}}_o)>0}{\operatorname{argmax}} K(i|f) = \underset{f(\mathbf{r}_o,\hat{\mathbf{s}}_o)>0}{\operatorname{argmax}} L(i|g) \tag{21}$$

The partial derivative of $f(\mathbf{r}_o, \hat{\mathbf{s}}_o)$ with respect to $L(i|g)$ becomes:

$$\frac{\partial L(i|g)}{\partial f(\mathbf{r}_o, \hat{\mathbf{s}}_o)} = \frac{\partial}{\partial f(\mathbf{r}_o, \hat{\mathbf{s}}_o)} \left\{\sum_{\hat{\mathbf{p}}} \int_{\mathbb{R}^3} \mathrm{d}\mathbf{r}_d \left[-g_{\hat{\mathbf{p}}}(\mathbf{r}_d) + i_{\hat{\mathbf{p}}}(\mathbf{r}_d)\ln\left(g_{\hat{\mathbf{p}}}(\mathbf{r}_d)\right) - \ln\big(i_{\hat{\mathbf{p}}}(\mathbf{r}_d)!\big)\right]\right\} \tag{22}$$

where $g_{\hat{\mathbf{p}}}(\mathbf{r}_d)$ can be represented by $f(\mathbf{r}_o, \hat{\mathbf{s}}_o)$ as:

$$g_{\hat{\mathbf{p}}}(\mathbf{r}_d) = \int_{\mathbb{R}^3} \mathrm{d}\mathbf{r}_o \int_{\mathbb{S}^2} \mathrm{d}\hat{\mathbf{s}}_o \, h_{\hat{\mathbf{p}}}(\mathbf{r}_d - \mathbf{r}_o, \hat{\mathbf{s}}_o) f(\mathbf{r}_o, \hat{\mathbf{s}}_o) \tag{23}$$

Based on Eq. (23), it can also be shown that the derivative of an element of $g_{\hat{\mathbf{p}}}(\mathbf{r}_d)$ with respect to some other element of $f(\mathbf{r}_o, \hat{\mathbf{s}}_o)$ can be writen as

$$\frac{\partial g_{\hat{\mathbf{p}}}(\mathbf{r}_d)}{\partial f(\mathbf{r}_o, \hat{\mathbf{s}}_o)} = h_{\hat{\mathbf{p}}}(\mathbf{r}_d - \mathbf{r}_o, \hat{\mathbf{s}}_o) \tag{24}$$

Hence, the entire partial derivative has the following simplified process:

$$\begin{aligned}
&\frac{\partial L(i|g)}{\partial f(\mathbf{r}_o, \hat{\mathbf{s}}_o)} \\
&= \frac{\partial}{\partial f(\mathbf{r}_o, \hat{\mathbf{s}}_o)} \left\{ \sum\nolimits_{\hat{\mathbf{p}}} \int_{\mathbb{R}^3} \mathrm{d}\mathbf{r}_d \left[ -g_{\hat{\mathbf{p}}}(\mathbf{r}_d) + i_{\hat{\mathbf{p}}}(\mathbf{r}_d) \ln\left( g_{\hat{\mathbf{p}}}(\mathbf{r}_d) \right) - \ln\left( i_{\hat{\mathbf{p}}}(\mathbf{r}_d)! \right) \right] \right\} \\
&= \sum\nolimits_{\hat{\mathbf{p}}} \int_{\mathbb{R}^3} \mathrm{d}\mathbf{r}_d \left\{ \frac{i_{\hat{\mathbf{p}}}(\mathbf{r}_d)}{g_{\hat{\mathbf{p}}}(\mathbf{r}_d)} \frac{\partial}{\partial f(\mathbf{r}_o, \hat{\mathbf{s}}_o)} g_{\hat{\mathbf{p}}}(\mathbf{r}_d) - \frac{\partial}{\partial f(\mathbf{r}_o, \hat{\mathbf{s}}_o)} g_{\hat{\mathbf{p}}}(\mathbf{r}_d) \right\} \\
&= \sum\nolimits_{\hat{\mathbf{p}}} \int_{\mathbb{R}^3} \mathrm{d}\mathbf{r}_d \left[ \frac{i_{\hat{\mathbf{p}}}(\mathbf{r}_d)}{g_{\hat{\mathbf{p}}}(\mathbf{r}_d)} h_{\hat{\mathbf{p}}}(\mathbf{r}_d - \mathbf{r}_o, \hat{\mathbf{s}}_o) - h_{\hat{\mathbf{p}}}(\mathbf{r}_d - \mathbf{r}_o, \hat{\mathbf{s}}_o) \right] \\
&= \sum\nolimits_{\hat{\mathbf{p}}} \int_{\mathbb{R}^3} \mathrm{d}\mathbf{r}_d \left[ \frac{i_{\hat{\mathbf{p}}}(\mathbf{r}_d)}{g_{\hat{\mathbf{p}}}(\mathbf{r}_d)} h_{\hat{\mathbf{p}}}(\mathbf{r}_o - \mathbf{r}_d, \hat{\mathbf{s}}_o) \right] - \sum\nolimits_{\hat{\mathbf{p}}} \int_{\mathbb{R}^3} \mathrm{d}\mathbf{r}_d \, h_{\hat{\mathbf{p}}}(\mathbf{r}_o - \mathbf{r}_d, \hat{\mathbf{s}}_o)
\end{aligned} \tag{25}$$

An extremum of this function occurs at a point where the derivative with respect to all components vanishes. To see whether the extremum is a minimum or a maximum, we take another derivative:

$$\begin{aligned}
&\frac{\partial^2 L(i|g)}{\partial f(\mathbf{r}_o, \hat{\mathbf{s}}_o) \partial f(\mathbf{r}'_o, \hat{\mathbf{s}}'_o)} \\
&= \frac{\partial}{\partial f(\mathbf{r}'_o, \hat{\mathbf{s}}'_o)} \left\{ \sum\nolimits_{\hat{\mathbf{p}}} \int_{\mathbb{R}^3} \mathrm{d}\mathbf{r}_d \left[ \frac{i_{\hat{\mathbf{p}}}(\mathbf{r}_d)}{g_{\hat{\mathbf{p}}}(\mathbf{r}_d)} h_{\hat{\mathbf{p}}}(\mathbf{r}_o - \mathbf{r}_d, \hat{\mathbf{s}}_o) \right] \right\} \\
&= \frac{\partial}{\partial f(\mathbf{r}'_o, \hat{\mathbf{s}}'_o)} \left\{ \sum\nolimits_{\hat{\mathbf{p}}} \int_{\mathbb{R}^3} \mathrm{d}\mathbf{r}_d \left[ \frac{i_{\hat{\mathbf{p}}}(\mathbf{r}_d)}{g_{\hat{\mathbf{p}}}(\mathbf{r}_d)} h_{\hat{\mathbf{p}}}(\mathbf{r}_o - \mathbf{r}_d, \hat{\mathbf{s}}_o) \right] \right\} \\
&= \sum\nolimits_{\hat{\mathbf{p}}} \int_{\mathbb{R}^3} \mathrm{d}\mathbf{r}_d \left[ -\frac{i_{\hat{\mathbf{p}}}(\mathbf{r}_d)}{g^2_{\hat{\mathbf{p}}}(\mathbf{r}_d)} \frac{\partial g_{\hat{\mathbf{p}}}(\mathbf{r}_d)}{\partial f(\mathbf{r}'_o, \hat{\mathbf{s}}'_o)} h_{\hat{\mathbf{p}}}(\mathbf{r}_d - \mathbf{r}_o, \hat{\mathbf{s}}_o) \right] \\
&= \sum\nolimits_{\hat{\mathbf{p}}} \int_{\mathbb{R}^3} \mathrm{d}\mathbf{r}_d \left[ -\frac{i_{\hat{\mathbf{p}}}(\mathbf{r}_d)}{g^2_{\hat{\mathbf{p}}}(\mathbf{r}_d)} h_{\hat{\mathbf{p}}}(\mathbf{r}_d - \mathbf{r}'_o, \hat{\mathbf{s}}'_o) h_{\hat{\mathbf{p}}}(\mathbf{r}_d - \mathbf{r}_o, \hat{\mathbf{s}}_o) \right]
\end{aligned} \tag{26}$$

All components of $i_{\hat{\mathbf{p}}}(\mathbf{r}_d)$, $g_{\hat{\mathbf{p}}}(\mathbf{r}_d)$ and $h_{\hat{\mathbf{p}}}(\mathbf{r}_o - \mathbf{r}_d, \hat{\mathbf{s}}_o)$ are nonnegative. Thus this second derivative is negative everywhere (i.e. the log-likelihood is concave), and any extremum must be a maximum.

Maximizing the likelihood is thus equivalent to letting the partial derivative equal to 0:

$$\frac{\partial L(i|g)}{\partial f(\mathbf{r}_o, \hat{\mathbf{s}}_o)} = 0 \tag{27}$$

so we have:

$$\sum\nolimits_{\hat{\mathbf{p}}}\int_{\mathbb{R}^3}\mathrm{d}\mathbf{r}_d\, h_{\hat{\mathbf{p}}}(\mathbf{r}_o-\mathbf{r}_d,\hat{\mathbf{s}}_o)=\sum\nolimits_{\hat{\mathbf{p}}}\int_{\mathbb{R}^3}\mathrm{d}\mathbf{r}_d\left[\frac{i_{\hat{\mathbf{p}}}(\mathbf{r}_d)}{g_{\hat{\mathbf{p}}}(\mathbf{r}_d)}h_{\hat{\mathbf{p}}}(\mathbf{r}_o-\mathbf{r}_d,\hat{\mathbf{s}}_o)\right] \tag{28}$$

and then:

$$f(\mathbf{r}_o,\hat{\mathbf{s}}_o)\sum\nolimits_{\hat{\mathbf{p}}}\int_{\mathbb{R}^3}\mathrm{d}\mathbf{r}_d\, h_{\hat{\mathbf{p}}}(\mathbf{r}_o-\mathbf{r}_d,\hat{\mathbf{s}}_o)=f(\mathbf{r}_o,\hat{\mathbf{s}}_o)\sum\nolimits_{\hat{\mathbf{p}}}\int_{\mathbb{R}^3}\mathrm{d}\mathbf{r}_d\left[\frac{i_{\hat{\mathbf{p}}}(\mathbf{r}_d)}{g_{\hat{\mathbf{p}}}(\mathbf{r}_d)}h_{\hat{\mathbf{p}}}(\mathbf{r}_o-\mathbf{r}_d,\hat{\mathbf{s}}_o)\right] \tag{29}$$

The GRL algorithm can then be derived as the fixed-point iteration of the above formula:

$$\begin{aligned}
&e_{k+1}(\mathbf{r}_o,\hat{\mathbf{s}}_o)\\
&=e_k(\mathbf{r}_o,\hat{\mathbf{s}}_o)\times\frac{1}{\sum_{\hat{\mathbf{p}}}\int_{\mathbb{R}^3}\mathrm{d}\mathbf{r}_d\, h_{\hat{\mathbf{p}}}(\mathbf{r}_o-\mathbf{r}_d,\hat{\mathbf{s}}_o)}\\
&\times\sum\nolimits_{\hat{\mathbf{p}}}\int_{\mathbb{R}^3}\mathrm{d}\mathbf{r}_d\left[\frac{i_{\hat{\mathbf{p}}}(\mathbf{r}_d)}{g_{\hat{\mathbf{p}}}(\mathbf{r}_d)}h_{\hat{\mathbf{p}}}(\mathbf{r}_o-\mathbf{r}_d,\hat{\mathbf{s}}_o)\right]\\
&=e_k(\mathbf{r}_o,\hat{\mathbf{s}}_o)\times\frac{1}{\sum_{\hat{\mathbf{p}}}\int_{\mathbb{R}^3}\mathrm{d}\mathbf{r}_d\, h_{\hat{\mathbf{p}}}(\mathbf{r}_o-\mathbf{r}_d,\hat{\mathbf{s}}_o)}\\
&\times\sum\nolimits_{\hat{\mathbf{p}}}\int_{\mathbb{R}^3}\mathrm{d}\mathbf{r}_d\left[\frac{i_{\hat{\mathbf{p}}}(\mathbf{r}_d)}{\int_{\mathbb{R}^3}\mathrm{d}\mathbf{r}_o\int_{\mathbb{S}^2}\mathrm{d}\hat{\mathbf{s}}_o\, h_{\hat{\mathbf{p}}}(\mathbf{r}_d-\mathbf{r}_o,\hat{\mathbf{s}}_o)e_k(\mathbf{r}_o,\hat{\mathbf{s}}_o)}\times h_{\hat{\mathbf{p}}}(\mathbf{r}_o-\mathbf{r}_d,\hat{\mathbf{s}}_o)\right]
\end{aligned} \tag{30}$$

For the PSF-related operations defined in Eq. (5)(6)(8), we can achieve the following simplified GRL expression:

$$e_{k+1}=e_k\times\frac{1}{c}\times\left(\frac{i}{e_k\star h}\diamond h^{\mathrm{back}}\right) \tag{31}$$

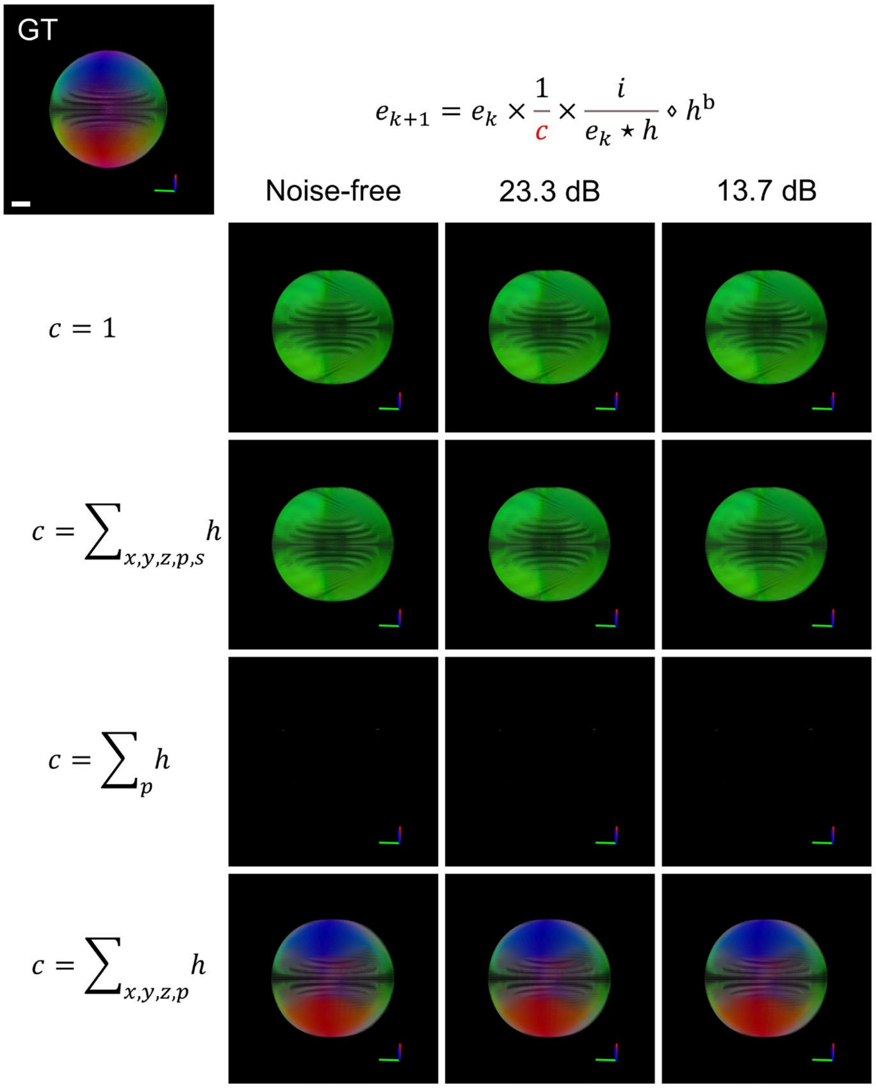

**Figure SN1, Impact of object space sensitive constant on GRL reconstruction.** GRL reconstructions with assorted object space sensitive constants *c* applied to raw data exhibiting varying noise levels, including 1) non-normalized, 2) globally normalized across the polarization *p*, orientation *s* and spatial *xyz* channels, 3) within the polarization channel *p*, and 4) optimally, within just the spatial *xyz* and polarization *p* channels. See also **Methods**. Scale bars: 2 µm.

### Supplementary Note 2, Optimization of GRL scheme

#### 2.1 Angular dimensionality reduction

2.1.1 Imaging process

When doing the calculation in the spherical domain, the brute-force approach would be to discretize $\hat{\mathbf{s}}_o$ into thousands of orientations to approximately cover the whole angular space $\mathbb{S}^2$, but such an approach would imply a massive computational cost. By contrast, decomposing orientations into the spherical harmonic (SH) coefficients is an effective tool for describing the angular distribution, and we can use SH coefficients $A_{lm}$ to describe the angular distribution $a(\hat{\mathbf{s}})$:

$$a(\hat{\mathbf{s}}) = \sum_{l=0}^{\infty} \sum_{m=-l}^{l} A_{lm} Y_{lm}(\hat{\mathbf{s}}) \tag{32}$$

where $Y_{lm}(\hat{\mathbf{s}})$ is the SH base function. In almost all kinds of PFM implementations, we think it is possible to perform such a conversion because of the angular band limit which truncates the SH coefficients to a small subset.

From Eq.(4), the ideal imaging process can be written as:

$$g_{\hat{\mathbf{p}}}(\mathbf{r}_d) = \sum_{lm} \int_{\mathbb{R}^3} \mathrm{d}\mathbf{r}_o \int_{\mathbb{S}^2} \mathrm{d}\hat{\mathbf{s}}_o \, H_{\hat{\mathbf{p}},lm}(\mathbf{r}_d - \mathbf{r}_o) F_{lm}(\mathbf{r}_o) \tag{33}$$

where we transfer the initial variables to angular-spectrum domain:

$$H_{\hat{\mathbf{p}},lm}(\mathbf{r}) = \int_{\mathbb{S}^2} \mathrm{d}\hat{\mathbf{s}}_o \, h_{\hat{\mathbf{p}}}(\mathbf{r}, \hat{\mathbf{s}}_o) Y_{lm}^*(\hat{\mathbf{s}}) \tag{34}$$

$$F_{lm}(\mathbf{r}_o) = \int_{\mathbb{S}^2} \mathrm{d}\hat{\mathbf{s}}_o \, f(\mathbf{r}, \hat{\mathbf{s}}_o) Y_{lm}^*(\hat{\mathbf{s}}) \tag{35}$$

2.1.2 Spherical harmonic multiplication and division

1) Multiplication

Let $a(\hat{\mathbf{s}})$ and $b(\hat{\mathbf{s}})$ be real-valued scalar functions defined on the unit sphere $\mathbb{S}^2$ with spherical harmonic expansions:

$$a(\hat{\mathbf{s}}) = \sum_{lm} A_{lm} Y_{lm}(\hat{\mathbf{s}}) \tag{36}$$

$$b(\hat{\mathbf{s}}) = \sum_{lm} B_{lm} Y_{lm}(\hat{\mathbf{s}}) \tag{37}$$

We define their pointwise product as:

$$c(\hat{\mathbf{s}}) = a(\hat{\mathbf{s}}) \cdot b(\hat{\mathbf{s}}) \tag{38}$$

which we would like to express in terms of expansion coefficients. First, we expand terms

$$c(\hat{\mathbf{s}}) = \left(\sum_{lm} A_{lm} Y_{lm}(\hat{\mathbf{s}})\right)\left(\sum_{l'm'} B_{l'm'} Y_{l'm'}(\hat{\mathbf{s}})\right) = \sum_{lm}\sum_{l'm'} A_{lm} B_{l'm'} Y_{lm}(\hat{\mathbf{s}}) Y_{l'm'}(\hat{\mathbf{s}}) \tag{39}$$

Next, we project $c(\hat{\mathbf{s}})$ onto the spherical harmonics by taking the inner product with $Y_{l''m''}(\hat{\mathbf{s}})$:

$$C_{l''m''} = \int_{\mathbb{S}^2} \mathrm{d}\hat{\mathbf{s}}\, c(\hat{\mathbf{s}})\, Y_{l''m''}(\hat{\mathbf{s}}) = \sum_{lm}\sum_{l'm'} A_{lm} B_{l'm'} \int_{\mathbb{S}^2} \mathrm{d}\hat{\mathbf{s}}\, Y_{lm}(\hat{\mathbf{s}}) Y_{l'm'}(\hat{\mathbf{s}}) Y_{l''m''}(\hat{\mathbf{s}}) \tag{40}$$

We simplify by defining the Gaunt coefficients as

$$G_{ll'l''}^{mm'm''} = \int_{\mathbb{S}^2} \mathrm{d}\hat{\mathbf{s}}\, Y_{lm}(\hat{\mathbf{s}}) Y_{l'm'}(\hat{\mathbf{s}}) Y_{l''m''}(\hat{\mathbf{s}}) \tag{41}$$

And substitute into Eq. (40)

$$C_{l''m''} = \sum_{lm}\sum_{l'm'} G_{ll'l''}^{mm'm''} A_{lm} B_{l'm'} \tag{42}$$

Sometimes it is valuable to rewrite this expression in matrix-vector form. If we define boldface vectors $\mathbf{A}$, $\mathbf{B}$, $\mathbf{C} \in \mathbb{R}^N$, where $N$ is the number of expansion coefficients in the product, we can rewrite Eq. (42) as

$$\mathbf{C} = \mathbf{M}(\mathbf{B}) \cdot \mathbf{A} \tag{43}$$

Where the matrix $\mathbf{M}(\mathbf{B}) \in \mathbb{R}^{N\times N}$ is

$$\mathbf{M}_{l''m''lm}(\mathbf{B}) = \sum_{l'm'} G_{ll'l''}^{mm'm''} B_{l'm'} \tag{44}$$

Note that multiplication commutes, i.e. the order does not matter, so Eq. (43) can be rewritten as

$$\mathbf{C} = \mathbf{M}(\mathbf{A}) \cdot \mathbf{B} \tag{45}$$

2) Division

Now consider computing the quotient of two spherical functions:

$$a(\hat{\mathbf{s}}) = \frac{c(\hat{\mathbf{s}})}{b(\hat{\mathbf{s}})} \tag{46}$$

Rearranging into a product $c(\hat{\mathbf{s}}) = a(\hat{\mathbf{s}}) \cdot b(\hat{\mathbf{s}})$ then applying the result from Eq. (43) yields:

$$\mathbf{C} = \mathbf{M}(\mathbf{B}) \cdot \mathbf{A} \tag{47}$$

If $\mathbf{M}(\mathbf{B})$ is invertible, the spherical harmonic coefficients of the quotient are given by

$$\mathbf{A} = [\mathbf{M}(\mathbf{B})]^{-1} \cdot \mathbf{C} \tag{48}$$

Note that division does not commute, i.e. the order matters, so we cannot swap the order of $\mathbf{B}$ and $\mathbf{C}$. Specifically,

$$\mathbf{A} \neq [\mathbf{M}(\mathbf{C})]^{-1} \cdot \mathbf{B} \tag{49}$$

2.1.3 Iterative process

To simplify the calculation and match the above dipole spatio-angular point spread function Eq. (33), we rewrite the operations of Eq. (9)-(13) in the SH domain. Let $\mathcal{F}_{\mathbb{S}^2}$ and $\mathcal{F}_{\mathbb{R}^3}$ denote the SH transform and Fourier transform, $\mathcal{F}_{\mathbb{S}^2}^{-1}$ and $\mathcal{F}_{\mathbb{R}^3}^{-1}$ the inverse of SH transform and Fourier transform. We will use the italic capital letters to represent the results of the Fourier or SH transform, and the block capitals letters to represent the results after both transforms, e.g., $E_{k,lm}$ is the result of $\mathcal{F}_{\mathbb{S}^2}\{e_k\}$ or $\mathcal{F}_{\mathbb{R}^3}\{e_k\}$ while $\mathrm{E}_{k,lm}$ represents the result of $\mathcal{F}_{\mathbb{S}^2}\mathcal{F}_{\mathbb{R}^3}\{e_k\}$.

First, the initial estimate is:

$$E_{0,lm}(\mathbf{r}) = \begin{cases} \sum_{\hat{\mathbf{p}}} i_{\hat{\mathbf{p}}}(\mathbf{r}), & l = 0, m = 0 \\ 0, & \text{others} \end{cases} \tag{50}$$

Then,

Step 1, forward projection in Eq. (9) changes to:

$$I'_{\hat{\mathbf{p}}}(\boldsymbol{\upsilon}) = \sum_{lm} \mathrm{H}_{\hat{\mathbf{p}},lm}(\boldsymbol{\upsilon}) \mathrm{E}_{k,lm}(\boldsymbol{\upsilon}) \tag{51}$$

Step 2, the forward correction ratio maintains the same as Eq. (10):

$$i''_{\hat{\mathbf{p}}}(\mathbf{r}_d) = \frac{i_{\hat{\mathbf{p}}}(\mathbf{r}_d)}{i'_{\hat{\mathbf{p}}}(\mathbf{r}_d)} \tag{52}$$

Step 3, the back-projecting update matrix in Eq. (11) changes to:

$$\mathrm{E}'_{k,lm}(\boldsymbol{\upsilon}) = \sum_{\hat{\mathbf{p}}} \mathrm{H}^{\mathrm{back}}_{\hat{\mathbf{p}},lm}(\boldsymbol{\upsilon}) I''_{\hat{\mathbf{p}}}(\boldsymbol{\upsilon}) \tag{53}$$

Step 4, the updated estimation in Eq. (12) can be changed to:

$$E''_{k,lm}(\mathbf{r}) = \sum_{l'm'} \sum_{l''m''} G_{ll'l''}^{mm'm''} E_{k,l'm'}(\mathbf{r}) E'_{k,l''m''}(\mathbf{r}) \tag{54}$$

in which, $G_{ll'l''}^{mm'm''}$ is the real Gaunt coefficient[4], and can be calculated by:

$$G_{ll'l''}^{mm'm''} = \int_{\mathbb{S}^2} \mathrm{d}\hat{\mathbf{s}}\, Y_{lm}(\hat{\mathbf{s}}) Y_{l'm'}(\hat{\mathbf{s}}) Y_{l''m''}(\hat{\mathbf{s}})\,. \tag{55}$$

Step 5, the latest estimation after sensitivity correction in Eq. (13) changes to:

$$E_{k+1,lm}(\mathbf{r}) = \sum_{l'm'} M_{lm,l'm'}^{-1}(\mathbf{r}) E_{k,l'm'}'' \,, \tag{56}$$

where:

$$M_{lm,l'm'}(\mathbf{r}) = \sum_{l''m''} G_{ll'l''}^{mm'm''} c_{l''m''}(\mathbf{r}) \,, \tag{57}$$

and $M_{lm,l'm'}^{-1}(\mathbf{r})$ means that, for each $\mathbf{r}$, we calculate the inverse of the 2D matrix $M_{lm,l'm'}$ build by dimension $lm$ and $l'm'$.

By fully transforming operations from the spherical domain to the SH domain, GRL can achieve a 30-and 15-fold reduction in terms of computation time and RAM requirement, respectively (**Figure SN2**).

**Algorithm S2: Generalized Richardson-Lucy in Spherical harmonics domain**

**Input**: $K$, $\mathrm{H}_{\hat{\mathbf{p}},lm}(\boldsymbol{\upsilon})$, $i_{\hat{\mathbf{p}}}(\mathbf{r})$

**Output**: $E_{K,lm}(\mathbf{r})$

$$E_{0,lm}(\mathbf{r}) = \begin{cases} \sum_{\hat{\mathbf{p}}} i_{\hat{\mathbf{p}}}(\mathbf{r}), & l = 0, m = 0 \\ 0, & \text{others} \end{cases}$$

$$c_{lm} = \sum_{\mathbf{p}} \int_{\mathbb{R}^3} \mathrm{d}\mathbf{r}\, h_{\hat{\mathbf{p}},lm}(\mathbf{r})$$

**For** $k = 0, 1, 2, \ldots, K-1$:

$$\mathrm{E}_{k,lm}(\boldsymbol{\upsilon}) = \mathcal{F}_{\mathbb{R}^3}\{E_{k,lm}(\mathbf{r})\}$$

$$I'_{\hat{\mathbf{p}}}(\boldsymbol{\upsilon}) = \sum_{lm} \mathrm{H}_{\hat{\mathbf{p}},lm}(\boldsymbol{\upsilon}) \mathrm{E}_{k,lm}(\boldsymbol{\upsilon})$$

$$i'_{\hat{\mathbf{p}}}(\mathbf{r}) = \mathcal{F}_{\mathbb{R}^3}^{-1}\{I'_{\hat{\mathbf{p}}}(\boldsymbol{\upsilon})\}$$

$$i''_{\hat{\mathbf{p}}}(\mathbf{r}) = \frac{i_{\hat{\mathbf{p}}}(\mathbf{r})}{i'_{\hat{\mathbf{p}}}(\mathbf{r})}$$

$$I''_{\hat{\mathbf{p}}}(\boldsymbol{\upsilon}) = \mathcal{F}_{\mathbb{R}^3}\{i''_{\hat{\mathbf{p}}}(\mathbf{r})\}$$

$$\mathrm{E}'_{k,lm}(\boldsymbol{\upsilon}) = \sum\nolimits_{\hat{\mathbf{p}}} \mathrm{H}^{\mathrm{back}}_{\hat{\mathbf{p}},lm}(\boldsymbol{\upsilon}) I''_{\hat{\mathbf{p}}}(\boldsymbol{\upsilon})$$

$$E'_{k,lm}(\mathbf{r}) = \mathcal{F}^{-1}_{\mathbb{R}^3}\{\mathrm{E}'_{k,lm}(\boldsymbol{\upsilon})\}$$

$$E''_{k,lm}(\mathbf{r}) = \sum\nolimits_{l'm'} \sum\nolimits_{l''m''} G^{mm'm''}_{ll'l''} E_{k,l'm'}(\mathbf{r}) E'_{k,l''m''}(\mathbf{r})$$

$$M_{lm,l'm'}(\mathbf{r}) = \sum\nolimits_{l''m''} G^{mm'm''}_{ll'l''} c_{l''m''}(\mathbf{r})$$

$$E_{k+1,lm}(\mathbf{r}) = \sum\nolimits_{l'm'} M^{-1}_{lm,l'm'}(\mathbf{r}) E''_{k,l'm'}$$

**End**

#### 2.2 Iterative framework modification

The traditional Richardson-Lucy algorithm has the following form:

$$e_{k+1} = e_k \times \frac{1}{c} \times \left(\frac{i}{e_k \star h} \star h^{\mathrm{back}}\right) \tag{58}$$

which involves a comparison of the measured projection data $i$ with calculated projections $e_k \star h$. When we consider extending the method to three or more dimensions, the arrays of measured and calculated projection data become excessively large. A modification was developed to handle the data without requiring such large arrays. This method, referred to as to the image space reconstruction algorithm[5] (ISRA), reverses the ordering of the comparison and back-projection steps by back-projecting the measured projection data ($i \star h^{\mathrm{back}}$, which can potentially be performed event-by-event in real time during data collection) and then uses this back-projected data image as the basis for comparison. The basic equation of this algorithm is given by:

$$e_{k+1} = e_k \times \left(\frac{i \star h^{\mathrm{back}}}{e_k \star h \star h^{\mathrm{back}}}\right) \tag{59}$$

and no longer requires the object space sensitive constant term $c$.

Regarding the generalized Richardson-Lucy algorithm in the spherical harmonics domain (Algorithm S2) in spatio-angular hyperspace, the arrays of measured and calculated projection data become excessively large due to the involving polarization channel in Eq. (51)-(53). We draw a direct analogy to GRL, and term this the efficient generalized Richardson-Lucy algorithm:

$$E_{k+1,lm}(\mathbf{r}) = E_{k,lm}(\mathbf{r}) \times_{\mathrm{Mul}} \frac{i_{\hat{\mathbf{p}}}(\mathbf{r}) \diamond H_{\hat{\mathbf{p}},lm}^{(\mathrm{back})}(\mathbf{r})}{E_{k,lm}(\mathbf{r}) \star H_{\hat{\mathbf{p}},lm}(\mathbf{r}) \diamond H_{\hat{\mathbf{p}},lm}^{(\mathrm{back})}(\mathbf{r})} \mathrm{Div} \tag{60}$$

which contains fewer steps comparing with Eq. (7) and these two terms, $i_{\hat{\mathbf{p}}}(\mathbf{r}) \diamond H_{\hat{\mathbf{p}},lm}^{(\mathrm{back})}(\mathbf{r})$ and $H_{\hat{\mathbf{p}},lm}(\mathbf{r}) \diamond H_{\hat{\mathbf{p}},lm}^{(\mathrm{back})}(\mathbf{r})$ can be calculated before iterations.

The forward projection $i_{\hat{\mathbf{p}}}(\mathbf{r}) \diamond H_{\hat{\mathbf{p}},lm}^{(\mathrm{back})}(\mathbf{r})$ is given by:

$$i_{\hat{\mathbf{p}}}(\mathbf{r}) \diamond H_{\hat{\mathbf{p}},lm}^{(\mathrm{back})}(\mathbf{r}) = \mathcal{F}_{\mathbb{R}^3}^{-1}\left\{\sum\nolimits_{\hat{\mathbf{p}}} \mathcal{F}_{\mathbb{R}^3}\left\{\mathrm{H}_{\hat{\mathbf{p}},l'm'}^{\mathrm{back}}(\mathbf{r})\right\}\mathcal{F}_{\mathbb{R}^3}\left\{i_{\hat{\mathbf{p}}}(\mathbf{r})\right\}\right\} \tag{61}$$

and the forward and backward projection $E_{k,lm}(\mathbf{r}) \star H_{\hat{\mathbf{p}},lm}(\mathbf{r}) \diamond H_{\hat{\mathbf{p}},lm}^{(\mathrm{back})}(\mathbf{r})$ is:

$$E'_{k,lm}(\mathbf{r}) = \mathcal{F}_{\mathbb{R}^3}^{-1}\left\{\sum\nolimits_{l'm'} \mathrm{H}_{lm,l'm'}^{\mathrm{comb}}(\boldsymbol{\upsilon})\mathcal{F}_{\mathbb{R}^3}\left\{\mathrm{E}_{k,l'm'}(\mathbf{r})\right\}\right\} \tag{62}$$

where the continuous forward and backward projectors can be combined into one operator:

$$\mathrm{H}_{lm,l'm'}^{\mathrm{comb}}(\boldsymbol{\upsilon}) = \sum\nolimits_{\hat{\mathbf{p}}} \mathcal{F}_{\mathbb{R}^3}\left\{H_{\hat{\mathbf{p}},lm}(\mathbf{r})\right\}\mathcal{F}_{\mathbb{R}^3}\left\{H_{\hat{\mathbf{p}},l'm'}^{\mathrm{back}}(\mathbf{r})\right\} \tag{63}$$

There are no more iterative operations involving the polarization $\hat{\mathbf{p}}$, reducing the computation especially when the number of polarization modulations is much more than that of SH coefficients (**Figure SN2**). Note that the vector of object-space sensitivity constants in Eq. (8) is no longer needed.

Here we give the backbone of our final eGRL algorithm:

**Algorithm S3: Efficient Generalized Richardson-Lucy algorithm**

Input: $K$, $\mathrm{H}_{\hat{\mathbf{p}},lm}(\boldsymbol{\upsilon})$, $\mathrm{H}_{\hat{\mathbf{p}},lm}^{\mathrm{back}}(\boldsymbol{\upsilon})$, $i_{\hat{\mathbf{p}}}(\mathbf{r})$

Output: $E_{K,lm}(\mathbf{r})$

$$E_{0,lm}(\mathbf{r}) = \begin{cases} \sum_{\hat{\mathbf{p}}} i_{\hat{\mathbf{p}}}(\mathbf{r}), & l = 0, m = 0 \\ 0, & \text{others} \end{cases}$$

$$I_{\hat{\mathbf{p}}}(\boldsymbol{\upsilon}) = \mathcal{F}_{\mathbb{R}^3}\left\{i_{\hat{\mathbf{p}}}(\mathbf{r})\right\}$$

$$\mathrm{E}_{lm}(\boldsymbol{\upsilon}) = \sum\nolimits_{\hat{\mathbf{p}}} \mathrm{H}_{\hat{\mathbf{p}},lm}^{\mathrm{back}}(\boldsymbol{\upsilon}) I_{\hat{\mathbf{p}}}(\boldsymbol{\upsilon})$$

$E_{lm}(\mathbf{r}) = \mathcal{F}_{\mathbb{R}^3}^{-1}\{\mathrm{E}_{lm}(\boldsymbol{\upsilon})\}$

$\mathrm{H}_{lm,l'm'}^{\mathrm{comb}}(\boldsymbol{\upsilon}) = \sum_{\hat{\mathbf{p}}} \mathrm{H}_{\hat{\mathbf{p}},lm}(\boldsymbol{\upsilon})\mathrm{H}_{\hat{\mathbf{p}},l'm'}^{\mathrm{back}}(\boldsymbol{\upsilon})$

For $k = 0, 1, 2, \dots, K-1$:

$\mathrm{E}_{k,lm}(\boldsymbol{\upsilon}) = \mathcal{F}_{\mathbb{R}^3}\{E_{k,lm}(\mathbf{r})\}$

$\mathrm{E}'_{k,lm}(\boldsymbol{\upsilon}) = \sum_{l'm'} \mathrm{H}_{lm,l'm'}^{\mathrm{comb}}(\boldsymbol{\upsilon})\mathrm{E}_{k,l'm'}(\boldsymbol{\upsilon})$

$E'_{k,lm}(\mathbf{r}) = \mathcal{F}_{\mathbb{R}^3}^{-1}\{\mathrm{E}'_{k,lm}(\boldsymbol{\upsilon})\}$

$M_{lm,l'm'}(\mathbf{r}) = \sum_{l''m''} G_{ll'l''}^{mm'm''} E'_{k,l''m''}(\mathbf{r})$

$E''_{k,lm}(\mathbf{r}) = \sum_{l'm'} M_{lm,l'm'}^{-1}(\mathbf{r})E_{l'm'}(\mathbf{r})$

$E_{k+1,lm}(\mathbf{r}) = \sum_{l'm'} \sum_{l''m''} G_{ll'l''}^{mm'm''} E_{k,l'm'}(\mathbf{r})E''_{k,l''m''}(\mathbf{r})$

End

---

and its additive form for dual-view microscopy like pol-diSPIM, i.e., spatio-angular deconvolution of polarization images in each view separately and then perform the additive fusion:

**Algorithm S4: Efficient Generalized Richardson-Lucy for pol-diSPIM (the additive form)**

---

Input: $K$, $\mathrm{H}_{\hat{\mathbf{p}},lm}^{(v1)}(\boldsymbol{\upsilon})$, $\mathrm{H}_{\hat{\mathbf{p}},lm}^{(v2)}(\boldsymbol{\upsilon})$, $\mathrm{H}_{\hat{\mathbf{p}},lm}^{\mathrm{back}(v1)}(\boldsymbol{\upsilon})$, $\mathrm{H}_{\hat{\mathbf{p}},lm}^{\mathrm{back}(v2)}(\boldsymbol{\upsilon})$, $i_{\hat{\mathbf{p}}}^{(v1)}(\mathbf{r})$, $i_{\hat{\mathbf{p}}}^{(v2)}(\mathbf{r})$

Output: $E_{K,lm}(\mathbf{r})$

---

$$E_{0,lm}(\mathbf{r}) = \begin{cases} \sum_{\hat{\mathbf{p}}} i_{\hat{\mathbf{p}}}^{(v1)}(\mathbf{r}) + \sum_{\hat{\mathbf{p}}} i_{\hat{\mathbf{p}}}^{(v2)}(\mathbf{r}), & l = 0, m = 0 \\ 0, & \text{others} \end{cases}$$

$I_{\hat{\mathbf{p}}}^{(v1)}(\boldsymbol{\upsilon}) = \mathcal{F}_{\mathbb{R}^3}\{i_{\hat{\mathbf{p}}}^{(v1)}(\mathbf{r})\}$

$I_{\hat{\mathbf{p}}}^{(v2)}(\boldsymbol{\upsilon}) = \mathcal{F}_{\mathbb{R}^3}\{i_{\hat{\mathbf{p}}}^{(v2)}(\mathbf{r})\}$

$\mathrm{E}_{lm}^{(v1)}(\boldsymbol{\upsilon}) = \sum_{\hat{\mathbf{p}}} \mathrm{H}_{\hat{\mathbf{p}},lm}^{\mathrm{back}(v1)}(\boldsymbol{\upsilon})I_{\hat{\mathbf{p}}}^{(v1)}(\boldsymbol{\upsilon})$

$E_{lm}^{(v1)}(\mathbf{r}) = \mathcal{F}_{\mathbb{R}^3}^{-1}\{\mathrm{E}_{lm}^{(v1)}(\boldsymbol{\upsilon})\}$

$$\mathrm{E}_{lm}^{(v2)}(\boldsymbol{\upsilon}) = \sum_{\hat{\mathbf{p}}} \mathrm{H}_{\hat{\mathbf{p}},lm}^{\mathrm{back}(v2)}(\boldsymbol{\upsilon}) I_{\hat{\mathbf{p}}}^{(v2)}(\boldsymbol{\upsilon})$$

$$E_{lm}^{(v2)}(\mathbf{r}) = \mathcal{F}_{\mathbb{R}^3}^{-1}\left\{\mathrm{E}_{lm}^{(v2)}(\boldsymbol{\upsilon})\right\}$$

$$\mathrm{H}_{lm,l'm'}^{\mathrm{comb}(v1)}(\boldsymbol{\upsilon}) = \sum_{\hat{\mathbf{p}}} \mathrm{H}_{\hat{\mathbf{p}},lm}^{(v1)}(\boldsymbol{\upsilon}) \mathrm{H}_{\hat{\mathbf{p}},l'm'}^{\mathrm{back}(v1)}(\boldsymbol{\upsilon})$$

$$\mathrm{H}_{lm,l'm'}^{\mathrm{comb}(v2)}(\boldsymbol{\upsilon}) = \sum_{\hat{\mathbf{p}}} \mathrm{H}_{\hat{\mathbf{p}},lm}^{(v2)}(\boldsymbol{\upsilon}) \mathrm{H}_{\hat{\mathbf{p}},l'm'}^{\mathrm{back}(v2)}(\boldsymbol{\upsilon})$$

For $k = 0, 1, 2, \dots, K-1$:

$$\mathrm{E}_{k,lm}(\boldsymbol{\upsilon}) = \mathcal{F}_{\mathbb{R}^3}\left\{E_{k,lm}(\mathbf{r})\right\}$$

View 1

$$\mathrm{E}_{k,lm}^{\prime(v1)}(\boldsymbol{\upsilon}) = \sum_{l'm'} \mathrm{H}_{lm,l'm'}^{\mathrm{comb}(v1)}(\boldsymbol{\upsilon}) \mathrm{E}_{k,l'm'}^{(v1)}(\boldsymbol{\upsilon})$$

$$E_{k,lm}^{\prime(v1)}(\mathbf{r}) = \mathcal{F}_{\mathbb{R}^3}^{-1}\left\{\mathrm{E}_{k,lm}^{\prime(v1)}(\boldsymbol{\upsilon})\right\}$$

$$M_{lm,l'm'}^{(v1)}(\mathbf{r}) = \sum_{l''m''} G_{ll'l''}^{mm'm''} E_{k,l''m''}^{\prime(v1)}(\mathbf{r})$$

$$E_{k,lm}^{\prime\prime(v1)}(\mathbf{r}) = \sum_{l'm'} M_{lm,l'm'}^{-1(v1)}(\mathbf{r}) E_{l'm'}^{(v1)}(\mathbf{r})$$

$$E_{k+1,lm}^{(v1)}(\mathbf{r}) = \sum_{l'm'} \sum_{l''m''} G_{ll'l''}^{mm'm''} E_{k,l'm'}^{(v1)}(\mathbf{r}) E_{k,l''m''}^{\prime\prime(v1)}(\mathbf{r})$$

View 2

$$\mathrm{E}_{k,lm}^{\prime(v2)}(\boldsymbol{\upsilon}) = \sum_{l'm'} \mathrm{H}_{lm,l'm'}^{\mathrm{comb}(v2)}(\boldsymbol{\upsilon}) \mathrm{E}_{k,l'm'}^{(v2)}(\boldsymbol{\upsilon})$$

$$E_{k,lm}^{\prime(v2)}(\mathbf{r}) = \mathcal{F}_{\mathbb{R}^3}^{-1}\left\{\mathrm{E}_{k,lm}^{\prime(v2)}(\boldsymbol{\upsilon})\right\}$$

$$M_{lm,l'm'}^{(v2)}(\mathbf{r}) = \sum_{l''m''} G_{ll'l''}^{mm'm''} E_{k,l''m''}^{\prime(v2)}(\mathbf{r})$$

$$E_{k,lm}^{\prime\prime(v2)}(\mathbf{r}) = \sum_{l'm'} M_{lm,l'm'}^{-1(v2)}(\mathbf{r}) E_{l'm'}^{(v2)}(\mathbf{r})$$

$$E_{k+1,lm}^{(v2)}(\mathbf{r}) = \sum_{l'm'} \sum_{l''m''} G_{ll'l''}^{mm'm''} E_{k,l'm'}^{(v2)}(\mathbf{r}) E_{k,l''m''}^{\prime\prime(v2)}(\mathbf{r})$$

$$E_{k+1,lm}(\mathbf{r}) = \frac{1}{2}\left(E_{k+1,lm}^{(v1)}(\mathbf{r}) + E_{k+1,lm}^{(v2)}(\mathbf{r})\right)$$

End

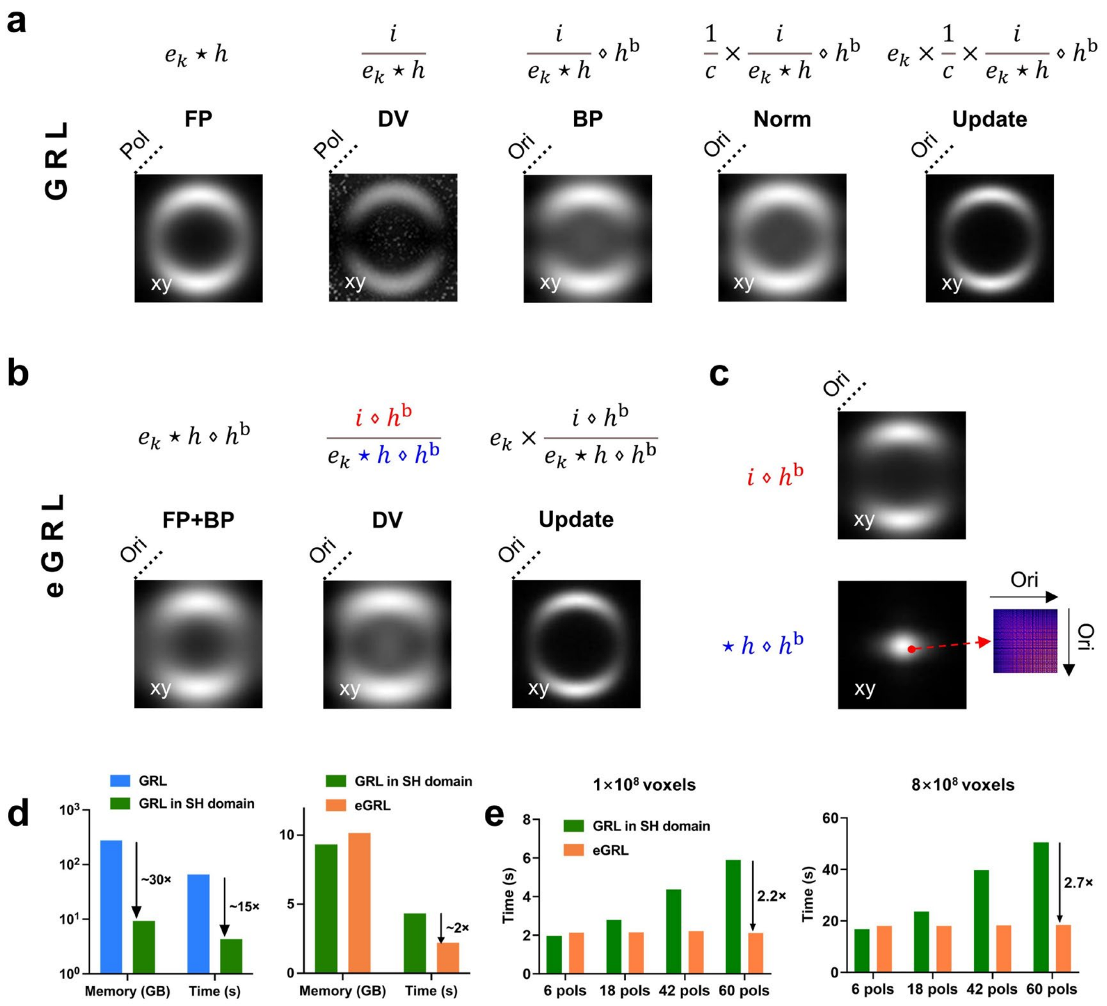

**Figure SN2, GRL and eGRL for hyperspace spatio-angular image reconstruction frameworks. a)** The structure of GRL for each iterative step includes FP (forward projection), DV (division), BP (back projection), Norm (normalization by object space sensitivity constant) and Update, with a similar form as the traditional Richardson-Lucy algorithm, but extending to hyperspace with additional polarization (Pol) or orientation (Ori) dimensions. **b), c)** Framework of eGRL variant of GRL. Note only the three steps in **b** need to be calculated in each iteration and the other steps in **c** can be precomputed, resulting in a faster runtime than GRL. **d)** Time and memory cost of GRL, GRL in SH domain (after angular domain transformation) and final eGRL reconstruction in a dataset with 100×100×100 voxels and 42 polarization modulations. Left: the reduction of time and memory cost by transferring the computation from spherical domain to SH domain; Right: comparison of time and memory cost between GRL and eGRL iteration structure, both in SH domain. Note eGRL in the SH domain offers a 30-fold improvement in both time and memory cost over the native GRL implementation in the spherical domain. **e)** Bar graphs of the time required to process the datasets with different number of voxels and polarization modulations (pols) by GRL and the variant eGRL, showing the eGRL is more efficient than the GRL.
